# From Sensory Detection to Motor Action: The Comprehensive *Drosophila* Taste-Feeding Connectome

**DOI:** 10.1101/2025.08.25.671814

**Authors:** Ibrahim Tastekin, Inês de Haan Vicente, Rory J. Beresford, Billy J. Morris, Isabella Beckett, Philipp Schlegel, Marina Gkantia, FlyEM Project Team, Cambridge Connectomics Group, Elizabeth C. Marin, Marta Costa, Gregory S.X.E. Jefferis, Carlos Ribeiro

## Abstract

Gustatory systems drive critical survival behaviors such as feeding, foraging, and social interactions. However, gustation remains one of the least mapped sensory modalities at the connectome level. Here, we present the first complete wiring diagram of the male *Drosophila* adult gustatory system, comprehensively reconstructing gustatory receptor neurons (GRNs) from peripheral organs in a contiguous electron microscopy volume spanning brain, cervical connective, and ventral nerve cord. Integrating this with existing datasets, we generated a pan-CNS, cross-sex connectome that reveals GRN diversity through connectivity-based clustering, molecular identity mapping, and sexual dimorphism analysis. We mapped all feeding motor neurons and traced complete sensory-to-motor pathways to feeding, foraging, endocrine, and social behavior circuits. The emerging circuit architectures reveal distinct circuits for nutrient assessment, motor control, neuroendocrine regulation, and courtship. This work defines the gustatory system’s organization at synaptic resolution and provides a framework for understanding how internal states modulate sensory-driven decisions across behavioral contexts.

## Introduction

Taste is an ancient chemosensory modality that every major animal lineage, from nematodes to insects, fish, and humans, uses to turn the chemical composition of the environment into immediate, value-based actions. By classifying molecules as attractive, neutral, or noxious, gustatory circuits couple detection directly to hard-wired behaviors such as ingestion, rejection, escape, or courtship, thereby steering core survival and fitness decisions in feeding and mating ^1,2^. It also conveys important value information which can be used by learning and memory systems to reweigh behavior and guide future choices ^3,4^. Importantly, gustatory circuits are continuously modulated by internal-state signals, including hunger, satiety, reproductive status, and competing behaviors, such that the very same taste cue can prompt drastically different behaviors depending on the animal’s physiological needs ^5^.

*Drosophila melanogaster* offers a uniquely tractable system for dissecting both sensorimotor transformations as well as complex behaviors, such as foraging, and how these are modulated by internal states. Its gustatory receptor neurons (GRNs) are molecularly typed, morphologically and behaviorally stereotyped ^1,6^, and genetically accessible ^7–9^, yet are embedded in a brain small enough to be reconstructed completely with recent advances in electron microscopy (EM) connectomics, while still allowing for complex and sophisticated behaviors. Gustatory pathways are also among the most sexually dimorphic sensory systems in the fly ^10,11^. Several GRN types and their brain targets differ between males and females to bias courtship and sex specific behaviors such as oviposition ^4,12^. But a comprehensive description of the gustatory system and its downstream circuits is still lacking. Importantly, in the fly, the recently developed connectome maps ^13,14^, combined with neurogenetic tools to monitor and manipulate unique neuronal cell types, make it possible to link single identified neurons to feeding, aversion, locomotor, or foraging programs at cellular resolution.

In insects, the impact of taste scales from individual fitness to ecosystems with a significant impact on human wellbeing ^15–17^. Adult female mosquitoes, for example, rely on stylet and labellar gustatory neurons to distinguish nectar from blood and to evaluate human skin chemistry, making taste a primary gateway for pathogen transmission ^15^. Likewise, the tarsal and palpal sensilla of herbivorous or pollinating insects assess phagostimulants and deterrents on leaves and flowers, determining both crop damage and pollination, which is essential for food security ^17,18^. Deciphering the neural logic of gustation in insects thus not only provides a tractable model for understanding gustation at a molecular and circuit level but also eventually manipulating the feeding and host-choice behaviors that underpin agriculture and the spreading of vector-borne diseases.

In flies, gustatory receptor neurons are not confined to the mouthpart (labellum), but reside in sensilla distributed across different body parts: three groups of pharyngeal sensilla, tarsal and tibial bristles on all six legs, and rows of chemosensory hairs along the wing margin ^19,20^. Axons from these anatomically dispersed sensory neurons travel via discrete nerves yet converge onto partially overlapping territories of the sub-esophageal zone (SEZ) and ventral nerve cord (VNC), integrating external sampling, postingestive and internal state feedback, and reproductive cues within a common circuit framework. At the motor end of this circuit, proboscis muscle motor neurons have been catalogued at single-cell resolution, and SEZ pathways that arrest locomotion upon appetitive taste have been delineated, providing defined behavioral outputs onto which the incoming gustatory tracts must map ^21–26^. This distributed architecture broadens the behavioral contexts in which taste can act—during walking, feeding, and mating, for example, and highlights the importance of whole central nervous system reconstructions for a complete understanding of gustatory processing.

EM connectomics has already delivered complete wiring diagrams for other *Drosophila* senses: a full inventory of the visual system allows for circuit-level models of motion and color vision ^27^, while full-brain reconstructions of olfactory and mechanosensory pathways reveal canonical motifs of sensory convergence and descending motor control ^28,29^. These resources include the complete adult female brain connectome (FAFB – FlyWire) ^14,30,31^ and the male ventral-nerve-cord connectome (MANC) ^32–34^, which together chart brain-to-VNC connectivity at synaptic resolution. In contrast, gustation remains one of the least mapped modalities at the EM connectome level. Only subsets of labellar bristle, wing bristle, and leg bristle GRNs have been annotated ^33,35^, and the long-range tracts that relay taste to premotor, learning, and foraging centers, and their potential sex-specific differences, have yet to be mapped at synaptic resolution in a single, contiguous dataset.

Here, we present a pan-central nervous system (CNS) gustatory connectome derived from a contiguous adult male serial-section EM volume that spans the brain, cervical connective, and ventral nerve cord. We traced and annotated every gustatory receptor neuron from the labellum, pharyngeal sensilla, legs, and anterior wing margins and registered them to light-level receptor maps. By cross-matching homologous neuronal cell types from the female brain (FAFB – FlyWire) and the male ventral nerve cord (MANC), we expand this analysis to integrate all available EM datasets, in order to assemble a comprehensive and sex-comparable wiring diagram of the fly gustatory system, allowing us to explore sexual dimorphisms in the gustatory system. We then clustered these neurons by fine-scale morphology and by the identity of their downstream synaptic partners, revealing distinct sensory-functional classes that we propose correspond to appetitive, aversive, and pheromonal pathways. The classes converge onto compact second-order hubs that broadcast to proboscis-motor neurons, descending-premotor neurons, locomotor-stop circuits, and neuroendocrine cells, completing the link from detection to defined motor and physiological outputs. This composite dataset closes a major sensory gap in *Drosophila* connectomics and provides a reference for cross-modal computation, targeted perturbation, and ecological applications.

## Results

### Characterization of the fly’s gustatory system at the light level

Since gustatory receptor neurons (GRNs) play a crucial role in *Drosophila melanogaster*’s perception of its chemical environment, these neurons were selected as our entry point to start defining the fly’s gustatory connectome. GRNs are housed in sensory organs, named taste sensilla ^1^. These structures are distributed through multiple parts of the fly’s body, and their morphology differs depending on their exact location ^10,20,36–39^ (Figure 1A). Although very stereotypical in terms of numbers across flies, sexual dimorphisms exist in specific taste sensilla numbers ^10,36^. To map each of the different types of taste sensilla and their location across the body, we used different genetic strategies combined with confocal microscopy to label and systematically characterize all GRNs (Figure 1A and 1B).

**Figure 1.**
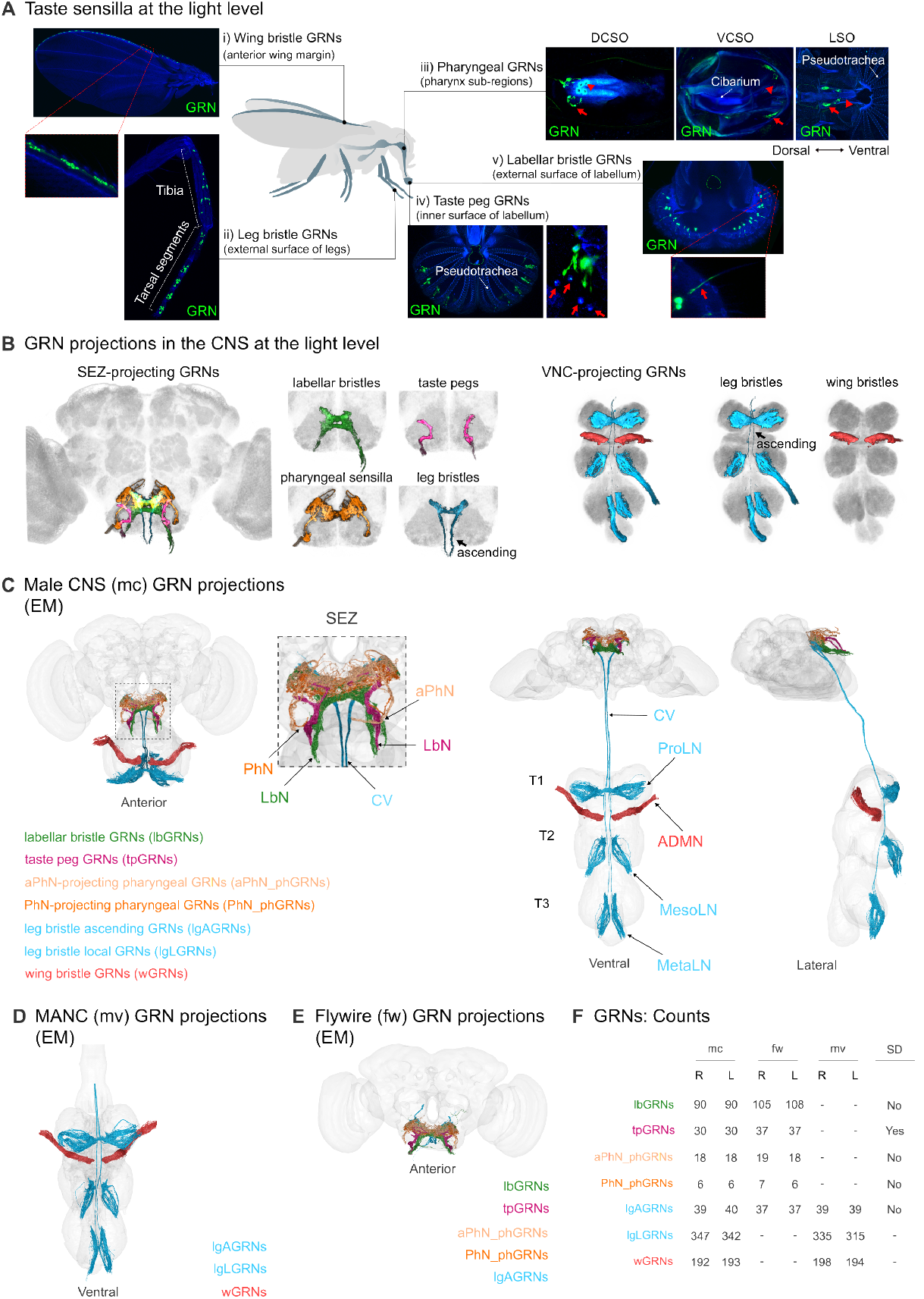
Characterization of the *Drosophila melanogaster* gustatory system. **A**. Confocal images using the membrane-tethered UAS-mCD8:GFP under the control of *poxn-GAL4* to label GRNs housed in taste bristles in the anterior margin of the wings **(i)**, on both the tarsi and tibia of all six legs **(ii)**, and on the external surface of the labellum **(v)**. The membrane-tethered UAS-mCD8:GFP was also used under the control of *R57F03-* and *Ir25a-GAL4* to label GRNs housed, respectively, in two other taste sensilla types: taste pegs located between pseudotrachea on the inner surface of the labellum **(ii)**; taste pores, marked with an arrowhead, across the pharynx **(iv)**, specifically, in the labral sensory organ (LSO), ventral cibarial sensory organ (VCSO), and dorsal cibarial sensory organ (DCSO). In **i)** the anterior wing margin is zoomed in to show the GRNs in the anterior wing margin in more detail. In **ii)** the tarsal segments and tibia are indicated. Arrows indicate the GRN soma in **iii)**. Cibarium and pseudotrachea are indicated. In **iv)** the dashed circle indicates the entry to the pharynx, and the pseudotrachea are indicated. The arrows indicate the taste peg structure (right panel). In **v)** the dashed circle indicates the entry to the pharynx. The arrow indicates the dendrites of the labellar bristle GRNs (bottom panel). **B**. Expression of membrane-tethered mCD8:GFP under the control of either the *Ir25a-* or the *poxn-GAL4* driver was used to visualize all GRN subclasses projecting, respectively, to the SEZ in the brain (left) or the VNC (right). The GRN axonal projections of each taste sensilla type are shown in a different color: labellar bristles in green; taste pegs in pink; pharyngeal sensilla in orange; ascending and local leg bristles in blue, and wing bristles in red. **C**. Rendering of all neurons identified per GRN subclass in the male CNS in three different views, anterior (left), ventral, and lateral (right). Identification of the GRNs in the connectome was achieved by defining seed planes across their specific entry nerve: labial nerve (LbN) for labellar bristle (green) and taste peg (pink) GRNs; accessory pharyngeal nerve (aPhN) for LSO and VCSO pharyngeal GRNs (light orange); pharyngeal nerve (PhN) for DCSO pharyngeal GRNs (dark orange); prothoracic (ProLN), mesothoracic (MesoLN), metathoracic (MetaLN) leg nerves, respectively, for fore-, mid- and hindleg bristle GRNs (blue); anterior dorsal mesothoracic nerve (ADMN) for wing bristle GRNs. Leg bristle GRNs are subdivided into ascending, which project through the cervical connective (CV) and arborize in the SEZ, and the local that terminate in the VNC. **D**. Ventral view of the renderings of the GRN subclasses – leg bristle ascending, leg bristle local, and wing bristle GRNs – identified in MANC. **E**. Anterior view of the renderings of the GRN subclasses – labellar bristle, taste peg, pharyngeal and leg bristle ascending GRNs – identified in FAFB – FlyWire. **F**. GRN counts per subclass across the right (R) and left (L) sides for male CNS, FAFB – FlyWire and MANC. A sexual dimorphism (SD) is found solely for the taste peg GRN counts. The difference in the counts of labellar bristle GRNs between male CNS and FAFB – FlyWire has been linked to automated segmentation errors.

Making direct contact with the external chemical environment, taste bristles are the most numerous and widely dispersed type of taste sensilla. They are thin and long hair-like structures with a pore at their tip, and their numbers vary across body parts, as does the number of GRNs they are innervated by ^10,36,37^. We expressed a membrane marker (mCD8:GFP) using *poxn-GAL4-14*, a driver which had been previously shown to drive expression in all taste bristles ^40^ (Figure 1A). As described previously, in both sexes, taste bristles are found on the tarsi and tibia of the flies’ six legs, on the anterior margin of the wings, and on the external surface of the labellum (located on the proboscis) (Figure 1Ai,1Aii and 1Av). In general, 30 taste bristles are found on each fly leg (Figure 1Aii), each housing two to four GRNs ^6,20,36^. Male forelegs are an exception, containing, generally, 50 taste bristles ^11,36^. Leg bristle GRNs (lgGRNs) are thought to play a key role in sampling the food in the environment during foraging and suppressing locomotion upon an appetitive response ^41^. The additional bristles on the male forelegs are important for sensing contact pheromones, crucial for courtship behavior ^42–44^. On the anterior margin of each male or female wing, 40 taste bristles have been reported, each innervated by four GRNs ^37^ (Figure 1Ai). Wing bristle GRNs (wGRNs) have been proposed to play a role in the exploration of ecological niches, especially in the context of sexual behavior ^45,46^. Each labial palp of both males and females is reported to be covered by approximately 31 bristles ^6,10,20^ (Figure 1Av). Labellar bristles are reported to vary in length and number of GRNs housed depending on their specific localization on the labellum ^10^. On the labellum’s periphery closest to where the two labial palps meet, there are short(S)- and long(L)-type bristles, which house four GRNs each. Occupying a more dorsal area of the labellum’s external surface, the intermediate(I)-type bristles are innervated by only two GRNs. Labellar bristle GRNs (lbGRNs) are thought to play a key role in mediating the decision to initiate feeding ^19,47,48^. When flies open their labellum, an event known as labellar abduction or spreading, food becomes accessible to pit-like sensilla, named taste pegs. These anatomically distinct taste sensilla are organized in rows between the pseudotrachea, a distinctive structure on the labellum’s inner surface ^10^, as shown using the *57F03-GAL4* driver, which we have previously demonstrated to label taste peg GRNs (tpGRNs) ^48^ (Figure 1Aiv). Each taste peg houses a single GRN, and their number is sexually dimorphic, with females having 20% more of these sensilla than males ^10^ (Figure 1F). In an accompanying paper, we show that a subset of tpGRNs play an important role in controlling the length of a feeding burst towards yeast, by regulating the efficiency of ingestion ^49^.

Upon ingestion, taste pores lining the pharynx function as internal gustatory sensors. As all GRNs in the pharynx have been reported to express IR25a ^39^, a broadly expressed gustatory co-receptor ^50,51^, we used *Ir25a-GAL4* to characterize the three specific pharyngeal subregions where these hairless sensilla reside (Figure 1Aiii). The labral sensory organ (LSO), which is composed of nine sensilla, conventionally numbered 1-9, is located ventrally on each side of the midline of the proboscis’ haustellum ^36,50^. Of those sensilla, only #7-9 house GRNs, with sensillum #7 being innervated by eight GRNs and sensilla #8 and #9 by one GRN each. The ventral cibarial sensory organ (VCSO) and the dorsal cibarial sensory organ (DCSO) are located within the proboscis’ rostrum, on each side of the pharynx ^39,52^. The VCSO has been reported to comprise either two or three sensilla, housing a total of eight GRNs per side, while the DCSO includes two sensilla per side with three GRNs each. The pharyngeal GRNs (phGRNs) are thought to act as gatekeepers, providing feedback about the quality of the food being ingested and, thus, impacting the decision of accepting and continuing to feed or rejecting it ^53,54^.

Besides GRNs, taste sensilla often also house one mechanosensory neuron (MSN) ^20^. Although being a highly relevant component of the fly’s feeding system ^55,56^, we have not included MSNs in the hereby presented connectome analysis.

### Description of the GRN projections in the CNS

According to the type of taste sensilla that house them and their localization on the fly body, we classified the GRNs into five broad subclasses: labellar bristle, taste peg, pharyngeal, leg bristle, and wing bristle GRNs (Figure 1A). Being bipolar sensory neurons, each GRN has a cell body that is located in the periphery, a single dendrite extending towards a pore at the sensillum extremity, through which tastants and other chemicals are sensed, and an axon that projects to specific neuropils in the CNS ^52,57^. Because these neurons either terminate and arborize in the SEZ or in the VNC, we grouped the GRN subclasses into SEZ-projecting and VNC-projecting GRNs (Figure 1B). To match the projections from the different body parts to the EM data, we first mapped the projection pattern in the CNS of each broad GRN sub-class at the light level, followed by manual segmentation of the different projection zones. For this, we used specific GRN-labelling lines for each GRN group. We used *Ir25a-GAL*4 driving UAS-mCD8:GFP to image the SEZ-projecting GRNs, which include all GRN subclasses, except the wGRNs and a subset of lgGRNs which project to the VNC (Figure 1B). In the SEZ, each GRN subclass projects to discrete regions in the neuropil ^57^. phGRNs enter most anteriorly in the SEZ and terminate in regions, including the ventral pharyngeal sensory center 1 (VPS1) and 2 (VPS2), and the posterior maxillary sensory zone 1 (PMS1), which is shared with lbGRNs that also cover other PMS zones ^57^. tpGRNs innervate the anterior maxillary sensory zone 1 (AMS1) ^57^. Contrary to the other subclasses, lgGRNs either project directly to the SEZ in the brain via the cervical connective, which we named leg ascending GRNs (lgAGRNs), or terminate in the ipsilateral VNC leg neuropils in the first (T1), second (T2), and third (T3) thoracic neuromeres, and we therefore called local leg GRNs (lgLGRNs) ^58^ (Figure 1B and 1C). The lgA-GRNs enter the gnathal ganglia (GNG) in the SEZ posteriorly and innervate a region corresponding to the anterior cerebrocervical fascicle (aCCF) terminals, known as aCCFG1^57^. These neurons have very few connections in the VNC. The lgLGRNs are mostly ipsilateral. However, the additional male-specific taste bristles on the male forelegs house bilateral neurons that cross the midline and innervate the leg neuropil on the contralateral side ^11^ (Figure 1B and 1C). The wGRNs terminate in a tightly packed ipsilateral projection in the ovoid, a specialized wing, and notum sensory neuropil on each side within T2^37^ (Figure 1B and 1C).

We also used this light level information together with existing reports ^20,36,59^ to identify the nerves by which the different GRN classes enter the CNS (Figure 1C). lbGRNs and tpGRNs project through the labial nerve (LbN) from the labellum to the brain. The phGRNs project to the brain via two other mouthpart nerves, the accessory pharyngeal nerve (aPhN), which includes the LSO and VCSO neurons, and the pharyngeal nerve (PhN) that contains the DCSO neurons. The wGRNs enter the VNC through the anterior dorsal mesothoracic nerve (ADMN) and the lgGRNs via the prothoracic (ProLN), mesothoracic (MesoLN), and metathoracic (MetaLN) nerves, depending if they innervate taste bristles on the fore-, mid-, or hindlegs, respectively.

### Identification of the GRNs across connectomes

Previous connectomes have covered only part of the CNS, either the brain or the VNC, in isolation. As a result, the gustatory system remains one of the last major sensory systems that have not been comprehensively characterized at the EM level. To close this gap, we focused our annotation and characterization effort on a recently generated male CNS volume, in which the brain and VNC are connected through the neck connective ^60^. This dataset gives us complete access to all GRN subclasses across different body parts within one volume (Figure 1C). Moreover, a substantial fraction of the GRNs’ direct downstream partners comprises descending (DNs) and ascending (ANs) neurons, whose complete form and connectivity are captured in this dataset. To find the GRNs in this volume, we used the available knowledge about the nerves through which each GRN subclass enters the CNS (Figure 1C). Neurons were identified as GRNs if their projection field and morphology corresponded to those previously identified in the literature and in our light level maps. We furthermore systematically proofread all the relevant neurons to ensure that the subsequent cell typing was of the highest possible quality. In addition, to avoid missing neurons due to cell segmentation errors, we also employed a complementary approach in which, for each subclass, we identified all the input neurons to the second-order partners of the identified GRNs and reconstructed unidentified upstream bodies that could potentially be poorly reconstructed GRNs, resulting in a comprehensive reconstruction of almost all the GRNs in the male CNS EM volume (Figure 1C). One of the major limitations of connectomics analysis is the small sample size. Thus, we extended our analysis to two publicly available datasets, the MANC ^32,33^ and the FAFB – FlyWire ^14,30,31^ connectomes, respectively, a male VNC and a female brain (Figures 1D and 1E). We completed the existing annotations of every GRN subclass in each volume.

Left-right and cross-dataset counts align closely with one another and with the numbers reported in the literature (Figure 1F). The few discrepancies fall into three clear categories: (i) subclasses with recognized sexual dimorphism, such as the 20% excess tpGRNs in females ^10^; (ii) subclasses prone to segmentation artefacts, notably lbGRNs whose thin, densely bundled axons are sometimes falsely split or merged; and (iii) regions of poorer tissue preservation in MANC, specifically of the leg nerves, which likely under-represent lgLGRNs ^33^. Based on these data, as well as our light level analysis, we propose that, except for the tpGRNs, sexual dimorphisms observed in the connectome counts are likely to result from technical artefacts. Nevertheless, aside from predictable sexual dimorphisms and these minor segmentation artefacts, all subclass counts differ by only two to three neurons per side across datasets (Figure 1F). One notable exception to the counts in the literature is the wing bristle population: we consistently recover 192-198 wGRNs per side, about 18% more than the long-standing estimate in the literature (∼160 neurons) ^20,37^. This upward revision suggests a denser gustatory coverage on the wing than previously appreciated.

Overall, we generated a near-complete GRN dataset across three different connectomes, allowing us to systematically cell type and characterize their downstream circuits. This provides us with the unique opportunity to generate functional hypotheses of how these circuits might initiate, terminate, or sustain foraging, feeding, and other gustatory-driven behaviors.

### Cell typing the labellar bristle GRNs

A neuronal cell type can, generally, be defined as a group of neurons that is more similar to neurons in another brain than to any other neurons in the same brain ^31^. Our approach for defining cell types was based on clustering our neurons of interest based on their synaptic connectivity, meaning the neurons they directly connect to, and validating the identified clusters through detailed anatomical inspection and comparison, combined with projection field-based matching. However, to be able to type the GRNs, especially in the male CNS that was originally mostly untyped, and then, co-cluster the neurons across connectomes, we also had to type their synaptic partners. This was an iterative two-step process that involved grouping the GRN partners using connectivity clustering and assigning a type, based on morphologically matching them to neurons in a more comprehensively typed connectome, like FAFB – Fly-Wire ^31,61^. Whenever necessary, the GRN partners were also proofread.

We started by typing lbGRNs as these are the best characterized gustatory neuron subclass in *Drosophila* ^12,35,62–65^. We decided to leverage our typing of GRNs in multiple datasets to provide extra robustness to our analysis. Therefore, we used both the lbGRNs we identified in the male CNS (Figures 2A and S1A) and, simultaneously, the corresponding ones in FAFB – FlyWire (Figure 2B and 2C), allowing us to both increase our sampling of neurons by including data from different brains as well as identifying potential sexual dimorphisms. We first clustered male and female lbGRNs separately using cosine similarity scores based on their output connectivity (Figure S1B and S1C), followed by a co-clustering step using both datasets (Figure 2D). Our analysis revealed four broad cell types for the lbGRNs, which we named LB1-4, based on and expanding on a previous cell typing of these neurons using the FAFB – FlyWire dataset ^31,35,66,67^. The clustering results are supported by clear differences in the overall morphologies of the four types and their consequently distinct projection fields in the SEZ (Figure 2E). Importantly, these features were identical across both connectomes, further strengthening our typing results (Figure S1D and S1E). Upon a complementary detailed inspection of the top downstream partners and the GRN morphologies within each cell type, we further divided LB1-4 into several subtypes (Figures 2D, 2E, S1B, and S1C). Each subtype displayed distinct morphological and innervation patterns, which matched between the two connectomes (Figure 2E). Moreover, since the fly brain is mostly symmetrical, besides matching morphologies on each side (Figure 2E), we also expected each type and subtype to have similar numbers of neurons across hemispheres, which is what we observed (Figure 2F). The major differences between hemispheres consisted of four neurons for the LB3d subtype in the male CNS and eight neurons for the LB3c subtype in FAFB – FlyWire. Furthermore, we could only identify the subtype LB2d in the male CNS dataset (Figure 2E). While these could represent sexual dimorphisms or inter-individual variability, these differences likely result from poorly reconstructed neurons due to the segmentation errors discussed above. As expected from the manual inspection, in the co-clustering, the LB1, LB2, and LB4 subtypes clustered together across datasets (Figure 2D). The exception was the LB3 subtypes, which is very likely due to technical challenges with segmenting these neurons. This is supported by the discrepancies observed in the counts of LB3a and LB3d GRNs between the two datasets.

**Figure 2.**
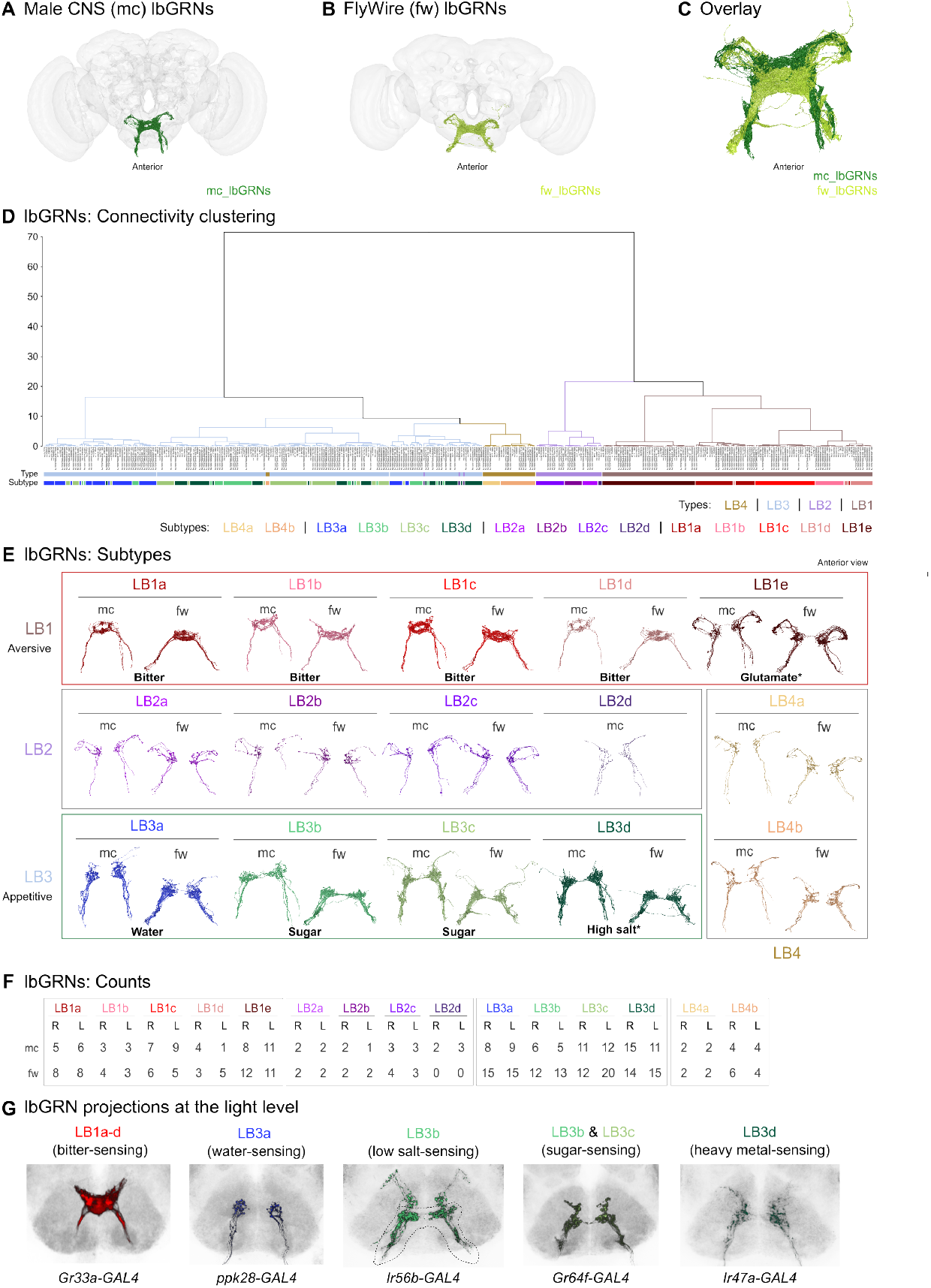
Cell typing the labellar bristle GRNs. **A**. Anterior view of the renderings of lbGRNs in the male CNS. **B**. Anterior view of the renderings of lbGRNs in FAFB – FlyWire. **C**. Anterior view of male CNS and FAFB – FlyWire lbGRNs overlaid in the male CNS space, showing the conserved broad anatomy of these neurons across different brains. **D**. Hierarchical co-clustering of male CNS and FAFB – FlyWire lbGRNs based on outputs using cosine similarity and a threshold of 2 synapses. Color bars represent the type (top) and subtype (bottom) defined for each male and female lbGRN according to the connectivity clustering for male CNS and FAFB – FlyWire neurons separately (Figures S1B and S1C). **E**. Anterior view of the renderings of lbGRNs per subtype across the right (R) and left (L) sides for male CNS (mc) and FAFB – FlyWire (fw), revealing their specific morphologies, which are preserved between different brains. The existence of LB2d only in the male CNS is very likely not due to a sexual dimorphism, but the result of technical difficulties due to automated segmentation. Outlined in red and green boxes are the types we propose to be, respectively, aversive and appetitive. The putative sensations of each subtype are also indicated. The asterisk (*) indicates the subtypes for which we do not show a receptor-GAL4 match but infer from the literature. **F**. lbGRN counts per subtype across hemispheres within each connectome. The challenges faced while reconstructing the male CNS lbGRNs have affected mostly the LB3a and LB3b subtypes. **G**. Semi-automated segmentation of confocal substacks for *Gr33a-, ppk28-, Ir56b-, Gr64f-* and *Ir47a-GAL4* positive lbGRNs were matched, respectively, to subtypes LB1a-d, LB3a, LB3b, LB3b-c and LB3d, suggesting they correspond to the bitter-, water-, low salt-, sugar- and aversive heavy metal ions-sensing lbGRNs.

Given the rich literature on the molecular and functional classes of lbGRNs, we next aimed to assign a receptor identity to as many types and/or subtypes identified in the connectome as we could. The sensing of the chemical environment by GRNs is mediated by specific receptors from four different receptor families, namely ionotropic receptor (IR), gustatory receptor (GR), transient receptor potential (trp), and pickpocket (ppk) channel families ^6,8,9,39,68–70^. Many of these receptors have been linked to sensing either a specific taste modality or particular pheromonal cues and resulting in specific behavioral outputs. We therefore generated highresolution confocal images of the projection patterns of specific receptor-GAL4 driver lines and matched them to our identified classes (Figure 2G).

The LB1a-d neurons project contralateral branches that form an interconnected ring in the midline, with specific variations depending on the subtype (Figure 2E). Using visual inspection in reference to anatomical landmarks, we decided that they best matched the *Gr33a-GAL4* driver projection pattern, which is expressed in all bitter-sensing GRNs (Figure 2G). Thus, we propose LB1a-d to be bitter-sensing subtypes that generate aversive responses. While in FAFB – FlyWire, these neurons were already annotated as bitter-sensing, our connectivity analysis across the two connectomes suggests that they can be further subdivided into the four proposed subtypes (LB1a-d). This is supported by the earlier observation of the existence of four different functional classes of bitter compound-responsive taste bristles, involving most S- and I-type bristles ^62^. Furthermore, L-type bristles were shown not to respond to bitter stimulation or house any GRN expressing bitter-sensing receptors ^62^. But recent data suggest that they generate a mild aversive response upon stimulation with specific amino acids like glutamate ^12^. This response has been linked to Ir94e-expressing GRNs ^12^, whose projection pattern matches the LB1e GRN morphology (Figure 2E). These neurons are mostly ipsilateral and have a single distinctive dorsolateral branch. While previously assigned a separate type ^35^, our connectivity analysis across multiple connectomes shows that they cluster together with canonical bitter-sensing GRNs, sharing specific downstream partners. We therefore propose that they are defined as the fifth LB1 subtype, making LB1 a broad aversive type.

Using similar lines of reasoning led us to propose that the LB3 type includes appetitive GRNs. The LB3a subtype morphology matched the projection pattern of *ppk28-GAL4* positive neurons, which have been proposed to be water-sensing GRNs ^71,72^ (Figure 2E and 2G). Both LB3b and LB3c projection patterns matched the *Gr64f-GAL4* positive neurons and are, thus, likely to correspond to sweet-sensing GRNs, mediating appetitive responses ^63^ (Figure 2E and 2G). Interestingly, the projections of LB3b neurons also seem to match the projection pattern of *Ir56b-GAL4* positive labellar GRNs, which have been shown to overlap with GRNs expressing sugar receptors and to be the main mediators of attractive responses to low salt ^64,73^ (Figure 2E and 2G). Morphologically, the LB3d GRNs match the glutamatergic *Ir7c-GAL4* and *ppk23-GAL4* positive neurons, which are involved in high salt avoidance responses ^65^. Additionally, they are morphologically similar to *Ir47a-GAL4* labelled neurons (Figure 2G), which were shown to overlap with *ppk23-GAL4* expression ^74^. The direct downstream connectivity of these neurons overlaps with appetitive subtypes, suggesting that the LB3d aversive GRNs might inhibit the activity of neurons also in attractive circuits via glutamatergic signaling.

Intriguingly, we have not found a receptor-GAL4 driver line that clearly matched the overall anatomies of the LB2 and LB4 types. Morphology-based comparisons had suggested that the LB2 type is related to LB1e neurons ^35^ (Figure S1F). However, our connectivity analysis using both connectomes revealed LB2 as a separate type that can be subdivided into LB2a-c, with the different subtypes having variations in their outputs and specific anatomies, which are observed across hemispheres in the two datasets (Figure 2D and 2E). Furthermore, our connectivity analysis and detailed anatomical comparisons defined LB4 as a novel GRN type that can be further divided into two subtypes, LB4a and LB4b. Broadly, these neurons are characterized by their mostly ipsilateral projections and a dorsolateral branch (Figure 2E). Specifically, LB4a neurons contain anterior branches, while LB4b neurons are, in relation, branching more ventrally and posteriorly. Existing annotations in FAFB – FlyWire assign LB4a neurons to the LB2 type and LB4b to the LB3 type. Close inspection revealed that LB4b neurons differ morphologically from LB3 neurons, as they project more dorsally (Figure S1E). Their overlapping outputs in both connectomes (Figures 2D and S1B and S1C) strengthen the conclusion that LB4a and LB4b neurons belong to the same type. In particular, these neurons share top downstream partners, like AN27X021 or PRW047, which also receive strong inputs from other GRN types from different subclasses.

In conclusion, by combining connectivity-based clustering within and between connectomes, anatomical comparisons, as well as matching with existing known molecular markers, we both identified known lbGRN types and extended the currently available annotations to include new subtypes matching known physiological and behavioral data. Intriguingly, our analysis also revealed a new type (i.e., LB4), which might represent novel, uncharacterized lbGRNs. Computational analyses described later in this paper support the functional relevance of this subdivision and also suggest what type of behaviors they might control.

### Cell typing the taste peg GRNs

tpGRNs have mostly been regarded as a uniform GRN subclass in the literature ^9,10,75^. Having not been cell typed before at the connectome level, we used an identical strategy to the one described above to type the tpGRNs we identified in the male CNS and FAFB – FlyWire datasets (Figures 3A-C and S2A). As we also show in a companion paper where we functionally characterize the involvement of these neurons in yeast feeding in female flies, tpGRNs consist of two cell types which differ in their morphology and direct downstream connectivity in both male and female brains ^49^ (Figures 3D, 3E, and S2B). One of the types, which we named dorsal tpGRNs (dtpGRNs), has dorsal projections that are absent in the other type, which we called claw tpGRNs (ctpGRNs) due to their claw-like projection patterns that are more evident in the FAFB – FlyWire ctp-GRNs ^49^. It is interesting to note that the anatomies of the two tpGRN types, while broadly similar between the two connectomes (Figure 3C), also have particular variations between the datasets that do not translate into evident connectivity differences (Figure 3D and 3E). This reflects well-known interindividual variabilities in the detailed projection patterns of gustatory neurons and highlights the advantage of including connectivity-focused analyses in the typing process to reveal functionally relevant cell types. ctpGRNs are the larger population in both datasets, but we only observe a bilateral surplus of ctpGRNs in FAFB – FlyWire (female), whereas dtpGRN counts are identical (Figure 3F). This matches the known taste peg sexual dimorphism and implies that the extra peg row, observed in females, is innervated exclusively by ctp-GRNs ^10^.

**Figure 3.**
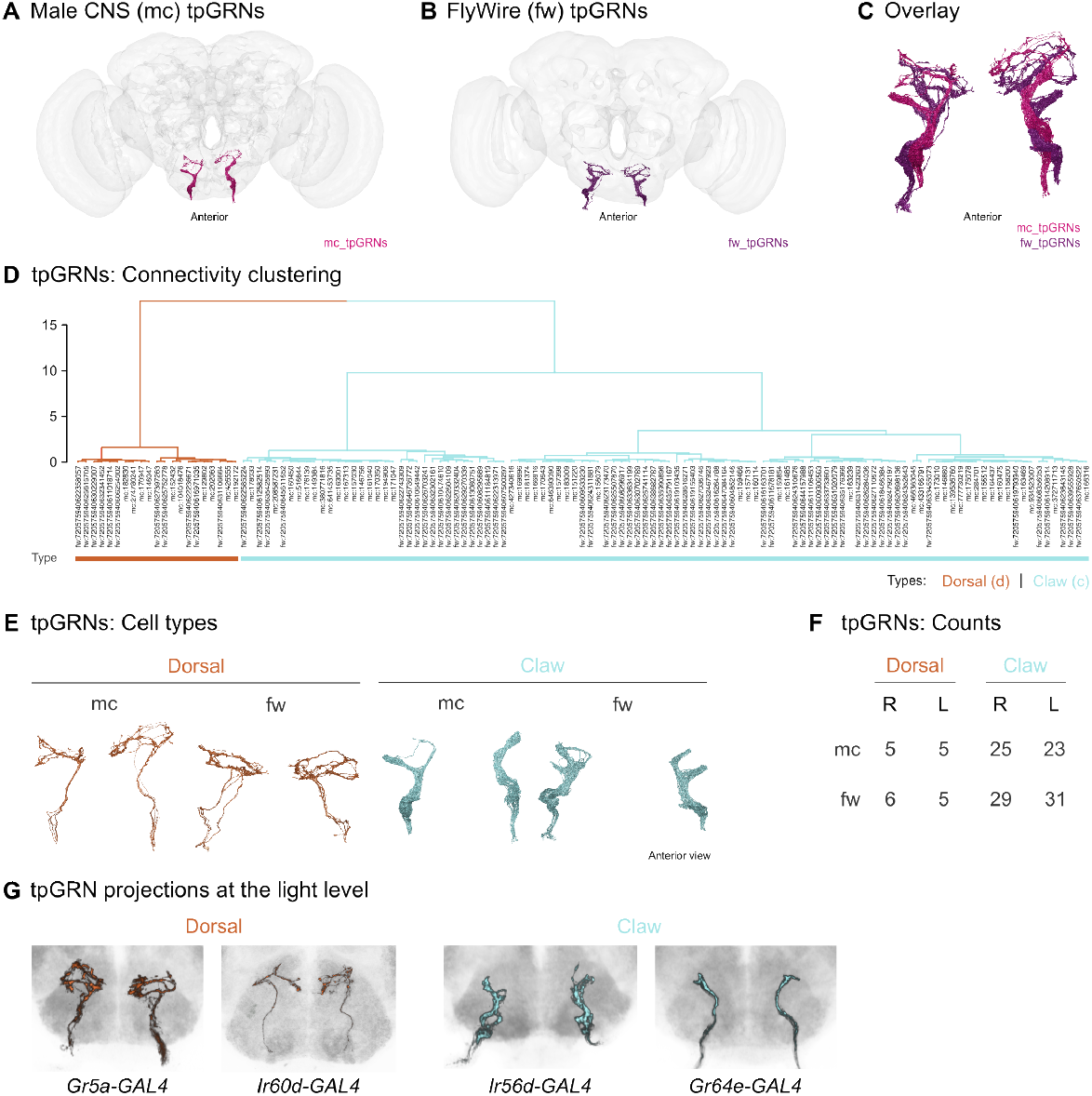
Cell typing the taste peg GRNs. **A**. Anterior view of the renderings of tpGRNs in the male CNS. **B**. Anterior view of the renderings of tpGRNs in FAFB – FlyWire. **C**. Anterior view of the male CNS and FAFB – FlyWire tpGRNs overlaid in the male CNS space. The overall anatomies are preserved, but inter-individual variabilities are highlighted via specific observable differences in the GRNs’ projection fields between the two brains. **D**. Hierarchical co-clustering of male CNS and FAFB – FlyWire tpGRNs based on outputs using cosine similarity and a threshold of 2 synapses, reveals two types, dorsal and claw tpGRNs, consistent with the connectivity clustering with solely the male and female tpGRNs (Figure S2). **E**. Anterior view of the renderings of the morphologically distinct dorsal and claw tpGRNs across the two hemispheres within male CNS and FAFB – FlyWire. Sparse anatomical variabilities between datasets within each type do not translate into direct connectivity differences. **F**. tpGRN type counts per side across male and female demonstrate that the reported extra peg row in female flies houses claw tpGRNs. **G**. Semi-automated segmentation of confocal substacks for *Gr5a-* and *Ir60d-GAL4* positive tpGRNs were matched specifically to the dorsal tpGRNs, while *Ir56d-* and *Gr64e-GAL4* positive tpGRNs were matched to claw tpGRNs.

At the molecular level, we propose that dtpGRNs are labeled by *Gr5a-* and *Ir60d-GAL4*, while ctpGRNs match both *Ir56d-* and *Gr64e-GAL4* projection patterns (Figure 2G). Functionally, we had previously shown that ctpGRNs specifically play a role in sensing carbonation through IR56d ^9^. Additionally, this tpGRN type might also be responsive to fatty acids, via IR56d, and to glycerol, as it is likely to express the glycerol receptor GR64e ^76,77^. In a companion paper, we functionally dissect a dtpGRN to a swallowing-controlling proboscis MN pathway that modulates a specific feeding microstructure parameter, the burst length, when amino acid-deprived flies feed on proteinaceous food ^49^. Our functional dissection of how these two tpGRN populations differentially control feeding, together with their distinct receptor profiles and other reported specific roles, are important experimental validations for the two tpGRN types we have defined at the connectome level.

### Cell typing the pharyngeal GRNs

phGRNs have not yet been comprehensively identified and cell typed at the connectome level. To do so, we used the same strategy as described for the other GRN subclasses (Figures 4A-C and S3A). All our connectivity clustering analyses, either with only the male CNS (Figure S3B), FAFB – FlyWire (Figure S3C), or the co-clustering of phGRNs from both datasets (Figure 4D), suggested the existence of 16 phGRN cell types, which we named PhG1-16. The PhG1 type was the only one we further subdivided into three subtypes, PhG1a-c, based on their specific connectivity and morphology. In contrast to lbGRNs and tpGRNs, phGRNs cell types are generally composed of either one or two right-left pairs of neurons (Figure 4F). It is striking to note that the terminal arborizations of each ph-GRN type and subtype are unique but mostly preserved in both connectomes and across hemispheres within the same brain (Figure 4E). When we overlay the transformed FAFB – FlyWire phGRN meshes with the male CNS ones, this morphological conservation can also be seen at the whole population level (Figure 4C). Most phGRN cell types enter the SEZ via the aPhN, coming from either the LSO or the VCSO ^36^, and have contralateral projections, except for a few types, which are exclusively ipsilateral (Figure 4E). Only PhG7-9 neurons project through the PhN, originating in the DCSO ^36^, and all send arborizations across the midline. We propose that a neuron from each of the three types is housed in each of the two DCSO sensilla on either side. This is supported by a detailed receptor-to-neuron map that revealed a mostly unique receptor profile for every phGRN ^39^. For the DCSO specifically, it was shown that each of the three GRNs within a sensillum expresses specific receptors, in a similar arrangement across both the ventral and dorsal sensilla ^39^. We have matched the *Ir10a-GAL4* driver to PhG9 (Figure 4G), one of the GRN types housed in the DCSO ^9^. Because of its characteristic morphology, we have also identified PhG9 as corresponding to the reported *Ir100a-GAL4* positive phGRNs ^9,39^. Both *Ir10a-* and *Ir100a-GAL4* have additionally been found to label phGRNs in the LSO (sensillum #8) ^9,39^, which we have morphologically matched to PhG11, a GRN type projecting through the aPhN.

**Figure 4.**
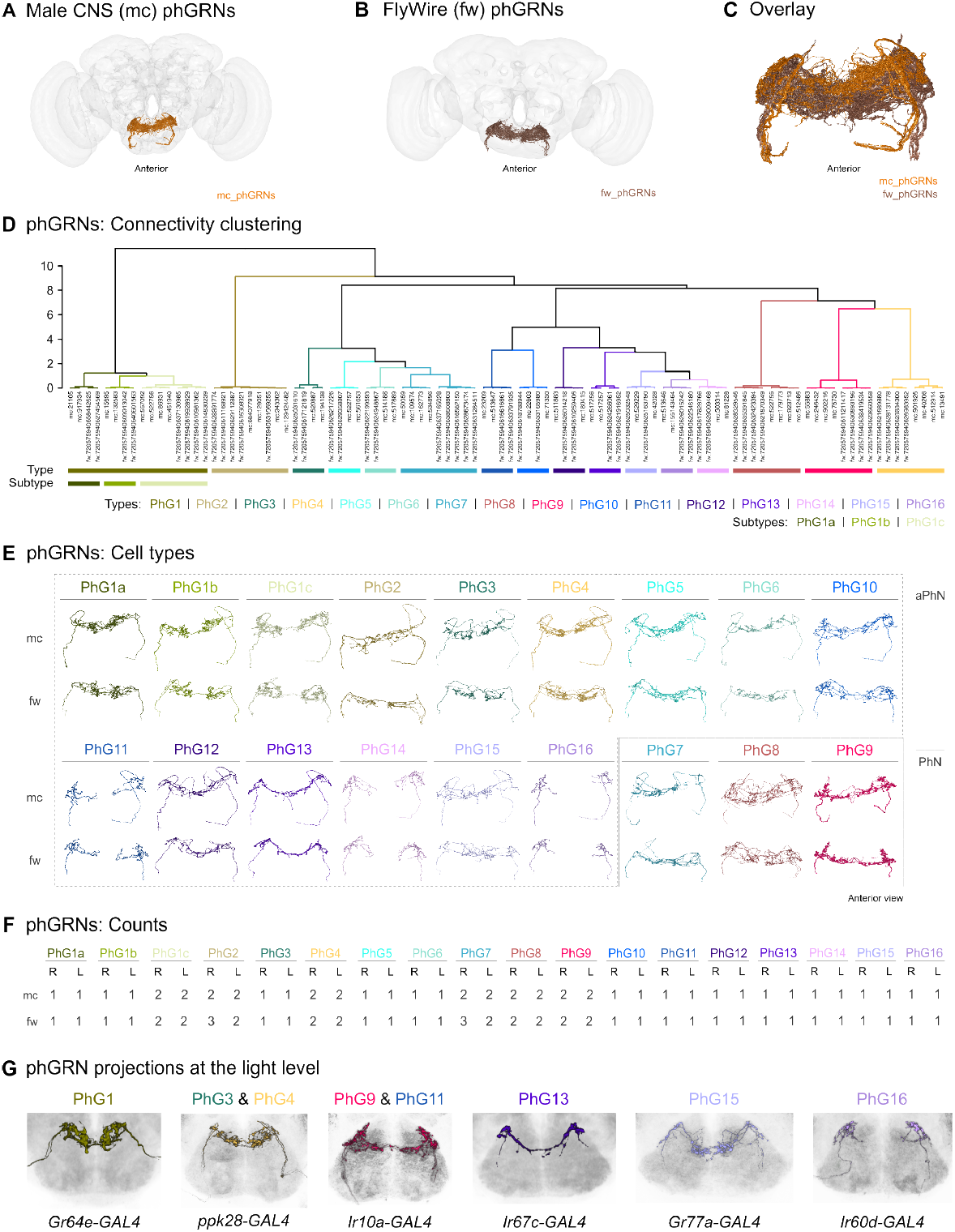
Cell typing the pharyngeal GRNs. **A**. Anterior view of phGRN renderings in the male CNS, showing their terminal arborizations in the SEZ. **B**. Anterior view of phGRN renderings in FAFB – FlyWire. **C**. Anterior view of the male CNS and FAFB – FlyWire phGRNs overlaid in the male CNS space, demonstrating the morphological preservation of this GRN subclass across brains. **D**. Hierarchical co-clustering of male CNS and FAFB – FlyWire phGRNs based on outputs using cosine similarity and a threshold of 5 synapses. Each neuron is colored in accordance with its type (top) and subtype (bottom), the latter only applied to PhG1. Male and female neurons of the corresponding types cluster together (Figure S3B and S3C). **E**. Anterior view of renderings of aPhN- and PhN-projecting phGRN types, the latter corresponding to PhG7-9, reveal unique morphologies that are conserved between different brains. **F**. phGRN counts per side compared between male CNS and FAFB – FlyWire shows a conservation per type and subtype across brains. **G**. Semi-automated segmentation of confocal substacks for *Gr64e-, ppk28-, Ir10a-, Ir67c-, Gr77a-* and *Ir60d-GAL4* positive phGRNs matched, respectively, to the PhG1, PhG3&4, PhG9&11, PhG13, PhG15 and PhG16 types.

By combining the receptor-to-neuron identification from the literature with our phGRN type matched to receptor-GAL4 projection patterns, we have further refined the mapping of GRN types to other specific LSO and VCSO sensilla. Here, we have matched *Gr64e-, ppk28-, Ir67c-, Gr77a-* and *Ir60d-GAL4* driver projection patterns to PhG1, PhG3&4, PhG13, PhG15 and PhG16, respectively (Figure 4G). PhG1 should, thus, correspond to neurons in the VCSO and LSO sensillum #7. This latter sensillum should also house PhG13. PhG13 is a particularly interesting type, as we have found it to be potentially sexually dimorphic in its outputs ^60^. One of its top partners in the male CNS is AN05B035, a putative male-specific ascending neuron that innervates the T1 segment of the VNC and projects to the SEZ, sending branches to a region surrounding the esophagus thought to be sexually dimorphic ^46,78^ (Figure S3D). In females, this phGRN type has been proposed to be involved in a circuit that integrates pheromone and taste information to control receptivity ^79^. PhG3&4 should be housed in the LSO and VCSO, while PhG15 and PhG16 should, respectively, correspond to a neuron in VCSO and LSO sensillum #7.

In line with published receptor profiles and our cell typing, the emerging picture is that phGRNs are the most diverse GRN type in terms of morphology and direct downstream connectivity. This diversity supports the idea that these neurons generate unique responses and play unique roles in controlling ingestion.

### Cell typing the leg bristle ascending GRNs

Anatomically, leg bristle GRNs can be divided into the ones that directly project to the brain (lgAGRNs) and the ones that arborize locally in the VNC (lgLGRNs) (Figure 1B and 1C). As an intact CNS, the male CNS dataset has allowed us, for the first time, to completely reconstruct the lgAGRNs (Figures 5A and S4A). We focused our comparative analysis on the FAFB – FlyWire lgAGRNs, given that these include the SEZ terminal arborizations, which capture most of their downstream targets. We identified the SA_VTV_1-5 and SA_VTV_7-10 types in FAFB – FlyWire as corresponding to lgAGRNs (Figure 5B and 5C). Separate clustering of lgA-GRNs based on their output connectivity in the female and male connectomes revealed nine lgAGRN types (LgAG1-9) (Figure S4B and S4D). When we performed the co-clustering of the neurons from both connectomes, the male and female neurons of the corresponding types clustered together, except for one male and one female LgAG4 neuron (Figure 5D). Differences in morphology did not influence their downstream connectivity (Figure 5C-E). The counts of each lgAGRN type are comparable between connectomes and hemispheres in the same connectome, with one neuron being the biggest difference found, and LgAG9 (the type with the fewest GRNs), having only one neuron identified in FAFB – FlyWire (Figure 5F).

**Figure 5.**
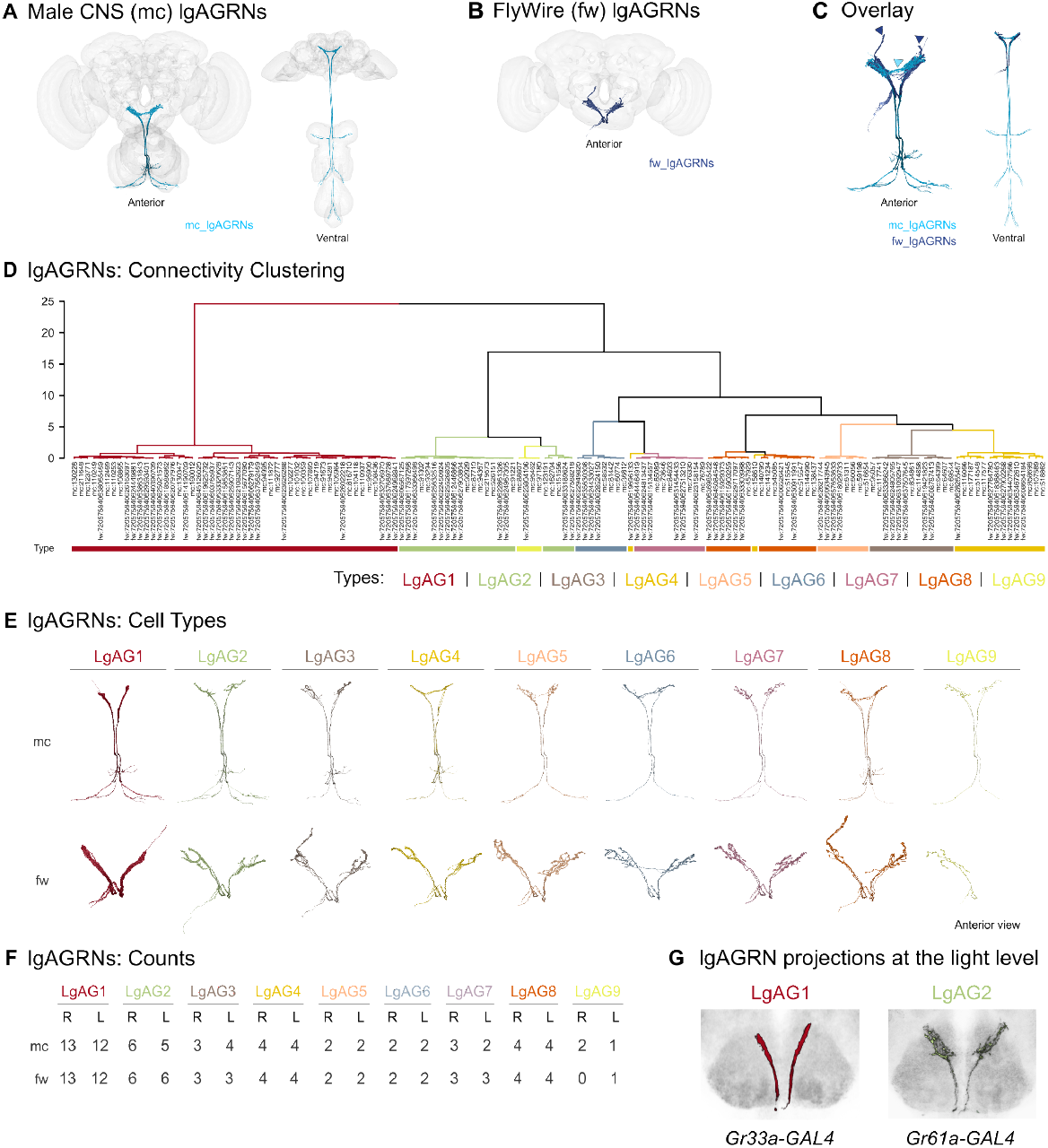
Cell typing the leg bristle ascending GRNs. **A**. Rendering of lgAGRNs in the male CNS in two different views, anterior and ventral, highlighting, respectively, the SEZ projections and axonal entry points in the VNC. **B**. Rendering of lgAGRNs in FAFB – FlyWire in an anterior view. In FAFB – Flywire, these neurons are restricted to their SEZ terminal arborizations. **C**. Anterior and ventral views of the male CNS and FAFB – FlyWire lgAGRNs overlaid in the male CNS space, revealing anatomical variabilities in their SEZ projections across the two connectomes (arrowhead). **D**. Hierarchical co-clustering of male CNS and FAFB – FlyWire lgAGRNs based on outputs using cosine similarity and a threshold of 2 synapses. Each neuron is colored in accordance with its type, which was defined by connectivity clustering the male and female neurons separately (Figures S4B and S4D). **E**. Anterior view of each lgAGRN cell type, LgAG1-9, rendering across male CNS and FAFB – FlyWire. Inter-individual morphological variabilities do not translate into direct downstream connectivity differences. **F**. LgAG1-9 counts per side are essentially consistent across the two connectomes. **G**. Semi-automated segmentation of confocal substacks for *Gr33a-* and *Gr61a-GAL4* lgAGRN positive populations matched, respectively, to the LgAG1 and LgAG2 types.

A great advantage of working with the male CNS dataset was that we were able to identify from which legs these types project from based on their VNC projections (forelegs to T1, midlegs to T2, and hindlegs to T3, Figure S4C). It is interesting to note that two types, LgAG1 and LgAG2, are found in all six legs. LgAG3, LgAG4, and LgAG8 are the three types found only in taste bristles on the mid- and hindlegs. And, the remaining ones, LgAG5, LgAG6, LgAG7, and LgAG9 project mainly from forelegs’ taste bristles. Although in previous studies, the expression of specific receptors has been mapped solely to foreleg GRNs, we could not find reports of receptors being exclusively expressed in mid- and hind-leg GRNs ^7,80^. It is therefore possible that the types restricted to mid- and hindleg GRNs do not differ in the receptors they express, but are connected differently to convey, for example, information about the location of gustatory stimuli.

From our effort to attribute molecular identities to these cell types, we have been able to match the *Gr33a-GAL4* driver projection pattern to LgAG1 neurons (Figure 5G). This suggests that these GRNs sense aversive compounds, generating an avoidance response ^2^. One of its top downstream partners is ANXXX470, better known as TPN3, a neuron which is required for conditioned taste aversion ^81^. These neurons are also supposed to express GR32a and sense secreted cuticular hydrocarbons to inhibit conspecific male and interspecies courtship ^2,82,83^. In addition, we have also matched *Gr61a-GAL4* positive lgAGRNs to the LgAG2 type, suggesting that this GRN type senses appetitive tastants and, thus, generates an acceptance response (Figure 5G).

Interestingly, as we observed for PhG13 GRNs, LgAG8 neurons also connect to AN05B035, a potentially male-specific neuron, suggesting, thus, that these neurons have sexually dimorphic outputs and could play important roles in sexual behaviors.

### Cell typing the wing bristle GRNs

For VNC-projecting GRNs, like the wGRNs, we compared neurons identified in the male CNS (Figure 6A) with the corresponding MANC neurons (Figure 6B), allowing us to account for interindividual differences in the wiring of the male gustatory system. Connectivity clustering using cosine similarity based on the wGRNs’ outputs revealed four types, which we named WG1-4 in both datasets (Figure S5A and S5B). The co-clustering of the male CNS and MANC wGRNs resulted in the clustering of the corresponding male CNS and MANC types in the same groups, except for WG3 (Figure 6C). In contrast to other GRN subclasses, the different wGRN types are basically formed by the same number of neurons (48-50 GRNs on each side) with minuscule deviations between connectomes and sides (Figure 6E). While their connectivity differs, morphologically, the different wGRN types are almost indistinguishable from each other (Figure 6D). However, if all the types are overlaid, it is possible to see that they innervate different regions in the neuropil, with WG1 innervating the most anterior area compared to the other types (Figure 6F).

**Figure 6.**
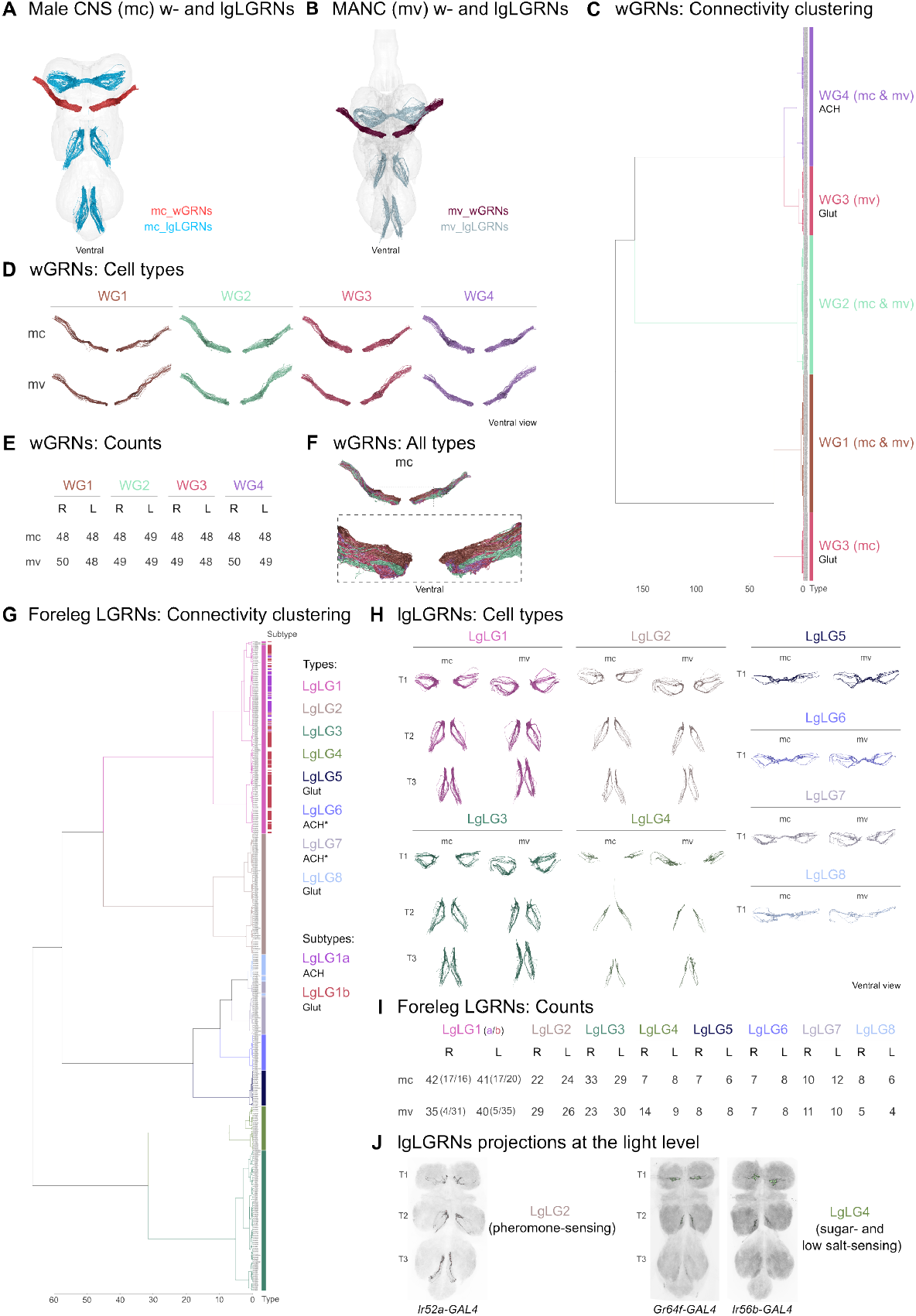
Cell typing the wing and leg bristle local GRNs. **A**. Ventral view of wGRN and lgLGRN renderings in the male CNS. While wGRNs arborize solely within T2 in the VNC, lgLGRNs project across all thoracic neuromeres, depending on whether they are housed in the fore-, mid- and hindleg taste bristles. **B**. Ventral view of wGRN and lgLGRN renderings in MANC. **C**. Hierarchical co-clustering of male CNS and MANC wGRNs based on outputs using cosine similarity and a defined threshold of 2 synapses. Neurotransmitter predictions are indicated. **D**. Ventral view of each wGRN cell type, WG1-4, rendering across the male CNS and MANC. **E**. wGRN type counts per side, which are essentially the same between types, are consistent across both male connectomes. **F**. Ventral view of all male CNS wGRN types overlaid to reveal their different projection fields in the neuropil. Bottom: zoomed in view of the boxed region on top. **G**. Hierarchical co-clustering of male CNS and MANC foreleg local GRNs based on outputs using cosine similarity and a defined threshold of 2 synapses. The neurons are colored according to their type (left), LgLG1-8. LgLG1 type neurons are subtyped as LgLG1a and LgLG1b, depending on whether they are predicted to release acetylcholine or glutamate. The respective co-clusterings of mid- and hindleg GRNs can be found in Figure S5C and S5E. **H**. Ventral view of the renderings of lgLGRN types across male CNS and MANC. LgLG1-4 span the T1-3 VNC segments while LgLG5-8 correspond exclusively to foreleg GRNs. **I**. Foreleg local GRN type counts per side are within the same range across the two connectomes. **J**. Semi-automated segmentation of confocal substacks for for *Ir52a-, Gr64f-*, and *Ir56b-GAL4* lgLGRN positive populations matched, respectively, to LgLG2, LgLG4, and LgLG4.

Because the anatomies of the different types are almost identical, we employed a different strategy to propose a molecular identity and, thus, a potential functional role for each type. When possible, we used the top downstream partners of the wGRN types that have been described in the literature to attempt an identification. PPN1, which is annotated as AN05B102a in the male CNS and as AN_AVLP_20 in FAFB – FlyWire, has been described as a courtship-promoting neuron in males ^84^. This neuron is activated by VGlut+/ fru+/ ppk23+/ ppk25+ GRNs, which have been shown to project from the wing taste bristles ^84^. Likewise, VGlut-/ fru+/ ppk23+/ ppk25-GRNs have also been shown to play a role in courtship behavior and exist in equal numbers across the same wing taste bristles ^84^. Besides other overlapping outputs, PPN1 is one of the top downstream partners of both the WG3 and WG4 GRN types, and since their predicted neurotransmitter is, respectively, glutamate and acetylcholine in both connectomes, we propose that the WG3 type corresponds to the VGlut+/ fru+/ ppk23+/ ppk25+ GRNs and the WG4 type to the VGlut-/ fru+/ ppk23+/ ppk25-GRNs. A different set of wGRNs, which also plays a role in courtship behavior, has been demonstrated to express fruitless and IR52a, with some neurons also expressing IR52b ^46,78^. Trans-Tango-based circuit tracing using both *Ir52a-* and *Ir52b-GAL4* has revealed that directly downstream of these GRNs, only in the male brain, there are neurons that have projections around the esophagus, which have been described to be sexually dimorphic ^46,78^. Besides WG3 and WG4, only the WG1 GRN type has those types of sexually dimorphic neurons as relevant partners (Figure S6A), leading us to propose that these GRNs are the previously identified courtship-promoting fru+/ Ir52a+ GRNs ^46,78^. Moreover, the literature suggests that wGRNs express sugar GRs, like GR43a, GR64f, and GR5a, and respond to sugar stimuli ^41,45,85,86^. A key downstream partner of WG2 is Dandelion, an AN typed as AN13B002 in the male CNS and as AN_GNG_68 in FAFB – FlyWire ^47^. Given that Dandelion is also a key downstream neuron of LB3b GRNs in the labellum, which we also propose to express sugar GRs, we propose that the WG2 GRNs are likely to detect sugar and trigger feeding. This approach of using key shared downstream partners to assign shared functions across the connectome, therefore, emerges as an important strategy to propose similar functional roles to different GRN cell types across multiple subclasses. A strategy we will employ more comprehensively in later analyses.

A last important observation is that all the defined male wGRN types connect strongly to specific second-order neurons, which are morphologically different between male CNS and FAFB – FlyWire and, thus, are likely to be sexually dimorphic (Figure S6A). AN05B102d and ANXXX151 (matching, respectively, AN_multi_69 and AN_multi_71 in FAFB – FlyWire), two AN types that are directly downstream of WG1, have projections around the esophagus, a region that has been suggested to be sexually dimorphic ^46,78^. The same occurs for another output AN of WG3 and WG4 GRNs, annotated as AN05B102c in the male CNS and AN_AVLP_PVLP_5 in FAFB – FlyWire. The WG2 GRNs connect to ANXXX013, which has been matched to AN_GNG_150 in FAFB – FlyWire and has an extended dorsal projection in the male CNS brain outside the SEZ. Taken together, these analyses reinforce the idea that wGRNs are sexually dimorphic in terms of their downstream connectivity and play a key role in mediating sexually dimorphic social behaviors in flies.

### Cell typing the wing bristle GRNs

Similar to the wGRNs, to cell type the lgLGRNs we used the neurons identified in the male CNS (Figure 6A) and MANC (Figure 6B). Because these neurons span the T1-3 segments of the VNC, we did the connectivity clustering separately for each pair of fore-, mid-, and hindlegs, which project to T1, T2, and T3, respectively. From the co-clustering of male CNS and MANC lgLGRNs, we defined four types, LgLG1-4, which are found across all legs (Figures 6G, S5C, and S5E). All the GRNs from these types have ipsilateral projections, terminating within the leg neuropil on the same side that they enter the VNC. Each of the four types has a characteristic morphology at the population level, arborizing in different regions of the neuropil (Figures 6H and S5G). The morphology of LgLG4 GRNs is the most distinct of the four types. The LgLG1 type is the only type we have further subtyped (Figures 6G, S5C, S5E and S5H). Given that they are predicted to release different neurotransmitters, we divided LgLG1 into LgLG1a and LgLG1b, respectively, depending on whether the GRN is predicted to express acetylcholine or glutamate. Considering their morphology, LgLG1a and LgLG1b seem to overlap in the same regions in the leg neuropils (Figure S5H). In terms of GRN counts in both connectomes, LgLG1 is the type with the most neurons, while LgLG4 is the type with the fewest GRNs across the fore-, mid-, and hindlegs (Figures 6J, S5D, and S5F). An observation regarding MANC is that there seems to be an overestimation of LgLG1b GRNs compared to LgLG1a, if we consider that the numbers of the two subtypes are essentially the same in the male CNS.

To assign a molecular identity to these lgLGRN types, we once more used top downstream partners that have been described in the literature and associated with GRNs expressing specific receptors and playing determined behavioral roles. Equal to the WG3 and WG4 types, LgLG1 GRNs also have PPN1 as their top downstream partner. Since both VGlut-/ fru+/ ppk23+/ ppk25- and VGlut+/ fru+/ ppk23+/ ppk25+ GRNs have been mapped to leg taste bristles ^84^, we propose that those molecular identities correspond to the LgLG1a and LgLG1b types. Similar to WG1 GRNs, the LgLG2 type has output neurons that extend projections to the sexually dimorphic region around the esophagus, and since IR52a and IR52b have been mapped to the same lgLGRNs, we propose that LgLG2 corresponds to fru-/ Ir52a+/ Ir52b+ GRNs ^42^. In addition, at the light microscopy level, we have anatomically matched the LgLG2 type to *Ir52a-GAL4* positive lgLGRNs (Figure 6I). Similar to WG2, LgLG3 GRNs have Dandelion as one of their top downstream partners. We therefore propose that it is likely to express sugar GRs, especially Gr5a, which is expressed by lgLGRNs ^85,86^. LgLG4 also has Dandelion as a relevant second-order neuron, and it shares other outputs, like DNg103, with other appetitive GRN types. We have matched the anatomy of LgLG4 GRNs to GR64f- and *Ir56b-GAL4* positive lgLGRNs (Figure 6I), which have been shown to generate attractive responses, respectively, to sugar and salt ^4,41,73^. Taken together, we propose that the LgLG4 type is likely to mediate appetitive sugar and low salt responses.

The co-clustering of lgLGRNs from the two connectomes revealed four additional cell types, LgLG5-8, which project exclusively from the forelegs to T1 (Figures 6G, S5C, and S5E). The neurons comprising these types have mostly contralateral projections, crossing the midline and innervating the neuropil on the opposite side of their entrance into the VNC (Figure 6H). It is well known that these contralaterally projecting neurons are sexually dimorphic, projecting from taste bristles that exist solely in males ^84^. They have been shown to play a role in contact pheromone sensation. The number of neurons within these types are comparable across both connectomes and between both sides of the same VNC (Figure 6J). In the male CNS, the predicted neurotransmitter for the LgLG5 and LgLG8 GRNs is glutamate, while for the LgLG6 and LgLG7 GRNs it is acetylcholine (Figure 6G). However, in MANC, all four types are predicted to express glutamate. Assuming there is some bias towards glutamate when predicting the neurotransmitter in MANC neurons, we propose LgLG5 and LgLG8 to be VGlut+/ fru+/ ppk23+/ ppk25+ GRNs and LgLG6 and LgLG7 to be VGlut-/ fru+/ ppk23+/ ppk25-GRNs. Bilateral GRN populations with either one of those molecular identities have been previously shown to exist in males ^84^. Interestingly, PPN1 is not a relevant second-order neuron of any of these cell types. However, many of their downstream partners are male-specific or sexually-dimorphic ANs (Figure S6B), supporting our proposal that these are male-specific pheromone-sensing neurons, mediating sexually dimorphic social behaviors.

Taken together, our cell typing effort of GRNs across three connectomes has allowed us to both identify known GRN types previously described using molecular, anatomical, physiological, and behavioral approaches, and to expand their classification into functionally relevant new cell types, as exemplified by our companion paper focusing on tp-GRNs ^49^. We also characterized inter-individual differences across datasets, as well as known and novel sexual dimorphisms. Thus, our GRN cell typing should represent the first comprehensive annotation and characterization of the gustatory system in any insect. This provides a unique opportunity to further explore the circuit basis of how these GRNs might mediate specific behaviors.

### Functional organization of the gustatory connectome

The activity of GRNs across different body parts should be integrated in downstream circuits to orchestrate the actions underlying complex feeding decisions ^19^. For example, during foraging, leg bristle GRNs slow locomotion upon food encounters and trigger proboscis extension to probe food quality together with labellar bristle GRNs. If the food is appetitive, the fly remains at the food spot and initiates food ingestion. Labellar bristle and taste peg GRNs get sequentially activated by food, while the swallowed nutrients are monitored by pharyngeal sensilla. To study GRN populations that converge on similar downstream circuits, and therefore likely encode similar valence and/or mediate similar behaviors, we performed UMAP embedding of the SEZ-projecting GRNs in male CNS and FAFB – FlyWire based on their immediate downstream connectivity (Figure 7A). Importantly, we included only the monomorphic second-order neurons to focus on common pathways in both sexes. Including dimorphic neurons did not change the clustering, as the number of dimorphic partners is much lower than the monomorphic partners in general ^60^. Our UMAP embedding clustered GRNs of the same type from both datasets together, validating both our cell typing as well as the clustering.

**Figure 7.**
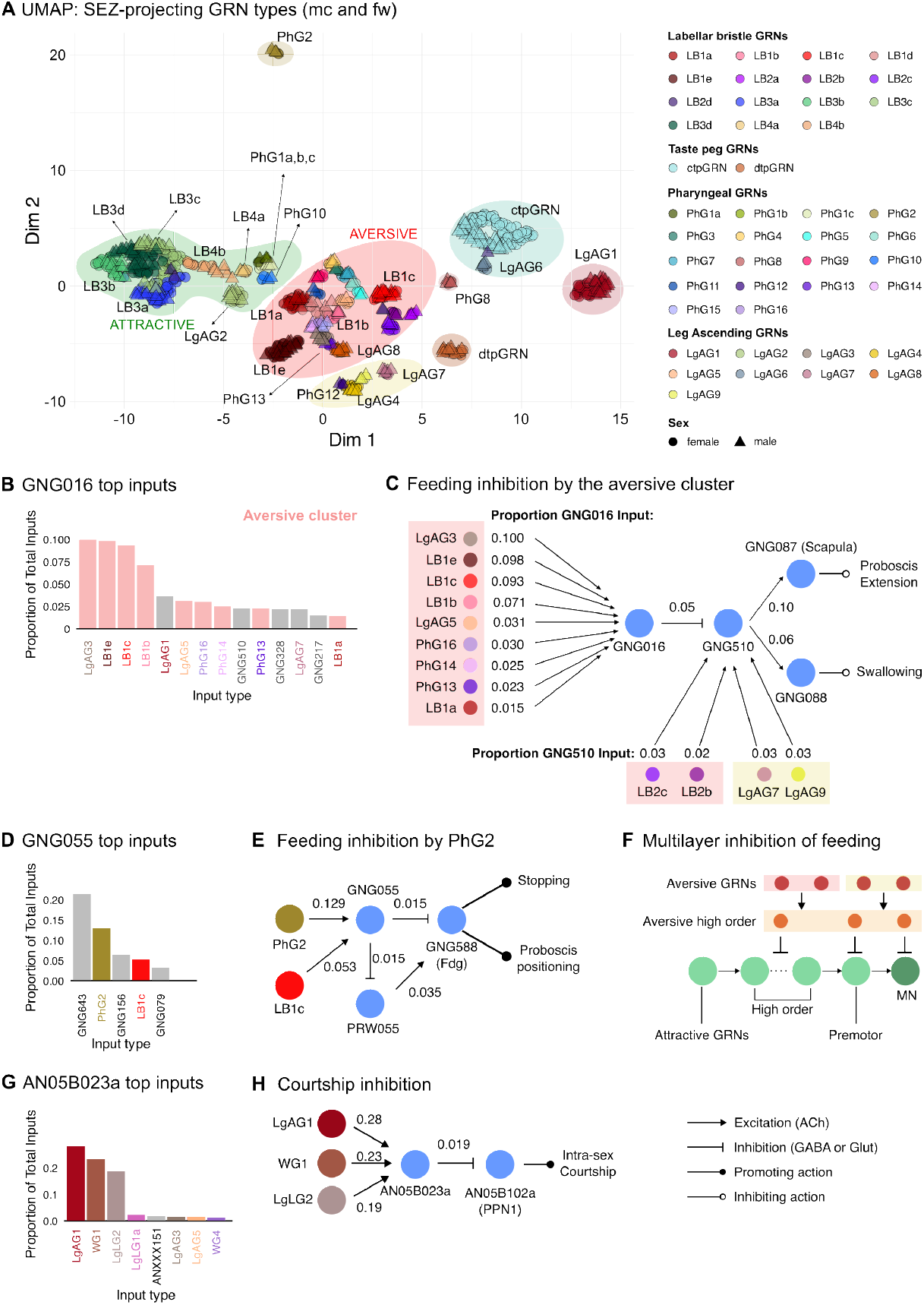
Functional organization of the gustatory connectome. **A**. UMAP embedding of SEZ-projecting GRN types from the male CNS and FAFB – FlyWire datasets based on their downstream connectivity. Each GRN is colored by its type. The shapes represent the sex of the animal. Circle: female (FlyWire), triangle: male (male CNS). Key GRN types in each cluster were labeled. Different clusters are encircled with distinct colors. The aversive cluster (red) and the attractive cluster (green) are labeled accordingly. Dim 1: UMAP dimension 1, Dim 2: UMAP dimension 2. **B**. Top GRN inputs to the second-order neuron, GNG016 shown as proportions of total inputs to GNG016. Input GRN types that are found in the aversive cluster in A are colored (red). **C**. The sensorimotor pathway mediating feeding inhibition by the aversive cluster at different circuit levels. Each GRN is colored according to its type. Red and yellow rectangles represent the aversive clusters in A. Each circle is a cell type. Synaptic weights are given as proportions of total inputs on downstream neurons. If a downstream cell type was previously identified in the literature, its name is indicated (e.g., Scapula). Otherwise, male CNS cell types are used. Scapula was proposed to inhibit feeding by inhibiting premotor neurons. GNG088 is predicted to be a GABAergic premotor neuron upstream of MNs controlling swallowing. **D**. Top inputs of GNG055, as described in B. GRN types, are highlighted by their respective colors. **E**. A parallel pathway for feeding inhibition via PhG2 and LB1c. Synaptic weights are given as proportions of total inputs on downstream neurons. **F**. Circuit model for nested inhibition of feeding by a diverse set of aversive GRN types. Feeding is inhibited at multiple circuit layers ranging from higher order to premotor to motor neurons (See Figure S8). **G**. Top inputs of another second-order neuron, AN05B023a as described in B. GRN types are highlighted by their respective colors. **H**. Candidate male – male courtship inhibition pathway. Synaptic weights are given as proportions of total inputs on downstream neurons. A cell type is considered excitatory if it is predicted to be cholinergic (ACh), inhibitory if predicted to be GABAergic, or glutamatergic.

Our UMAP analysis revealed eight main clusters (Figure 7A). One of the main clusters contains the bitter-sensing LB1 subtypes (Figure 2), and we therefore decided to name it the aversive cluster (Figures 7A and S7A). It also contains several phGRN types (PhG3-7, 9, 11, 13, 14, 15, and 16) and lgAGRN types (LgAG3, LgAG5, and LgAG8). As the pharynx is an important checkpoint for assessing food quality before the passage into the digestive system, we propose that these phGRN types are important for limiting food intake. Indeed, it has been shown that different combinations of phGRNs are required to suppress food intake and avoid bitter compounds ^87^. Functional studies dissecting the common circuitry downstream of the aversive phGRNs will be important to unravel the mechanisms of the combinatorial logic underlying pharyngeal sensorimotor transformations. While we could not assign a function to the LB2a-d types based on known receptor expressions, these are found in this cluster, suggesting that they might mediate aversive responses. Interestingly, despite having sexually dimorphic outputs, PhG13 GRNs from both males and females are also found in this cluster. We propose that, similar to LgAG1s ^2,82^, IR67c positive neurons might contribute to avoiding ingesting bitter compounds as well as regulating sexual behavior.

We deduced that another major cluster must contain GRNs mediating attractive behaviors, as it encompasses all LB3 GRNs, which we proposed to correspond to sugar and other phagostimulatory GRNs (Figures 2, 7A, and S7B). LB4a and b, which we could not assign a molecular identity, are also in this cluster, suggesting that they might convey attractive information (Figure S7B). Two pharyngeal GRN types, PhG1 (including all subtypes) and PhG10, are also in this attractive cluster. This agrees with our receptor-GAL4 matching, suggesting that PhG1s express GR64e (Figure 4G), which is important for detecting glycerol and inducing proboscis extension towards it ^2,39,63,76^. Similarly, the LgAG2 type, an lgAGRN type we have matched to *Gr61a-GAL4* positive neurons, is also part of this cluster. We therefore predict that neurons of this type sense appetitive compounds.

LgAGs 4, 7, and 9, and PhG12 comprise another cluster (Figure 7A). Although we cannot assign a clear role to the neurons in this cluster, we propose that these GRNs also mediate aversive responses, as they share GNG016 as a downstream partner with GRNs in the other major aversive cluster (Figure 7B).

ctpGRNs and LgAG6 form a unique cluster with unknown behavioral relevance (Figure 7A). Likewise, PhG2, PhG8, and dtpGRNs also form separate clusters. In an accompanying paper, we show that dtpGRNs anticipatorily facilitate ingestion to increase the length of yeast feeding bursts and therefore protein intake ^49^. Finally, LgAG1 GRNs from both male CNS and the FAFB – FlyWire datasets form a unique cluster (Figure 7A). As we mentioned above, this GRN type is important for sexual behavior in *Drosophila* as well as aversive responses.

Because these clusters arise from GRNs that converge onto common downstream neurons, they provide a unique entry point for predicting the functional logic of the circuits and the behavioral programs they underlie. GRNs from the major aversive clusters, as well as LgAG1, which was shown to be important for avoidance ^2^, synapse with GNG016 (Figure 7B and 7C). Of note, GNG016 is predicted to be a GABAergic neuron, and it has feedback connections with the aversive GRNs it receives inputs from (Figure S8A). This feed-back mechanism could, for example, serve as gain control, sharpening signals, or creating oscillations. However, the functional relevance can only be answered with careful experimental dissections. Our analysis shows that GNG016 signals to GNG510 (Figures 7C and S8B), which outputs onto GNG087 (Scapula) (Figures 7C and S8C). Interestingly, Scapula was proposed to inhibit proboscis extension by directly suppressing the activity of a premotor neuron ^47^. GNG510 also connects to GNG088 (Figures 7C, S8C and S8E), a GABAergic neuron that we predict to directly inhibit proboscis motor neurons that are involved in pharyngeal pumping (Figures 7C, 8E, and S8F). Interestingly, GNG510 receives direct inputs from GRNs from both aversive clusters, including LB2b, LB2c, LgAG7, and LgAG9 (Figures 7C and S8B). Moreover, Scapula also receives direct inputs from another aversive GRN type, LB1c (Figure S8D), demonstrating that the aversion circuits receive GRN inputs across multiple layers. Together, this circuit suggests that aversive gustatory cues are likely to suppress feeding by inhibiting proboscis extension and food swallowing.

This aversive circuit is not unique, as other parallel circuits inhibiting feeding can be identified. A good example is the circuit downstream of PhG2 (Figure 7E). GNG055 receives strong inputs from PhG2 and LB1c and is predicted to inhibit Fdg (GNG588), which has been previously shown to be an important neuron coordinating feeding and halting forward locomotion ^23,47,88^. Therefore, our findings suggest that aversive GRNs suppress feeding at different, parallel levels (Figure 7F). They influence specific downstream neurons that inhibit feeding at different levels of the sensorimotor arc, ranging from higher-order to premotor to motor neurons.

Given its importance for modulating social behavior, we also inspected the LgAG1 cluster (Figure 7A). AN05B023a, a GABAergic neuron downstream of LgAG1 (Figure 7G and 7H), and LgAG1 are each other’s top reciprocal synaptic partners. AN05B023a, in turn, inhibits PPN1 (AN05B102a), a neuron known to promote courtship ^84^. Since GR32a+ LgAG1 neurons have been shown to suppress interspecies and male-male courtship ^2,82^, we propose that they do so by inhibiting PPN1 (Figure 7H). Surprisingly, male-female courtship-promoting, fru+/Ir52a+ WG1 and fru-/Ir52a+/Ir52b+ LgLG2 GRNs are also strong upstream partners of AN05B023a (Figure 7G and 7H). Dissecting how AN05B023a integrates pheromone inputs from different GRN types will be key to understanding its role in controlling courtship behavior.

To sum up, our UMAP analysis confirms our initial clustering and expands our initial functional cell typing, allowing us to predict the valence of the recognized stimuli. Our analysis also suggests that the activity of GRN types from different body parts is integrated by common downstream partners based on valence (attractive vs. aversive) to control and coordinate specific behavioral outputs (feeding, sexual behavior, ingestion). By mapping key second-order neurons downstream of different functional clusters to different actions such as feeding aversion and courtship inhibition, we uncovered circuit architectures that can integrate inputs from different GRN types to coordinate flexible actions. This creates strong predictions that can be directly tested with the available genetic tools.

### Characterization of SEZ motor outputs from the GRN cell types

While the UMAP approach allows us to make broad predictions of the functional relevance of GRNs on behavior, these are inferred from the known function of specific interneurons. A detailed understanding of behavior requires the detailed mapping of the sensory-motor arc from sensory neurons to unique motor neurons innervating specific muscles. One of the main roles of gustatory systems is to guide feeding behavior. Specific motor neurons (MNs) mediate stereotyped motor responses upon the activation of the corresponding sensory system, which, when appropriately coordinated, lead to specific behavioral outputs. Similar to almost all GRN subclasses, all MNs mediating the feeding behavior of the fly are also localized in the SEZ, making this neuropil the key brain region which implements feeding-relevant sensorimotor transformations ^89–91^. To link GRN cell types to different behavioral outputs, we identified and cell typed all SEZ motor neurons in the male CNS and FAFB – FlyWire datasets.

The feeding MNs have their cell bodies located in the brain, and exhibit characteristic branching patterns in the SEZ ^21,24,25^. Their axons exit the brain via one of the two ipsilateral mouthpart nerves, the PhN or the LbN. To identify the neurons in the male CNS dataset (Figure 8A), we searched for morphological matches to all the MNs exiting through the PhN and the LbN in FAFB – FlyWire (Figure 8B). Whenever a match was not found, we used the previously described seed plane strategy to find and proofread the MNs. Validating our identification in the male CNS, we found matching cell counts for the MNs exiting via the LbN between hemispheres and to FAFB – FlyWire (Figure 8C). For the MNs exiting the PhN, we observed differences across the right and left sides within the male CNS, similarly to what was annotated in FAFB – FlyWire, and between the two connectomes. This might either correspond to inter-individual variabilities or challenges in the reconstruction and identification of the missing neurons.

**Figure 8.**
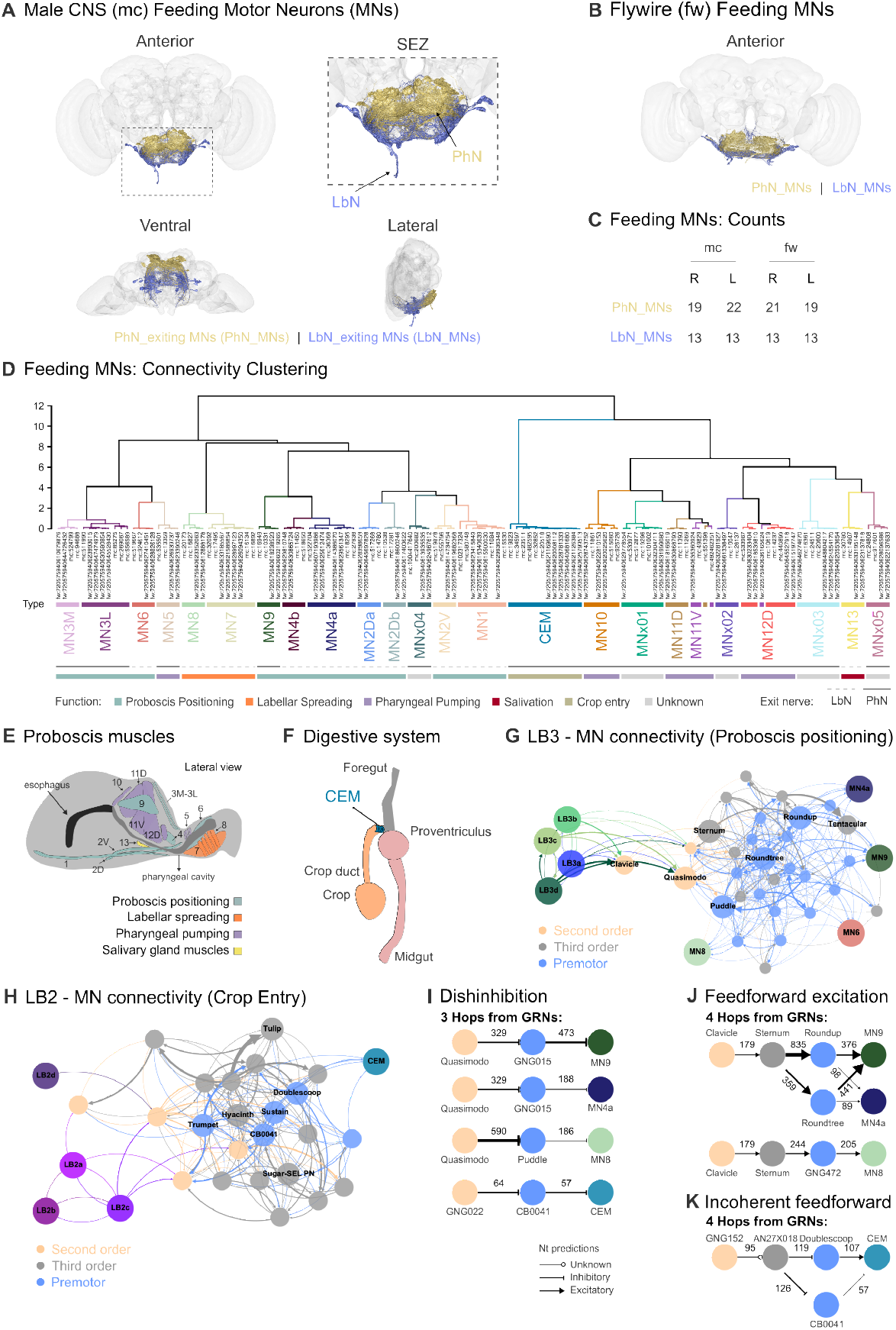
Characterization of the feeding motor outputs. **A**. Rendering of the feeding motor neurons in the male CNS in three different views, anterior, ventral, and lateral, colored according to their exiting nerve: labial (LbN) or pharyngeal (PhN) nerve. The SEZ is highlighted to showcase the different projection fields of the two neuronal populations. **B**. Rendering of the feeding motor neurons in FAFB – FlyWire in an anterior view. **C**. PhN- or LbN-exiting MN counts per side reveal comparable numbers between the two connectomes. **D**. Hierarchical co-clustering of the male CNS and FAFB – FlyWire feeding MNs based on inputs using cosine similarity and a threshold of 5 synapses. Male and female MN types (Figures S9A and S9B) cluster mostly together. When known, the MN type’s general function is indicated as well as its exiting nerve. **E**. Schematic, lateral view of the fly’s proboscis muscles, colored according to their general role, which can be controlling the proboscis’ positioning (blue), to either extend or retract it, mediating labellar spreading (orange), pharyngeal pumping (purple), or controlling salivation by innervating the salivary gland muscles (yellow). **F**. Schematic representation of the anterior digestive system with the location of the muscles at the crop entrance that are innervated by CEM. **G**. The sensorimotor pathway connecting attractive lbGRN subtypes, LB3a-d, to feeding motor neurons controlling different steps of proboscis extension (MNs 9, 4a, 6 and 8). The pathway is reconstructed by extracting cell types with high effective connectivity scores with both GRNs and MNs (>80th quantile) at each hop. Each circle is a cell type. Second-order neurons are colored light orange. Premotor neurons are colored light blue. If a downstream cell type has been previously studied or a driver line exists, its name is indicated. **H**. Same as G for the sensorimotor pathway connecting LB2 subtypes to crop entry motor neurons (CEMs). **I**. Disinhibition of feeding MNs by GABAergic second-order neurons, Quasimodo and GNG022. Whether a neuron is inhibitory/excitatory is derived from neurotransmitter predictions, as in Figure 7. Absolute synaptic weights are given. **J**. Same as I for feedforward excitation of feeding MNs. Clavicle is a key second-order neuron mediating feedforward excitation. **K**. Incoherent feedforward loop connecting LB2s to CEM. GNG152 is a key second-order neuron, with low confidence neurotransmitter predictions (unknown sign for connection).

Similar to the strategy used for the GRNs, we used connectivity clustering based on cosine similarity of the MN inputs to define different feeding MN cell types. We furthermore visually inspected and compared the corresponding morphologies of the neurons in each cluster to validate that they were correctly matched. The resulting separate connectivity clustering of male CNS (Figure S9A) and FAFB – FlyWire feeding MNs (Figure S9B) allowed us to identify 24 different cell types. Importantly, the co-clustering of the MNs from both connectomes revealed the same 24 types, with the corresponding male and female neurons clustering together (Figure 8D). Only one male CNS MN, typed MN11D, and one FAFB – FlyWire MN, annotated as MN11V, did not cluster with their types across datasets. In the case of the male CNS MN, this is very likely due to an incomplete reconstruction of the neuron.

To assign the identified MN cell types to different muscles and therefore functional classes, we took advantage of detailed previous work done on feeding-MN identification and characterization at the light microscopy level ^21,22,24,25,92^. Most feeding MNs innervate muscles in the proboscis and are, thus, globally known as proboscis MNs. The fly’s proboscis is a segmented appendage formed by the labellum, haustellum, and rostrum (Figure 8E), which are individually controlled and sequentially moved through specific joints to contact food ^21,24,25^. Feeding MNs can be categorized into five groups depending on their overall role (Figures 8E, 8F and S10). (i) Proboscis MNs 1, 2D, 2V, 3M, 3L, 4, 6, and 9 have been shown to control different steps of proboscis positioning, which includes the extension of its different segments, upon stimulation of appetitive gustatory GRNs (Figures 8E and S10). (ii) Proboscis MNs 7 and 8 mediate labellar opening/spreading in response to an attractive stimulus (Figures 8E and S10). (iii) Proboscis MNs 11D and 12D have been shown to control pharyngeal pumping ^23,49,93^, allowing the food to reach the esophagus. Given that they target muscles around the pharynx, MNs 5, 10, 11V, and 12V have been proposed to have similar functions in pharyngeal pumping (Figures 8E and S10). (iv) Proboscis MN13 is predicted to play a role in controlling salivation as the muscle it innervates inserts at the junction of the salivary duct with the pharynx ^25^ (Figures 8E and S10). MN12V ^25^ was the only proboscis MN for which no anatomical match was found in either male CNS or FAFB – FlyWire. It is also important to note that we also identified MN2D- and MN4-like neurons, which we have typed as MN2Db and MN4b, respectively. These types have similar upstream connectivity to the originally identified MNs 2D and 4, annotated, respectively, as MN2Da and MN4a, and morphologies similar enough to resemble those neurons but sufficiently different to be their own type (Figures 8D and S9C). (v) Another feeding MN, named CEM, which innervates muscles controlling crop entrance, has been shown to have a role in the postingestive decision to let food pass to the crop or not ^92^ (Figure 8F). Previously identified in FAFB – FlyWire, we also found CEM in the male CNS dataset (Figures 8D and S9C). Furthermore, in both datasets, we identified 5 MN types, named MNx01-05 (Figure 8D), that, to our knowledge, have never been described in the literature. Anatomically, each MN cell type is distinct, but identifiable across connectomes (Figure S9C). Some project ipsilaterally like MN2Da, MN7, MNx04, and MNx05, while most are bilateral. In general, as previously reported, SEZ MNs are either one or two right-left pairs of neurons (Figure S9D).

This functional specialization is also reflected in the connectivity clustering, revealed by three main clusters (Figure 8D). The first cluster is formed by MNs that are involved in the movement of the proboscis parts, such as positioning of the rostrum and haustellum, as well as spreading of the labellum ^21,24,25^. The second cluster is mainly formed by MNs involved in pharyngeal pumping ^21,25^, and the third cluster is formed by the CEM neurons controlling crop entry ^92^. This suggests that MNs form functional units, each of which is controlled by common premotor circuits to coordinate proboscis positioning and ingestion.

### Characterization of Sensorimotor Pathways Controlling Feeding Motor Actions

Because the EM datasets contain all GRNs, feeding MNs, descending interneurons controlling locomotion, and other relevant neurons in the feeding circuits, we are uniquely positioned to map how sensory information is transformed into actions. Taste is especially well-suited for this analysis, as taste-evoked behaviors are largely hard-wired. To do so, we used a strategy relying on calculating the effective connectivity scores between different GRN–effector neuron (e.g. MN) type pairs (see Methods) ^94^. Effective connectivity is an inferred estimate of how strongly activity in a given GRN type could influence an MN, considering not only direct synaptic connections but also indirect routes through intervening neurons. Importantly, it normalizes synaptic weights by the total input into target neurons at each layer. In this sense, it is analogous to evaluating synaptic drive in a physiological experiment: strong direct inputs contribute heavily, while longer multi-synaptic paths (numbered as hops) contribute progressively less. The metric thus captures the cumulative potential impact of a sensory input on a motor output across the polysynaptic network. As the maximum number of neurons a GRN can reach over multiple hops plateaus at 6-7 hops, we started by calculating the effective connectivity of each proboscis GRN-MN pair for up to 7 hops (Figures S11 and S12). High effective connectivity scores indicate that GRNs strongly influence specific MNs. Hierarchical clustering of effective connective scores between lbGRN-MN pairs led to two main clusters of lbGRNs in the male CNS dataset (Figure S11A). The first cluster is formed by the attractive lbGRNs, LB3a-d, and the aversive LB1a. These GRNs have relatively high effective connectivity scores with MN types that are involved in proboscis positioning and labellar spreading. This observation is consistent with the large body of knowledge showing that activation of phagostimulatory lbGRNs such as GR64e is sufficient to trigger proboscis extension. Interestingly, the second cluster, which is formed by LB1b, c, d, e, LB2s, and LB4s, has very low effective connectivity scores with proboscis positioning MNs. In contrast, these neurons are predicted to be strongly connected to MNs that control pharyngeal pumping and crop entry. As our cell typing and the UMAP analysis suggest that LB2s mediate aversion and LB4s mediate attraction, we propose that LB2s do so by suppressing the ingestion of aversive compounds while LB4s promote ingestion. LB2s have particularly high effective connectivity scores with CEM neurons, which have been suggested to control food passage in the fly digestive system ^92^. We therefore propose that the control of food passage in the digestive system might be an important function of LB2s. Importantly, clustering effective connectivity scores using the FAFB – FlyWire dataset led to similar results, with only small differences in some subclusters (Figure S11A).

Another way to dissect how GRNs affect feeding behavior is to compute which MNs they reach with the least number of hops. We therefore computed the effective connectivity scores between GRN – MN pairs at each hop and ranked them according to which reached the corresponding MNs with the least number of hops (Figure S11B). We focused on MNs that are key for controlling different steps of feeding: MN9 (rostrum protraction), MN4a (haustellum extension), MN6 (labellar extension), MN8 (labellar abduction/spreading), MN11D or V (pharyngeal pumping), and CEM (crop entry). LB3a-c (sugar and other phagostimulatory GRNs) and LB1a (bitter GRN) reach maximum effective connectivity scores with proboscis positioning and labellar spreading MNs after 4-5 hops (Figure S11B). In addition, they show a sharp increase in effective connectivity score already after 2-3 hops. While it is at first sight counterintuitive that bitter and sugar GRNs are predicted to reach proboscis positioning MNs with the same effectiveness, this agrees with the previous findings that aversive GRNs mediate aversion via suppressing feeding-promoting sensorimotor pathways and not by acting on other MNs, such as the ones mediating proboscis retraction ^47^. Similar to what we observed in the previous analysis, LB2 GRNs reach high effective connectivity scores with pharyngeal pumping MNs and CEMs in fewer hops than other GRN types (Figure S11B). This further suggests that they are important for modulating ingestive and postingestive behaviors. These findings suggest that proboscis extension and ingestion are mediated by a multilayered circuitry, which allows for the implementation of complex feedforward and feedback regulatory mechanisms. We applied the same approach to tpGRNs and phGRNs (Figure S12). As we report in our companion paper, dtpGRNs have strong connections with pharyngeal pumping MNs as well as the CEM neurons ^49^. ctpGRNs exhibit interesting dimorphisms between the two datasets (Figure S12A). While they are strongly connected to proboscis positioning MNs in the male dataset, they are more strongly connected to pharyngeal pumping MNs in the female dataset. As we identified genetic driver lines labeling this GRN type, further functional dissection will clarify the functional relevance of this potential sexual dimorphism. Interestingly, ctpGRNs reach high effective connectivity scores with proboscis positioning MNs in fewer hops when compared to dtpGRNs (Figure S12B). dtpGRNs and ctpGRNs have similar profiles for MN11V. However, dtpGRNs reach CEMs in a few hops and have comparably high effective connectivity scores with these MNs. phGRNs seem to be poorly connected to proboscis positioning MNs except for PhG2 (Figure S12C). In general, phGRNs are strongly connected to pharyngeal pumping MNs and CEM. This supports the idea that the main function of phGRNs is to control food passage to the digestive system and to therefore, serve as gatekeepers of the body. Something, which is not surprising given their anatomical location.

As stated, PhG2 reaches high effective connectivity scores with proboscis positioning and pharyngeal pumping MNs after 3-4 hops (Figure S12D). Most phGRNs reach high effective connectivity scores with MN11V in a few hops, with PhG2 having the highest score. As previously mentioned, PhG2 is likely to inhibit proboscis extension and ingestion by suppressing different MNs (Figure 7E), explaining why it is connected to all feeding-relevant MNs. Finally, PhG3, PhG4, PhG6, and PhG7 reach high effective connectivity scores with CEM in only 4 hops (Figure S12D). As we proposed earlier, PhG3 and 4 are likely to be ppk28+, water-sensing phGRNs. This observation suggests that water entry to the crop is tightly regulated by these phGRNs. Intriguingly, PhG12-14 and PhG16 seem to be weakly connected to any feeding or ingestion MN when compared to other phGRNs in the male CNS (Figure S12C and S12D), suggesting that they might instruct other behaviors like learning. In FAFB – FlyWire, PhG15 also joins this cluster, while being strongly connected to CEM in male CNS (Figure S12C).

To reconstruct a minimal sensorimotor circuit that connects lbGRNs LB3a-d to proboscis positioning motor neurons (MNs 9, 6, 4 and 8), as well as LB2s to CEM, we selected neurons with high effective connectivity scores with both GRNs and MNs at each hop (Figure 8G and 8H; see Methods). This analysis revealed key second-order, higher-order, and premotor neuron types, some of which have already been identified as controlling feeding in other studies ^49^, suggesting that our approach extracts important circuit features underlying behavior. Careful dissection of these sensorimotor pathways revealed two distinct circuit motifs which could be observed multiple times across the feeding circuit (Figure 8I and 8J). First, we observed a three-hop motif which leads to the disinhibition of different MNs controlling proboscis positioning and crop entry (Figure 8I). In one example, the activation of the GABAergic second-order neuron Quasimodo by phagostimulatory GRNs leads to the disinhibition of MNs 9, 4a, and 8, therefore enabling them to get activated (Figure 8I). Second, we observed a longer, four-hop feedforward excitation pathway mediated by the key cholinergic second-order neuron Clavicle (ANXXX462), which should lead to the activation of the same MNs (Figure 8J). We propose that the temporal dynamics produced by the combination of three-hop disinhibition and four-hop feedforward excitation might allow for the generation of complex rhythmic proboscis extension and retraction cycles, such as observed when the fly feeds on solid food ^95^. We furthermore observed a four-hop incoherent feedforward circuit controlling the CEM neurons via the cholinergic Doublescoop and glutamatergic CB0041 neurons (Figure 8K). As genetic driver lines labeling these cell types exist ^96^, it will be possible to dissect the circuit dynamics controlling crop entry by labellar bristle inputs.

Taken together, our analysis suggests that each step of the motor sequence underlying fly feeding is controlled by a group of specific GRNs that are distributed across different taste sensilla.

### Feedforward Control of Neuromodulation by GRNs

One of the most striking features of taste is its ability to modulate peripheral physiology before nutrients are absorbed into the body. In mammals, this is exemplified by the cephalic phase insulin response (CPIR), where food-related sensory signals activate parasympathetic pathways to trigger insulin secretion in advance of food digestion. Beyond insulin, food perception is sufficient to modulate the activity of brain regions that regulate satiety states ^97–99^ and to impact peripheral metabolism, including hepatic homeostasis ^100^, lipolysis ^101^, and adipose tissue thermogenesis ^102^. We therefore asked to what extent GRNs are wired into neuroendocrine circuits that prepare the organism for metabolic challenges. Given that neurosecretory cells (NSCs) that produce various hormones have been identified in the FAFB – FlyWire and male CNS datasets ^60,103^, we computed effective connectivity scores between male CNS GRNs and NSCs predicted to express insulin (labeled as IPCs), hugin (labeled as Hugin neurons), ion transport peptide (labeled as ITP neurons), CAPA (labeled as CAPA neurons), diuretic hormone 31 (labeled as DH31 neurons), diuretic hormone 44 (labeled as DH44 neurons), corazonin (labeled as CRZ neurons), or myosuppressin (labeled as DMS neurons) (Figure S13A). One cluster comprising most phGRN types, dtpGRNs, and the labellar LB2 and LB4 types had particularly high connectivity scores with DMS, DH44, and IPC neurosecretory neurons (Figure S13A). Interestingly, these GRNs also have high effective connectivity with MNs that control pharyngeal pumping and ingestion (Figures S11 and S12). Other GRNs, especially lgGRNs and wGRNs, showed generally low effective connectivity with NSCs (Figure S13A). This makes sense, as simple food sampling via leg and wing bristles, without ingestion, should not lead to an anticipatory remodeling of animal physiology. Similarly, LgAG1 neurons, which we predict are involved in sexual behavior, are also in this cluster. Our findings suggest that taste perception and neuronal signaling at early stages of ingestion might prime metabolic programs for satiety, like in mammals ^104^. To start dissecting the neuronal circuits doing so, we focused on the connectivity of putative sugar-sensing PhG1s with insulin-secreting IPCs. Our circuit reconstruction shows that PhG1a-c connect to IPC in three hops through a layered, convergent feed-forward relay (Figure S13B). Interestingly, CB0041 and the Sustain neuron, which are involved in controlling the length of yeast feeding bursts ^49^, connect phGRN types to IPCs. The involvement of Sustain and CB0041 suggests that the same interneurons that control feeding burst length may also couple ingestion to IPC activity, linking microstructure of feeding with systemic metabolic regulation.

### Control of Proboscis Motor Neurons by Leg and Wing GRNs

Foraging flies detect food first using their bristle sensilla on each of the six legs before they extend their proboscis, showing that food detection by leg GRNs is sufficient to trigger proboscis extension ^19^. To identify candidate circuits that transform appetitive stimulation of appendages into proboscis movements, we decided to compute the effective connectivity scores between leg and wing bristle GRNs and feeding motor neurons. We normalized the scores by the total leg/wing GRN scores to highlight the relative connection of each GRN type to each MN (Figure 9A). Our analysis showed that wGRNs and lgGRNs are poorly connected to feeding MNs except for the LgLG4 type, which is included in the cluster that is more strongly connected to feeding MNs. The GRN types we proposed to be appetitive, LgAG2 and LgLG4 (Figures 5E, 6H and 7A), are clustered together (Figure 9A), suggesting that their activity is integrated to generate proboscis extension response to appetitive stimuli. To predict which leg GRN types exert the most direct influence on feeding motor neurons, we calculated effective connectivity scores and identified those reaching the highest scores with distinct feeding motor neurons in the fewest hops (Figure S14A). Among them, LgLG4 (which we predict to express the sugar receptor GR64f) stands out, reaching high connectivity scores after few hops with each of the motor neurons controlling the entire feeding motor sequence: MN9 (rostrum protraction), MN4a (haustellum extension), MN6 (labellar extension), MN8 (labellar spread), MN11D (pharyngeal pumping), and CEM (crop entry). On average, however, LgLG4 requires 7 hops to reach its maximum effective connectivity score with feeding MNs, two more than most labellar GRNs. This fits with the well-known observation that labellar gustatory stimulation is more efficient in eliciting proboscis extension than leg stimulation ^105^.

**Figure 9.**
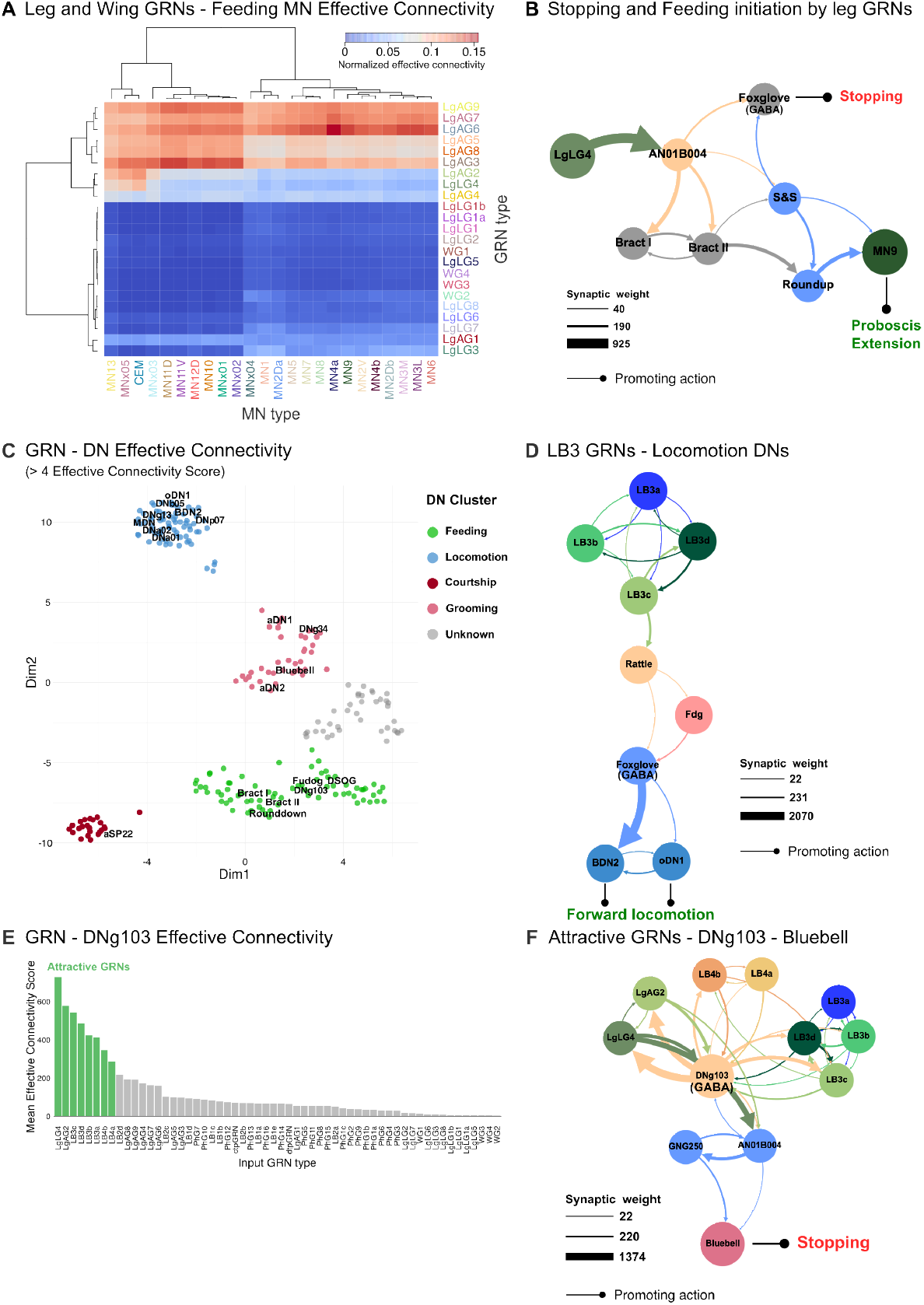
Control of different motor actions by different types of GRNs via ascending and descending pathways. **A**. Heatmap for effective connectivity from leg and wing GRNs to feeding motor neurons. Effective connectivity scores were normalized by the total effective connectivity to each MN. For effective connectivity calculations, a maximum of seven hops was used. Ward’s method was used for hierarchical clustering and building the dendrograms. **B**. LgLG4s downstream pathway mediates stopping and proboscis extension. If a downstream cell type has been previously studied or a driver line exists, the name from the literature is used. Otherwise, male CNS cell types are used. Each circle is a cell type. Second-order neurons are colored light orange. Premotor neurons are colored light blue. Absolute synaptic weights are given. **C**. UMAP embedding of DNs based on their effective connectivity scores with GRN types. Only DNs that have >4 effective connectivity score are shown. Each circle is a DN cell type, and each cluster is color-coded. DNs studied previously and in this study are labeled. **D**. Attractive lbGRNs control locomotion. If a downstream cell type has been previously studied or a driver line exists, the name in the literature is used. Otherwise, male CNS cell types are used. Each circle is a cell type. Absolute synaptic weights are given. BD2 and oDN1 are two DN types that have been shown to be involved in forward locomotion ^26,88^. **E**. GRN – DNg103 effective connectivity. GRNs in the attractive cluster (Figure 7) are highlighted in green. **F**. Attractive GRNs and GABAergic DNg103 form a negative feedback loop to control Bluebell, whose activation leads to stopping ^26,88^, via AN01B004 and GNG250. Each circle is a cell type. Absolute synaptic weights are given. If a downstream cell type has been previously studied or a driver line exists, the name in the literature is used. Absolute synaptic weights are given.

Careful analysis of the LgLG4 downstream circuits revealed that LgLG4 connects to Bract neurons in two hops via the ascending neuron AN01B004 (Figure 9B). Bract neurons are also downstream of labellar sugar-sensing GRNs, and their optogenetic activation is sufficient to trigger proboscis extension ^47^. Bract II connects to MN9 via two premotor neurons, Sink and Synch (S&S) ^96^ and Roundup ^47^. S&S, predicted to be cholinergic, receives direct inputs from AN01B004 and outputs directly onto Roundup. Together, this circuit forms a sensory-motor arc connecting LgLG4 to MN9 through multiple positive feedforward loops.

Beyond proboscis extension, LgLG4 also engages downstream pathways that coordinate feeding with locomotor suppression and ingestion. Feeding and forward locomotion are mutually exclusive ^19^. Flies feeding on a food patch suppress forward locomotion while still exhibiting micromovements around the patch ^106,107^. Consistent with this, LgLG4 connects to Foxglove, a GABAergic neuron that suppresses forward walking ^88^, both directly via AN01B004 and indirectly via the S&S neuron (Figure 9B). In addition, another ascending neuron, AN05B106, connects LgLG4 GRNs to the pharyngeal pumping motor neuron MN11D (Figure S14B) and crop-entry motor neuron CEM (Figure S14C). We therefore propose that the LgLG4 pathway exemplifies how sensory input can be wired into circuits that simultaneously inhibit competing behaviors and promote feeding, thereby enforcing the behavioral switch from exploration to exploitation.

### Descending control of motor actions by GRNs

Descending neurons (DNs) are information bottlenecks that connect the brain with the VNC. They integrate sensory information and behavioral states, and orchestrate various complex actions such as locomotion, courtship, and grooming ^108^. As aforementioned, gustatory signals are important modulators of locomotion during foraging and sexual behavior. Furthermore, the detection of pathogenic bacteria triggers hygienic grooming via the activation of taste bristles on the anterior wing margin ^109^.

To understand how descending circuits integrate gustatory inputs, we performed a UMAP embedding of DN cell types based on their effective connectivity with different GRN types (Figure 9C). This analysis revealed five clusters, which we annotated by identifying functionally characterized DNs within them (Figure 9C). Strikingly, DNs known to control similar motor actions clustered together: for example, most locomotion-related DNs ^108^ were part of a “locomotion cluster”; DNs that promote or suppress feeding are clustered in a “feeding cluster”; and the courtship-promoting DN aSP22^110^ is found in a separate “courtship cluster”. One cluster, however, lacked functionally characterized members and therefore remains of unknown function. These results indicate that gustatory information is selectively routed into descending pathways that control distinct behavioral domains, providing a circuit framework through which taste can influence diverse motor programs.

As mentioned earlier, forward locomotion is suppressed during feeding. We therefore asked whether we could find evidence that appetitive proboscis GRNs contribute to this phenomenon. Our analysis revealed that LB3c, an appetitive lbGRN subtype, connects to two forward locomotion-promoting DNs, BDN2 and oDN126, through a circuit involving the interneurons Rattle, Fdg, and Foxglove (Figure 9D). Within this circuit, Rattle and Fdg form a positive feed-forward loop that activates Foxglove. Foxglove, in turn, inhibits both BDN2 and oDN1, thereby suppressing forward locomotion ^26^. This identifies a direct neural pathway by which appetitive proboscis input could suppress locomotion, ensuring that the detection of food reliably promotes feeding over exploration.

A particular descending neuron, DNg103, caught our attention because it shows very high effective connectivity scores with all attractive GRN types identified in this study (Figures 9E and S7B). DNg103 is predicted to be a GABAergic descending neuron that receives direct inputs from LgLG4, LgAG2, all LB3 and LB4 subtypes, with LgLG4 as its strongest GRN partner. Strikingly, DNg103 sends strong inhibitory feedback connections onto these attractive GRN types (Figure 9F). Such negative feedback loops are known to be sufficient to generate oscillatory dynamics, such as in transcriptional circadian oscillations ^111^. Downstream, DNg103 connects to Bluebell, a neuron that induces stopping ^26^, via cholinergic AN01B004 and GABAergic GNG250 (Figure 9F). We therefore speculate that the reciprocal negative feed-back between attractive GRNs and DNg103 could generate toggle switch/oscillatory activity in DNg103, which, in turn, would lead to stopping-walking sequences characteristic of feeding-related micromovements ^106^. Such dynamics may provide a circuit mechanism for the micro-exploration of food patches that occurs once feeding is initiated.

Finally, we propose a novel role for ctpGRNs during courtship. Among all GRNs, ctpGRNs have the highest effective connectivity score with the courtship-promoting DN aSP22 (Figure S15A). Our analysis revealed that ctpGRN connects to aSP22 in 3 hops through multiple pathways, suggesting that IR56d-positive ctpGRN (Figure 3G) could contribute to the modulation of courtship (Figure S15B). As ctp-GRNs are located between the pseudotrachea within the labellum, we speculate that they are likely to contribute to the courtship motor sequence triggered when males lick the genitalia of females.

Altogether, we used state-of-the-art connectomics to reconstruct and cell type the entire gustatory connectome and feeding motor neurons of *Drosophila melanogaster* in male CNS, MANC, and FAFB – FlyWire datasets (Figures 1 and 8). We combined morphological comparisons, synaptic connectivity, and neurotransmitter predictions to identify unique GRN types and match them with receptor expression patterns, allowing us to predict the tastants they detect (Figures 2-6 and 10A). UMAP embedding of downstream connectivity revealed novel attractive and aversive GRN types (Figures 7A and 10A). By analyzing effective connectivity between GRNs and effector neurons, we uncovered key higher-order neurons and circuit motifs that might transform gustatory information into diverse outputs, such as neurosecretion, halting, proboscis extension, swallowing, ingestion, and courtship (Figures 8, 9, and 10B). Together, our work lays out a detailed gustatory sensorimotor connectome across multiple EM datasets, establishing a roadmap and hypothesis-generating platform to functionally dissect the neural circuit logic of conserved, state-dependent behavioral motor sequences such as feeding and mating.

**Figure 10.**
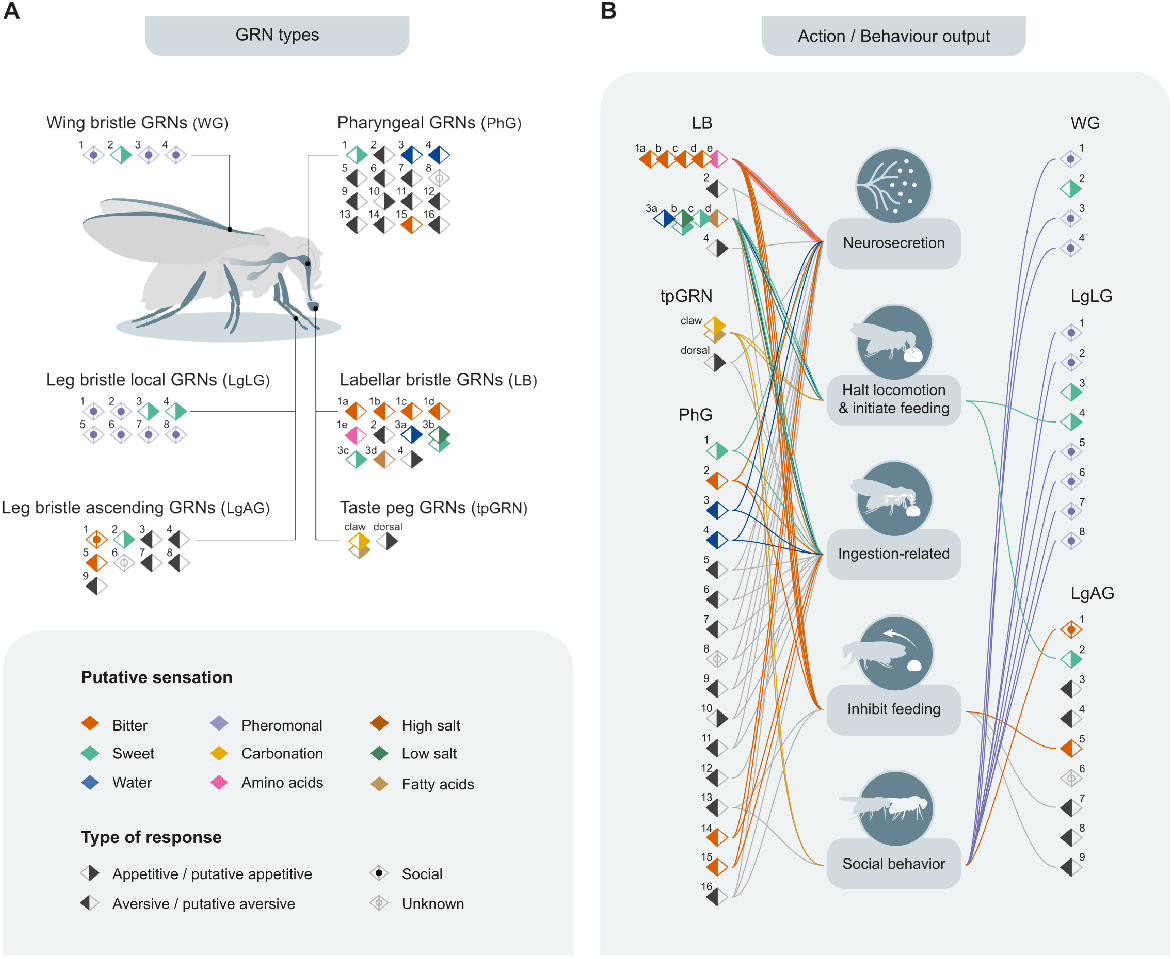
Summary of the fly gustatory connectome and its actions. **A**. Schematic summary of different GRN types across different body parts and their sensory profiles based on our predictions. The type of responses (appetitive, aversive, or social) which we predict to be controlled by these GRN types are also shown. Colors indicate the putative sensation, and different shapes indicate the predicted responses mediated by different GRN types. **B**. Actions and behavioral outputs mediated by different GRN types. Lines connect GRN types to actions we predict they mediate. Most GRN types have a predicted sensation, valence (response), and action. Social behavior comprises sexual behaviors.

## Discussion

In his seminal work, “The Hungry Fly” ^19^, Vincent D. Dethier meticulously described the gustatory sensilla of the black blow fly, its feeding system, and its behavior and noted: “All of the intricate behavior patterns described here suggest that a study of the relation between sensory input and motor output would be a rewarding endeavor.” Advances in molecular genetics and histological methods in *Drosophila melanogaster* allowed the field to pursue his vision, accelerated by the identification of taste receptors and followed by systematic physiological, anatomical, and behavioral studies that dissected the neural circuit mechanisms underlying feeding ^1,6^. Advances in EM connectomics in flies ^13,14,30,31^ now provide a unique opportunity to map the entire gustatory sensorimotor connectome with unprecedented detail. Work in fly larvae has already demonstrated the power of this approach, revealing how specific gustatory inputs are wired to motor and neuroendocrine outputs ^89^. In adults, however, circuit knowledge had remained fragmentary. Previous studies have provided important insights into labellar bristle GRNs, including some morphological types, and their functional connection to the motor neuron 9^47,112^, but a comprehensive overview of gustatory circuits across body parts and motor outputs has been lacking.

### Cell typing of GRNs

Here, we mapped the complete gustatory connectome of the fruit fly across three EM datasets: FAFB – FlyWire (female brain) ^14,30,31^, MANC (male ventral nerve cord) ^32,33^, and male CNS (entire male central nervous system) ^60^. The male CNS volume in particular enabled us to identify and annotate the full repertoire of gustatory receptor neurons in a single dataset (Figure 10). By combining neuroanatomical comparisons with synaptic connectivity and a consensus cell type atlas, we were able to fine-tune the annotation of GRN subtypes ^60,94^, allowing us to further refine the GRN types that recur across independent datasets. This cross-dataset reproducibility strongly supports our celltyping rather than the different subclasses being due to individual variability. This approach revealed a remarkably diverse gustatory system, comprising four labellar bristle GRN types (lbGRNs; with a total of 15 subtypes), two taste peg GRN types (tpGRNs), 16 pharyngeal GRN types (phGRNs), nine directly ascending leg bristle GRN types (lgAGRNs), four wing bristle GRN types (wGRNs), and eight locally projecting leg bristle GRN types (lgLGRNs).

Notably, most pharyngeal GRNs are just a single pair of neurons, each projecting to distinct subsets of downstream circuits. This organization may provide a form of spatial discrimination within the pharynx, where different ingestion checkpoints are monitored by dedicated circuit units. A similar logic has been proposed for the expansion of cell types in the dorsal root ganglia in vertebrates, where increased cellular diversity could enhance the spatial resolution of somatosensory inputs ^113,114^.

By matching the projection patterns of these GRNs to reporter lines, we could identify many of the previously described receptor-GAL4 cell types, adding further validity to our classification. In addition, we identified several GRN types that could not be matched to known driver lines, suggesting the existence of novel, yet unexplored GRN types. Moreover, five aversive GRN classes that had been defined in classic electrophysiological and functional imaging studies reappear in our connectomic analysis, providing an independent line of support for our annotations ^12,62^. In a companion study ^49^, we further functionally validated part of our cell typing by showing that taste peg GRNs differ in their function, and that only one type (dtpGRNs) is necessary to regulate the length of yeast feeding bursts. This contrasts with the classic view that tpGRNs are one homogeneous group. Together, these lines of evidence support the robustness of our cell typing and reveal the strength of defining cell types using morphology and connectivity data across EM datasets. We propose that GRN types have distinct but overlapping roles in a wide range of biological functions, including foraging, feeding, sexual behavior, and neuromodulation (Figure 10).

### Valence and Modularity

Our study further revealed how GRN diversity maps onto valence. UMAP embedding of GRNs based on downstream connectivity revealed that previously characterized attractive and aversive GRN types segregate into distinct clusters. These clusters also contain previously uncharacterized GRN types, suggesting that they likely encode the same valence as other GRNs in these clusters. Notably, both attractive and aversive clusters contained GRN types from different body parts, including legs, labellum, and pharynx, indicating that valence information from distributed gustatory sensilla on different sensory organs is integrated to generate unified perceptual decisions. At the same time, distinct sensorimotor pathways downstream of each GRN type would preserve compound-specific information to orchestrate the appropriate motor programs (Figure 10A and 10B). We propose that this modular organization, integrated valence coding combined with pathway-specific motor control, enables flexible adaptation of feeding behavior to changing external conditions and internal states.

### Circuit motifs and functional validation

To move beyond direct synaptic partners, we used an effective connectivity metric that quantifies the relative strength of pathways linking GRNs to motor neurons across multiple synaptic hops ^94^. This approach collapses the complexity of many possible routes into a normalized measure of influence, allowing us to identify the pathways most likely to be functionally relevant. Crucially, because we first identified and annotated the complete set of proboscis movement and ingestion motor neurons, we could bridge GRN inputs to specific motor outputs, rather than relying on other proxies. We identified key second-order, higher-order, and premotor neurons that receive inputs from a diverse set of aversive GRNs. We confirmed the functional validity of this approach in an accompanying study, where we showed that dtpGRNs are directly connected to pharyngeal pumping MNs, and that this pathway is important for regulating the length of feeding bursts on yeast by modulating ingestion efficiency ^49^.

Applying this analysis revealed two recurrent motifs by which attractive GRNs can control proboscis movements. First, a fast disinhibition pathway gates access to motor neurons controlling proboscis positioning and labellar spreading. Such disinhibition might act as a thresholding mechanism, ensuring that the feeding motor program is only triggered when the stimulus is sufficiently strong. Second, a longer feedforward excitatory pathway recruits the same motor neurons with presumably slower dynamics, potentially coordinating their sustained activation during ongoing feeding (Figure 10A and 10B). We propose that this circuit architecture ensures avoiding the intake of harmful compounds by tonically inhibiting feeding at multiple circuit layers and potentially coordinating their sustained activation during ongoing feeding via feedforward excitation. It will be crucial to test these hypotheses with functional experiments and activity recording approaches.

Interestingly, recent vertebrate work points to an analogous logic. A study in mice describes a multi-hop disinhibition motif spanning the amygdala–parabrachial–brainstem pathway that gates licking premotor neurons and thereby regulates lick-bout duration ^115,116^. This suggests that recurrent disinhibition may represent a conserved circuit strategy for structuring ingestive bouts across species, from flies to mammals. More broadly, these findings highlight how motifs revealed by connectomics in the fly gustatory system can inform general principles of how sensory inputs are transformed into adaptive motor programs.

### Specialization and flexibility

Our analysis also shows that most labellar bristle GRNs are more strongly connected to MNs controlling proboscis positioning, whereas most pharyngeal GRNs are predicted to control ingestion-related MNs. This general division reflects their anatomical position in the feeding sequence. Yet, exceptions such as LB2s connecting strongly to crop entry and PhG2s to proboscis positioning MNs highlight that not all GRNs follow this stereotypy. Such exceptions suggest functional flexibility, with select GRN types potentially allowing cross-talk between different subroutines of feeding.

Interestingly, effective connectivity patterns were largely conserved across sexes, with one notable exception: ctp-GRNs seem to favor proboscis positioning over pharyngeal pumping MNs in the male, but the reverse is true in the female. Moreover, their connectivity to the courtship-promoting descending neuron aSP22 raises the possibility that ctpGRN might be involved in sexually dimorphic behaviors.

### Sequential checkpoints and action control

Gustatory sensilla are exposed to chemical stimuli in a defined sequence during foraging and feeding: GRNs on the legs touch food first, allowing the fly to probe the food and decide if the proboscis should be extended. Labellar bristles then assess food quality, and if appropriate, labellar spread exposes taste pegs. Finally, coordinated pharyngeal muscle contractions pump the food into the cibarium, activating pharyngeal GRNs. Meanwhile, the fly suppresses forward locomotion to continue eating on the same food patch. Each GRN subclass thus acts as a checkpoint for transitions between subsequent motor actions based on their anatomical locations. These transitions are state-dependent, as demonstrated in both flies and mammals, where nutrient deprivation enhances initiation probabilities at multiple nodes of the feeding sequence ^5^.

Calculating the effective connectivity between leg GRNs and proboscis motor neurons, we showed that the attractive leg GRN type LgLG4 is likely to control proboscis extension via an ascending neuron while suppressing forward locomotion through another interneuron. In turn, attractive labellar bristle GRNs can suppress locomotion by inhibiting forward locomotion DNs. In another study, we functionally characterized a sensorimotor circuit downstream of dtpGRNs that allows subsequent proboscis extensions by facilitating food swallowing ^49^. Together, these circuits illustrate how checkpoints are enforced and chained to ensure coherent action sequences, tuned by nutrient and mating states.

### Decision bottlenecks in descending pathways

Taste is also important for controlling other behaviors, such as social behaviors and hygienic grooming. Descending neurons (DNs) act as bottlenecks where sensory information converges to be transformed into coordinated actions. UMAP embedding of DNs based on their effective connectivity with GRN types revealed clusters likely specialized for different actions. We identified a second-order inhibitory DN, DNg103, connected to attractive GRNs in a negative feed-back loop. We speculate that this circuit generates oscillations, producing the characteristic intermittent stopping and moving “micromovements” in feeding flies via the stopping inducing DN, Bluebell. Comparable descending bottlenecks also exist in mammals, where gustatory signals in the brainstem converge onto reticulospinal and orofacial premotor neurons to implement licking and swallowing programs ^117,118^.

### Anticipatory control of physiological state

Sensory perception is an important determinant of internal state modulation ^119^. Here, we systematically explore how gustatory information in flies can reach neurosecretory cells via synaptic connections. Strikingly, only GRNs exposed to food upon ingestion or those tightly coupled to ingestion-related MNs are strongly connected to these endocrine outputs. This suggests a strategy to avoid undesired anticipatory changes in physiology based on noisy or irrelevant external cues, favoring signals linked to actual ingestion. Notably, putative sugar-sensing PhG1s connect to insulin-producing neurons, arguing that ingestion of caloric food can modify the satiety state in an anticipatory feedforward way. These mechanisms allow the fly to anticipatorily reprogram its physiology to prepare for the digestion of incoming food and switch from a catabolic to an anabolic state. This parallels cephalic-phase responses in mammals, where oropharyngeal inputs trigger endocrine release ^120,121^, though in flies, such modulation appears more tightly gated by ingestion checkpoints.

### Conclusion

Our work provides a comprehensive and detailed gustatory sensorimotor connectome across multiple EM datasets in flies, establishing a blueprint to functionally dissect the neural circuit logic of conserved, state-dependent behavioral motor sequences underlying foraging, feeding, and mating. The detailed cell typing and effective connectivity analysis outlined here generate testable hypotheses about the logic of feeding control, from sensory checkpoints to descending bottlenecks, neuroendocrine coupling, and sexual dimorphisms. The powerful neurogenetic tools available in *Drosophila* make it possible to follow up on these predictions at cellular resolution. We are now in a unique position to unravel neural mechanisms underlying complex, adaptive sensorimotor transformations underlying foraging and feeding behaviors.

## Consortia

The members of the Janelia FlyEM Project Team are Stuart Berg, Stephan Preibisch, Wei Qiu, Shin-ya Takemura, Michael Cook, Kenneth J Hayworth, Gary B Huang, William T Katz, Zhiyuan Lu, Christopher Ordish, Tyler Paterson, Eric T Trautman, Feng Li, Iris Ali, Brandon S Canino, Jody Clements, Samantha Finley-May, Miriam A Flynn, Imran Hameed, Gary Patrick Hopkins, Philip M Hubbard, Julie Kovalyak, Shirley A Lauchie, Meghan Leonard, Alanna Lohff, Charli A Maldonado, Caroline Mooney, Nneoma Okeoma, Donald J Olbris, Birava Patel, Emily M Phillips, Stephen M Plaza, Jennifer Rivas Salinas, Ashley L Scott, Louis A Scuderi, Claire Smith, Rob Svirskas, Satoko Takemura, Alexander Thomson, Lowell Umayam, John J Walsh, C Shan Xu, Emily A Yakal, Tansy Yang, Reed George, Vivek Jayaraman, Wyatt Korff, Geoffrey W Meissner, Jan Funke, Christopher Knecht, Louis K Scheffer, Gwyneth M Card, Harald F Hess, Gerald M Rubin and Gregory SXE Jefferis. The members of the Cambridge Connectomics Group based in the Department of Zoology, University of Cambridge and MRC Laboratory of Molecular Biology are Marta Costa, Isabella R Beckett, Philipp Schlegel, Elizabeth C Marin, Alexandra MC Fragniere, Andrew S Champion, Marina Gkantia, Florian Kämpf, Tomke Stürner, Catherine R Whittle, Billy J Morris, Markus W Pleijzier, Yijie Yin, Griffin Badalamente, Paul Brooks, Sebastian Cachero, Bhumpanya Chaisrisawatsuk, Márcia dos Santos, Christopher R Dunne, Katharina Eichler, Ladann Kiassat, Ilina Moitra, Sung Soo Moon, Eva J Munnelly, Anika Pai, Alana Richards, Holly Whittome and Gregory SXE Jefferis.

## Acknowledgments

We thank Ana Paula Elias for her help with brain dissections and immunostainings. We thank Gabrielle Sterne and Attilio P. Ceretti for sharing MN annotations in FlyWire, Tyler Sizemore, Hubert Amrein, and the whole Behaviour and Metabolism laboratory for many fruitful discussions, valuable feedback throughout the project, and comments on the manuscript. We thank the Champalimaud Fly and Rodent Platforms and the Champalimaud Advanced BioImaging and BioOptics Experimental Platform. Lines obtained from the Bloomington Drosophila Stock Center (NIH P40OD018537) were used in this study. I.T. was supported by a Marie Skłodowska-Curie postdoctoral fellowship (H2020-WF-01-2018-867459), R.B. was supported by a doctoral fellowship 2021.07285.BD from the Portuguese Foundation for Science and Technology (FCT). The project leading to these results has received funding from FCT (PTDC/MED-NEU/4001/2021), and the “la Caixa” Banking Foundation under the project codes HR17-00539 and HR23-00516 to C.R., Wellcome Trust Collaborative Award (220343/Z/20/Z) to G.S.X.E.J. with G. Rubin, G. Card, and S. Waddell, and an MRC Neuronex2 award (*MC*_*EX*_*MR/T* 046279*/*1) and core support from the MRC (MC-U105188491) to G.S.X.E.J., A FENS Kavli Alumni Scientific Exchange Award From the Kavli Foundation and the Federation of European Neuroscience Societies to C.R. and G.S.X.E.J.. Research at the Centre for the Unknown is supported by the Champalimaud Foundation and by Portuguese national funds through FCT - Fundação para a Ciência e a Tecnologia - in the context of the project UIDB/04443/2020, the research infrastructure CONGENTO, co-financed by Lisboa Regional Operational Programme (Lisboa2020), under the PORTUGAL 2020 Partnership Agreement, through the European Regional Development Fund (ERDF) and FCT under the project LISBOA-01-0145-FEDER-022170, and the PPBI - LISBOA-01-0145-FEDER-022122.

## Author contributions

Conceptualization, I.T., G.S.X.E.J. and C.R; methodology, I.T. and C.R.; investigation, I.T., I.V., R.J.B., B.J.M, I.B, P.S., E.C.M, M.C. and G.S.X.E.J.; formal analysis, I.T., I.V., R.J.B., B.J.M, I.B, P.S., E.C.M, M.C. and G.S.X.E.J.; visualization, I.T. and I.V.; software I.T., P.S. and G.S.X.E.J.; writing, I.T., I.V. and C.R.; supervision, I.T., E.C.M., M.C., G.S.X.E.J. and C.R.; funding acquisition, G.S.X.E.J. and C.R.

## Resource availability

### Lead contact

Requests for further information and resources should be directed to and will be fulfilled by the lead contact, Carlos Ribeiro.

### Materials availability

This study did not generate new, unique reagents.

### Data and code availability

The body IDs of GRN and MN cell types are shared in Supplementary Table 2. The Male CNS connectome dataset is publicly available (https://male-cns.janelia.org/). The FAFB – FlyWire connectome dataset is publicly available (https://codex.flywire.ai/?dataset=fafb). The MANC connectome dataset is publicly available (https://neuprint.janelia.org/). Any additional information required to reanalyze the data reported in this work is available from the lead contact upon request.

## Materials and Methods

### Immunostaining and confocal imaging of brains and ventral nerve cords

Brains of 8-10-day-old, fully fed, mated females and males expressing either UAS-GFP or UAS-RedStinger, UAS-mCD8:GFP under the control of specific GAL4 driver lines were used. The lines used in this study are shown in Supplemental Table 1. Briefly, brains were dissected in 4°C 1X PBS (10173433, Fisher Scientific, UK) after a quick passage through EtOH (4146052, Carlo Elba) and were then transferred to formaldehyde solution (4% paraformaldehyde, P6148, Sigma-Aldrich in 1X PBS + 10% Triton-X, X100, Sigma-Aldrich) and incubated for 20-30 min at room temperature. Samples were then washed three times in PBST (0.5% Triton-X in PBS) and then blocked with 10% normal goat serum (16210-064, Invitrogen) in PBST for 15-60 min at room temperature. Samples were then incubated in primary antibody solutions (Rabbit anti-GFP, TP401, Torrey Pines Biolabs at 1:6000 and Mouse anti-NC82, Developmental Studies Hybridoma Bank at 1:10 in 5% normal goat serum in PBST). Primary antibody incubations were performed for 3 days at 4°C with rocking. They were then washed in PBST 2-3 times for 10-15 min at room temperature. The secondary antibodies were applied (Anti-mouse A594, A11032, Invitrogen at 1:500 and Anti-rabbit A488, A11008, Invitrogen at 1:500 in 5% normal goat serum in PBST) and brains were then incubated for 3 days at 4°C with rocking. They were again washed in PBST 2-3 times for 10-15 min at room temperature. Samples were mounted in Vectashield Mounting Medium (H-1000, Vector Laboratories). Images were captured on an inverted Zeiss LSM 980 (Carl Zeiss Co., Oberkochen, Germany) using a Plan-ApoChromat 20x/0.8 air lens objective (Carl Zeiss Co., Oberkochen, Germany).

### Confocal imaging of body parts

Wings, legs, and heads of flies expressing 20X-UAS-mCD8:GFP and UAS-RedStinger (#BDSC: 8546) under *poxn-GAL4* ^40^, *IR25a-GAL4* (#BDSC: 41728), or *R57F03-GAL4* (#BDSC: 46386) were dissected using fine forceps (Dumond #5 forcepts, F.S.T., US) and mounted on a coverslip in either 1X PBS or 10S mineral oil (24627.188, VWR, France). Images were captured on an inverted Zeiss LSM 980 (Carl Zeiss Co., Oberkochen, Germany) using a Plan-ApoChromat 20x/ 0.8 N.A. air lens objective (Carl Zeiss Co., Oberkochen, Germany) or a Plan-ApoChromat 25X/0.8 N.A. multiimmersion lens.

### GRN and MN identification, annotation, and proofreading

As sensory neurons, the GRNs’ axons of each subclass (labellar bristle, taste peg, pharyngeal, wing bristle, and leg bristle GRNs) project to the CNS through specific nerves. Thus, in the male CNS, the GRNs of each subclass were identified and annotated by defining seed planes across their respective entering nerves. In FAFB – FlyWire and MANC, the GRNs were identified similarly or using previously defined annotations. To identify and annotate the MNs in the male CNS, we searched for morphological matches to previously annotated MNs in FAFB – Flywire, in combination with defining seed planes through their exiting nerves. The automated segmentation of each GRN and MN was individually evaluated to determine if proofreading was required. Any relevant splits or merges were identified and performed.

### Connectomic partner analysis

The natverse package in R (R Development Core Team, version 4.1.0) was used to analyze the connectome dataset. The Dplyr package (https://github.com/tidyverse/dplyr) was used to facilitate data manipulation. To retrieve the synaptic-level connectome dataset and plot meshes, the fafbseg (for FAFB, FlyWire v783), malecns, and manc packages were used via the coconatfly package (https://github.com/natverse/coconatfly?tab=readme-ov-file) ^31^. For synaptic connectivity predictions, we used a cleft threshold = 50. Synaptic weight thresholds were indicated whenever applied. Graph representations of neuronal connectivity were plotted using Gephi (https://gephi.org/, version 0.10) to highlight the relative connectivity of nodes. The neurons were treated as nodes, and the synaptic connectivity between neurons was treated as directed edges scaled by the synaptic weights for each neuron pair. For the layout of the graphs, the Built-in YiFan Hu Proportional method in Gephi was used. YiFan Hu’s method is a time-efficient method based on force-directed algorithm models that aim to minimize the energy in the system iteratively.

### Matching neurons to the FAFB – FlyWire dataset

To cell type the GRNs and MNs in the male CNS based on their connectivity, we first cell typed their synaptic partners by matching them to their corresponding counterparts in FAFB – FlyWire ^14,31,61^. We started by iteratively grouping the partners of our neurons of interest with similar connectivity and morphology. To then assign a FAFB – FlyWire type to each group, we compared the morphologies of the male CNS neurons to their top five best matches in FAFB – FlyWire using neuroglancer, which allows the simultaneous visualization of male CNS meshes and transformed FlyWire meshes to the male CNS space. The top FAFB – FlyWire matches were found using pre-computed NBLAST scores resulting from comparisons between pairs of FAFB – FlyWire and male CNS neuron skeletons that were registered to a common template space ^60^. Finally, we selected the FAFB – FlyWire type with the closest morphology.

### GRN and MN cell typing

To cell type the GRNs and MNs across connectomes, we used connectivity clustering based, respectively, on their outputs and inputs. As sensory neurons, each GRN establishes very few connections, and, so, except for pharyngeal GRNs, we used a threshold of 2 synapses for each GRN subclass. For MNs, we used the standard threshold of 5 synapses. To cluster neurons from either dataset or, when co-clustering from two different connectomes, we used cosine similarity scores that were computed using the coconatfly package (https://github.com/natverse/coconatfly). Briefly, this method computes the cosine similarity for pairs of connectivity vectors for neurons in the n-dimensional synaptic connectivity space. To assign types to defined connectivity clusters, we visualized and compared the morphologies of the neurons in each cluster to confirm a match. A complementary, detailed inspection of top downstream partners and anatomical differences was used to further subtype the GRNs. The GRN and MN body IDs and their corresponding cell types are shown in Supplemental Table 2.

### Effective connectivity algorithm

To make predictions on how a neuron’s influence may impact another neuron across multiple synapses, we employed an effective connectivity algorithm based on serial matrix multiplication, adapted from a recent study ^94^. To calculate effective connectivity scores between a given input and output neuron (or population) in the male CNS or FAFB – FlyWire datasets, we used natverse, fafbseg, and malecns R tools to first construct an all-by-all connectivity matrix representing the absolute number of synaptic connections between every pair of neurons in the dataset. This was then converted into a proportional matrix, where each neuron-to-neuron connection is expressed as the fraction of the postsynaptic neuron’s total input synapses. Starting from a chosen set of ‘input’ neurons, we modeled the propagation of their influence through successive downstream layers by multiplying the corresponding rows of the proportional connectivity matrix by the full all-by-all proportional matrix. Each multiplication yielded an updated connectivity profile that represents how the input neuron’s influence is redistributed through the outputs of its direct partners, effectively extending the predicted impact one synaptic ‘hop’ further downstream. Iterating this procedure produced connectivity estimates across multiple layers of the network, up to a set number of ‘desired layers’. For each layer, we then extracted the effective connectivity values associated with a defined set of output neurons. To allow comparisons across inputs and layers, raw connection strengths were normalized by dividing each neuron’s score by the mean score of all neurons with non-zero values in that layer. This normalization ensured that effective connectivity scores reflected relative rather than absolute strengths, thereby highlighting disproportionately strong or weak pathways while controlling for global differences in connection density across layers.

### Calculating significant paths

To construct the significant contributing neurons in a circuit between focal input and output neurons (or populations), we serially calculated effective connectivity scores from a focal input type to trace the strongest routes of influence across multiple synaptic layers. Starting from the downstream partners of the chosen input neurons, we first computed their effective connectivity score to the target population as described above. For each downstream layer, we then identified neurons whose normalized effective connectivity scores exceeded a threshold (80th quantile), thereby selecting candidates that most strongly underpin the transmission of influence from input to output. To further refine these candidates, we applied a pruning step in which each neuron was also evaluated for its effective connectivity from the original input population, ensuring that only those nodes maintaining a strong connection from the input population were retained. This set of candidates was then selected as a new ‘input’ with which to calculate effective connectivity to the output node in the next iteration, until the desired number of layers was reached. From this procedure, we were left with a list of influential neurons within the input to output circuit, from which we generated an all-by-all connectivity matrix, which was in turn used to generate circuit graphs using the Gephi graphing software.

### Reachability analysis

To calculate reachability analysis between GRN and MN types, we ran successive iterations of effective connectivity calculations between each input and output pairing, while sequentially increasing the number of ‘desired layers’, used by the algorithm to model propagation of signal across multiple synaptic ‘hops’. The evolution of the effective connectivity scores between each cell type connection was then plotted as a function of the increasing synaptic layer value.

## Bibliography

1. Kristin Scott. Gustatory processing in drosophila melanogaster. Annual Review of Entomology, 63:15–30, 2018. ISSN 0066-4170, 1545-4487. doi: 10.1146/annurev-ento-020117-043331. Publisher: Annual Reviews.

2. Seok Jun Moon, Youngseok Lee, Yuchen Jiao, and Craig Montell. A Drosophila gustatory receptor essential for aversive taste and inhibiting male-to-male courtship. Current Biology, 19(19):1623–1627, 2009. ISSN 0960-9822. doi: 10.1016/j.cub.2009.07.061.

3. Wolf Huetteroth, Emmanuel Perisse, Suewei Lin, Martín Klappenbach, Christopher Burke, and Scott Waddell. Sweet taste and nutrient value subdivide rewarding dopaminergic neurons in Drosophila. Current Biology, 25(6):751–758, 2015. ISSN 0960-9822. doi: 10.1016/j.cub.2015.01.036.

4. Hsueh-Ling Chen, Dorsa Motevalli, Ulrich Stern, and Chung-Hui Yang. A functional division of drosophila sweet taste neurons that is value-based and task-specific. Proceedings of the National Academy of Sciences, 119(3):e2110158119, 2022. doi: 10.1073/pnas.2110158119. Publisher: Proceedings of the National Academy of Sciences.

5. Daniel Münch, Gili Ezra-Nevo, Ana Patrícia Francisco, Ibrahim Tastekin, and Carlos Ribeiro. Nutrient homeostasis - translating internal states to behavior. Current Opinion in Neurobiology, 60:67–75, 2020. ISSN 1873-6882. doi: 10.1016/j.conb.2019.10.004.

6. Hubert Amrein and Natasha Thorne. Gustatory perception and behavior in Drosophila melanogaster. Current Biology, 15(17):R673–R684, 2005. ISSN 0960-9822. doi: 10.1016/j.cub.2005.08.021.

7. Jae Young Kwon, Anupama Dahanukar, Linnea A. Weiss, and John R. Carlson. A map of taste neuron projections in the drosophila CNS. Journal of Biosciences, 39(4):565–574, 2014. ISSN 0973-7138. doi: 10.1007/s12038-014-9448-6.

8. Tong-Wey Koh, Zhe He, Srinivas Gorur-Shandilya, Karen Menuz, Nikki K. Larter, Shannon Stewart, and John R. Carlson. The drosophila IR20a clade of ionotropic receptors are candidate taste and pheromone receptors. Neuron, 83(4):850–865, 2014. ISSN 0896-6273. doi: 10.1016/j.neuron.2014.07.012. Publisher: Elsevier.

9. Juan Antonio Sánchez-Alcañiz, Ana Florencia Silbering, Vincent Croset, Giovanna Zappia, Anantha Krishna Sivasubramaniam, Liliane Abuin, Saumya Yashmohini Sahai, Daniel Münch, Kathrin Steck, Thomas O. Auer, Steeve Cruchet, G. Larisa Neagu-Maier, Simon G. Sprecher, Carlos Ribeiro, Nilay Yapici, and Richard Benton. An expression atlas of variant ionotropic glutamate receptors identifies a molecular basis of carbonation sensing. Nature Communications, 9(1):4252, 2018. ISSN 2041-1723. doi: 10.1038/s41467-018-06453-1. Publisher: Nature Publishing Group.

10. S. Shanbhag, S.-K. Park, C. Pikielny, and R. Steinbrecht. Gustatory organs of drosophila melanogaster: fine structure and expression of the putative odorant-binding protein PBPRP2. Cell and Tissue Research, 304(3):423–437, 2001. ISSN 1432-0878. doi: 10.1007/s004410100388.

11. D. R. Possidente and R. K. Murphey. Genetic control of sexually dimorphic axon morphology in Drosophila sensory neurons. Developmental Biology, 132(2):448–457, 1989. ISSN 0012-1606. doi: 10.1016/0012-1606(89)90241-8.

12. Jacqueline Guillemin, Jinfang Li, Viktoriya Li, Sasha A. T. McDowell, Kayla Audette, Grace Davis, Meghan Jelen, Samy Slamani, Liam Kelliher, Michael D. Gordon, and Molly Stanley. Taste cells expressing ionotropic receptor 94e reciprocally impact feeding and egg laying in drosophila. Cell Reports, 43(8), 2024. ISSN 2211-1247. doi: 10.1016/j.celrep.2024.114625. Publisher: Elsevier.

13. Louis K Scheffer, C Shan Xu, Michal Januszewski, Zhiyuan Lu, Shin-ya Takemura, Kenneth J Hayworth, Gary B Huang, Kazunori Shinomiya, Jeremy Maitlin-Shepard, Stuart Berg, Jody Clements, Philip M Hubbard, William T Katz, Lowell Umayam, Ting Zhao, David Ackerman, Tim Blakely, John Bogovic, Tom Dolafi, Dagmar Kainmueller, Takashi Kawase, Khaled A Khairy, Laramie Leavitt, Peter H Li, Larry Lindsey, Nicole Neubarth, Donald J Olbris, Hideo Otsuna, Eric T Trautman, Masayoshi Ito, Alexander S Bates, Jens Goldammer, Tanya Wolff, Robert Svirskas, Philipp Schlegel, Erika Neace, Christopher J Knecht, Chelsea X Alvarado, Dennis A Bailey, Samantha Ballinger, Jolanta A Borycz, Brandon S Canino, Natasha Cheatham, Michael Cook, Marisa Dreher, Octave Duclos, Bryon Eubanks, Kelli Fairbanks, Samantha Finley, Nora Forknall, Audrey Francis, Gary Patrick Hopkins, Emily M Joyce, SungJin Kim, Nicole A Kirk, Julie Kovalyak, Shirley A Lauchie, Alanna Lohff, Charli Maldonado, Emily A Manley, Sari McLin, Caroline Mooney, Miatta Ndama, Omotara Ogundeyi, Nneoma Okeoma, Christopher Ordish, Nicholas Padilla, Christopher M Patrick, Tyler Paterson, Elliott E Phillips, Emily M Phillips, Neha Rampally, Caitlin Ribeiro, Madelaine K Robertson, Jon Thomson Rymer, Sean M Ryan, Megan Sammons, Anne K Scott, Ashley L Scott, Aya Shinomiya, Claire Smith, Kelsey Smith, Natalie L Smith, Margaret A Sobeski, Alia Suleiman, Jackie Swift, Satoko Takemura, Iris Talebi, Dorota Tarnogorska, Emily Tenshaw, Temour Tokhi, John J Walsh, Tansy Yang, Jane Anne Horne, Feng Li, Ruchi Parekh, Patricia K Rivlin, Vivek Jayaraman, Marta Costa, Gregory SXE Jefferis, Kei Ito, Stephan Saalfeld, Reed George, Ian A Meinertzhagen, Gerald M Rubin, Harald F Hess, Viren Jain, and Stephen M Plaza. A connectome and analysis of the adult drosophila central brain. eLife, 9:e57443, 2020. ISSN 2050-084X. doi: 10.7554/eLife.57443. Publisher: eLife Sciences Publications, Ltd.

14. Sven Dorkenwald, Arie Matsliah, Amy R. Sterling, Philipp Schlegel, Szi-chieh Yu, Claire E. McKellar, Albert Lin, Marta Costa, Katharina Eichler, Yijie Yin, Will Silversmith, Casey Schneider-Mizell, Chris S. Jordan, Derrick Brittain, Akhilesh Halageri, Kai Kuehner, Oluwaseun Ogedengbe, Ryan Morey, Jay Gager, Krzysztof Kruk, Eric Perlman, Runzhe Yang, David Deutsch, Doug Bland, Marissa Sorek, Ran Lu, Thomas Macrina, Kisuk Lee, J. Alexander Bae, Shang Mu, Barak Nehoran, Eric Mitchell, Sergiy Popovych, Jingpeng Wu, Zhen Jia, Manuel A. Castro, Nico Kemnitz, Dodam Ih, Alexander Shakeel Bates, Nils Eckstein, Jan Funke, Forrest Collman, Davi D. Bock, Gregory S. X. E. Jefferis, H. Sebastian Seung, and Mala Murthy. Neuronal wiring diagram of an adult brain. Nature, 634 (8032):124–138, 2024. ISSN 1476-4687. doi: 10.1038/s41586-024-07558-y. Publisher: Nature Publishing Group.

15. Lisa S. Baik and John R. Carlson. The mosquito taste system and disease control. Proceedings of the National Academy of Sciences, 117(52):32848–32856, 2020. doi: 10.1073/pnas.2013076117. Publisher: Proceedings of the National Academy of Sciences.

16. Lisa S. Baik, Gaëlle J. S. Talross, Sydney Gray, Himani S. Pattisam, Taylor N. Peterson, James E. Nidetz, Felix J. H. Hol, and John R. Carlson. Mosquito taste responses to human and floral cues guide biting and feeding. Nature, 635(8039):639–646, 2024. ISSN 1476-4687. doi: 10.1038/s41586-024-08047-y. Publisher: Nature Publishing Group.

17. Felicity Muth, Jacob S. Francis, and Anne S. Leonard. Bees use the taste of pollen to determine which flowers to visit. Biology Letters, 12(7):20160356, 2016. ISSN 1744-9561. doi: 10.1098/rsbl.2016.0356.

18. Hany KM Dweck, Gaëlle JS Talross, Wanyue Wang, and John R Carlson. Evolutionary shifts in taste coding in the fruit pest drosophila suzukii. eLife, 10:e64317, 2021. ISSN 2050-084X. doi: 10.7554/eLife.64317. Publisher: eLife Sciences Publications, Ltd.

19. V. G. Dethier. The hungry fly: A physiological study of the behavior associated with feeding. The hungry fly: A physiological study of the behavior associated with feeding. Harvard U Press, 1976. Pages: 489.

20. R. F. Stocker. The organization of the chemosensory system in drosophila melanogaster: a review. Cell and Tissue Research, 275(1):3–26, 1994. ISSN 0302-766X. doi: 10.1007/BF00305372.

21. K. P. Rajashekhar and R. N. Singh. Organization of motor neurons innervating the proboscis musculature in Drosophila melanogaster meigen (diptera : Drosophilidae). International Journal of Insect Morphology and Embryology, 23(3):225–242, 1994. ISSN 0020-7322. doi: 10.1016/0020-7322(94)90020-5.

22. Michael D. Gordon and Kristin Scott. Motor control in a Drosophila taste circuit. Neuron, 61(3):373–384, 2009. ISSN 0896-6273. doi: 10.1016/j.neuron.2008.12.033.

23. Thomas F. Flood, Shinya Iguchi, Michael Gorczyca, Benjamin White, Kei Ito, and Motojiro Yoshihara. A single pair of interneurons commands the drosophila feeding motor program. Nature, 499(7456):83–87, 2013. ISSN 1476-4687. doi: 10.1038/nature12208. Publisher: Nature Publishing Group.

24. Olivia Schwarz, Ali Asgar Bohra, Xinyu Liu, Heinrich Reichert, Krishnaswamy VijayRaghavan, and Jan Pielage. Motor control of drosophila feeding behavior. eLife, 6:e19892, 2017. ISSN 2050-084X. doi: 10.7554/eLife.19892. Publisher: eLife Sciences Publications, Ltd.

25. Claire E McKellar, Igor Siwanowicz, Barry J Dickson, and Julie H Simpson. Controlling motor neurons of every muscle for fly proboscis reaching. eLife, 9:e54978, 2020. ISSN 2050-084X. doi: 10.7554/eLife.54978. Publisher: eLife Sciences Publications, Ltd.

26. Neha Sapkal, Nino Mancini, Divya Sthanu Kumar, Nico Spiller, Kazuma Murakami, Gianna Vitelli, Benjamin Bargeron, Kate Maier, Katharina Eichler, Gregory S. X. E. Jefferis, Philip K. Shiu, Gabriella R. Sterne, and Salil S. Bidaye. Neural circuit mechanisms underlying context-specific halting in drosophila. Nature, 634(8032):191–200, 2024. ISSN 1476-4687. doi: 10.1038/s41586-024-07854-7. Publisher: Nature Publishing Group.

27. Aljoscha Nern, Frank Loesche, Shin-ya Takemura, Laura E. Burnett, Marisa Dreher, Eyal Gruntman, Judith Hoeller, Gary B. Huang, Michał Januszewski, Nathan C. Klapoetke, Sanna Koskela, Kit D. Longden, Zhiyuan Lu, Stephan Preibisch, Wei Qiu, Edward M. Rogers, Pavithraa Seenivasan, Arthur Zhao, John Bogovic, Brandon S. Canino, Jody Clements, Michael Cook, Samantha Finley-May, Miriam A. Flynn, Imran Hameed, Alexandra M. C. Fragniere, Kenneth J. Hayworth, Gary Patrick Hopkins, Philip M. Hubbard, William T. Katz, Julie Kovalyak, Shirley A. Lauchie, Meghan Leonard, Alanna Lohff, Charli A. Maldonado, Caroline Mooney, Nneoma Okeoma, Donald J. Olbris, Christopher Ordish, Tyler Paterson, Emily M. Phillips, Tobias Pietzsch, Jennifer Rivas Salinas, Patricia K. Rivlin, Philipp Schlegel, Ashley L. Scott, Louis A. Scuderi, Satoko Takemura, Iris Talebi, Alexander Thomson, Eric T. Trautman, Lowell Umayam, Claire Walsh, John J. Walsh, C. Shan Xu, Emily A. Yakal, Tansy Yang, Ting Zhao, Jan Funke, Reed George, Harald F. Hess, Gregory S. X. E. Jefferis, Christopher Knecht, Wyatt Korff, Stephen M. Plaza, Sandro Romani, Stephan Saalfeld, Louis K. Scheffer, Stuart Berg, Gerald M. Rubin, and Michael B. Reiser. Connectome-driven neural inventory of a complete visual system. Nature, 641(8065):1225–1237, 2025. ISSN 1476-4687. doi: 10.1038/s41586-025-08746-0. Publisher: Nature Publishing Group.

28. Alexander S. Bates, Philipp Schlegel, Ruairi J. V. Roberts, Nikolas Drummond, Imaan F. M. Tamimi, Robert Turnbull, Xincheng Zhao, Elizabeth C. Marin, Patricia D. Popovici, Serene Dhawan, Arian Jamasb, Alexandre Javier, Laia Serratosa Capdevila, Feng Li, Gerald M. Rubin, Scott Waddell, Davi D. Bock, Marta Costa, and Gregory S. X. E. Jefferis. Complete connectomic reconstruction of olfactory projection neurons in the fly brain. Current biology: CB, 30(16):3183–3199.e6, 2020. ISSN 1879-0445. doi: 10.1016/j.cub.2020.06.042.

29. Katharina Eichler, Stefanie Hampel, Adrián Alejandro-García, Steven A. Calle-Schuler, Alexis Santana-Cruz, Lucia Kmecova, Jonathan M. Blagburn, Eric D. Hoopfer, and Andrew M. Seeds. Somatotopic organization among parallel sensory pathways that promote a grooming sequence in drosophila. eLife, 12, 2024. doi: 10.7554/eLife.87602.2. Publisher: eLife Sciences Publications Limited.

30. Zhihao Zheng, J. Scott Lauritzen, Eric Perlman, Camenzind G. Robinson, Matthew Nichols, Daniel Milkie, Omar Torrens, John Price, Corey B. Fisher, Nadiya Sharifi, Steven A. Calle-Schuler, Lucia Kmecova, Iqbal J. Ali, Bill Karsh, Eric T. Trautman, John A. Bogovic, Philipp Hanslovsky, Gregory S. X. E. Jefferis, Michael Kazhdan, Khaled Khairy, Stephan Saalfeld, Richard D. Fetter, and Davi D. Bock. A complete electron microscopy volume of the brain of adult Drosophila melanogaster. Cell, 174(3):730–743.e22, 2018. ISSN 0092-8674. doi: 10.1016/j.cell.2018.06.019.

31. Philipp Schlegel, Yijie Yin, Alexander S. Bates, Sven Dorkenwald, Katharina Eichler, Paul Brooks, Daniel S. Han, Marina Gkantia, Marcia dos Santos, Eva J. Munnelly, Griffin Badalamente, Laia Serratosa Capdevila, Varun A. Sane, Alexandra M. C. Fragniere, Ladann Kiassat, Markus W. Pleijzier, Tomke Stürner, Imaan F. M. Tamimi, Christopher R. Dunne, Irene Salgarella, Alexandre Javier, Siqi Fang, Eric Perlman, Tom Kazimiers, Sridhar R. Jagannathan, Arie Matsliah, Amy R. Sterling, Szi-chieh Yu, Claire E. McKellar, Marta Costa, H. Sebastian Seung, Mala Murthy, Volker Hartenstein, Davi D. Bock, and Gregory S. X. E. Jefferis. Whole-brain annotation and multi-connectome cell typing of drosophila. Nature, 634(8032):139–152, 2024. ISSN 1476-4687. doi: 10.1038/s41586-024-07686-5. Publisher: Nature Publishing Group.

32. Shin-ya Takemura, Kenneth J. Hayworth, Gary B. Huang, Michal Januszewski, Zhiyuan Lu, Elizabeth C. Marin, Stephan Preibisch, C. Shan Xu, John Bogovic, Andrew S. Champion, Han SJ Cheong, Marta Costa, Katharina Eichler, William Katz, Christopher Knecht, Feng Li, Billy J. Morris, Christopher Ordish, Patricia K. Rivlin, Philipp Schlegel, Kazunori Shinomiya, Tomke Stürner, Ting Zhao, Griffin Badalamente, Dennis Bailey, Paul Brooks, Brandon S. Canino, Jody Clements, Michael Cook, Octave Duclos, Christopher R. Dunne, Kelli Fairbanks, Siqi Fang, Samantha Finley-May, Audrey Francis, Reed George, Marina Gkantia, Kyle Harrington, Gary Patrick Hopkins, Joseph Hsu, Philip M. Hubbard, Alexandre Javier, Dagmar Kainmueller, Wyatt Korff, Julie Kovalyak, Dominik Krzeminski, Shirley A. Lauchie, Alanna Lohff, Charli Maldonado, Emily A. Manley, Caroline Mooney, Erika Neace, Matthew Nichols, Omotara Ogundeyi, Nneoma Okeoma, Tyler Paterson, Elliott Phillips, Emily M. Phillips, Caitlin Ribeiro, Sean M. Ryan, Jon Thomson Rymer, Anne K. Scott, Ashley L. Scott, David Shepherd, Aya Shinomiya, Claire Smith, Natalie Smith, Alia Suleiman, Satoko Takemura, Iris Talebi, Imaan FM Tamimi, Eric T. Trautman, Lowell Umayam, John J. Walsh, Tansy Yang, Gerald M. Rubin, Louis K. Scheffer, Jan Funke, Stephan Saalfeld, Harald F. Hess, Stephen M. Plaza, Gwyneth M. Card, Gregory SXE Jefferis, and Stuart Berg. A connectome of the male drosophila ventral nerve cord. eLife, 13, 2024. doi: 10.7554/eLife.97769.1. Publisher: eLife Sciences Publications Limited.

33. Elizabeth C. Marin, Billy J. Morris, Tomke Stürner, Andrew S. Champion, Dominik Krzeminski, Griffin Badalamente, Marina Gkantia, Christopher R. Dunne, Katharina Eichler, Shin-ya Takemura, Imaan FM Tamimi, Siqi Fang, Sung Soo Moon, Han SJ Cheong, Feng Li, Philipp Schlegel, Sebastian E. Ahnert, Stuart Berg, Janelia FlyEM Project Team, Gwyneth M. Card, Marta Costa, David Shepherd, and Gregory SXE Jefferis. Systematic annotation of a complete adult male drosophila nerve cord connectome reveals principles of functional organisation. eLife, 13, 2024. doi: 10.7554/eLife.97766.1. Publisher: eLife Sciences Publications Limited.

34. Han SJ Cheong, Katharina Eichler, Tomke Stürner, Samuel K. Asinof, Andrew S. Champion, Elizabeth C. Marin, Tess B. Oram, Marissa Sumathipala, Lalanti Venkatasubramanian, Shigehiro Namiki, Igor Siwanowicz, Marta Costa, Stuart Berg, Janelia FlyEM Project Team, Gregory SXE Jefferis, and Gwyneth M. Card. Transforming descending input into motor output: An analysis of the drosophila male adult nerve cord connectome. eLife, 13, 2025. doi: 10.7554/eLife.96084.2. Publisher: eLife Sciences Publications Limited.

35. Stefanie Engert, Gabriella R Sterne, Davi D Bock, and Kristin Scott. Drosophila gustatory projections are segregated by taste modality and connectivity. eLife, 11:e78110, 2022. ISSN 2050-084X. doi: 10.7554/eLife.78110. Publisher: eLife Sciences Publications, Ltd.

36. Shubha V. Nayak and R. Naresh Singh. Sensilla on the tarsal segments and mouthparts of adult Drosophila melanogaster meigen (diptera : Drosophilidae). International Journal of Insect Morphology and Embryology, 12(5):273–291, 1983. ISSN 0020-7322. doi: 10.1016/0020-7322(83)90023-5.

37. John Palka, Peter A. Lawrence, and H. Stephen Hart. Neural projection patterns from homeotic tissue of Drosophila studied in bithorax mutants and mosaics. Developmental Biology, 69(2):549–575, 1979. ISSN 0012-1606. doi: 10.1016/0012-1606(79)90311-7.

38. John Palka. Neuronal specificity and its development in the drosophila wing disc and its derivatives. Journal of Neurobiology, 24(6):788–802, 1993. ISSN 1097-4695. doi: 10.1002/neu.480240607. _eprint: https://onlinelibrary.wiley.com/doi/pdf/10.1002/neu.480240607.

39. Yu-Chieh David Chen and Anupama Dahanukar. Molecular and cellular organization of taste neurons in adult Drosophila pharynx. Cell Reports, 21(10):2978–2991, 2017. ISSN 2211-1247. doi: 10.1016/j.celrep.2017.11.041.

40. Werner Boll and Markus Noll. The drosophila pox neuro gene: control of male courtship behavior and fertility as revealed by a complete dissection of all enhancers. Development (Cambridge, England), 129(24):5667–5681, 2002. ISSN 0950-1991. doi: 10.1242/dev.00157.

41. Vladimiros Thoma, Stephan Knapek, Shogo Arai, Marion Hartl, Hiroshi Kohsaka, Pudith Sirigrivatanawong, Ayako Abe, Koichi Hashimoto, and Hiromu Tanimoto. Functional dissociation in sweet taste receptor neurons between and within taste organs of drosophila. Nature Communications, 7(1):10678, 2016. ISSN 2041-1723. doi: 10.1038/ncomms10678. Publisher: Nature Publishing Group.

42. H. M. Robertson. Chemical stimuli eliciting courtship by males inDrosophila melanogaster. Experientia, 39(3):333–335, 1983. ISSN 0014-4754. doi: 10.1007/BF01955335.

43. Beika Lu, Angela LaMora, Yishan Sun, Michael J. Welsh, and Yehuda Ben-Shahar. ppk23-dependent chemosensory functions contribute to courtship behavior in drosophila melanogaster. PLOS Genetics, 8(3):e1002587, 2012. ISSN 1553-7404. doi: 10.1371/journal.pgen.1002587. Publisher: Public Library of Science.

44. Robert Thistle, Peter Cameron, Azeen Ghorayshi, Lisa Dennison, and Kristin Scott. Contact chemoreceptors mediate male-male repulsion and male-female attraction during drosophila courtship. Cell, 149(5):1140–1151, 2012. ISSN 0092-8674, 1097-4172. doi: 10.1016/j.cell.2012.03.045. Publisher: Elsevier.

45. Hussein Raad, Jean-François Ferveur, Neil Ledger, Maria Capovilla, and Alain Robichon. Functional gustatory role of chemoreceptors in Drosophila wings. Cell Reports, 15(7): 1442–1454, 2016. ISSN 2211-1247. doi: 10.1016/j.celrep.2016.04.040.

46. Zhe He, Yichen Luo, Xueying Shang, Jennifer S. Sun, and John R. Carlson. Chemosensory sensilla of the drosophila wing express a candidate ionotropic pheromone receptor. PLOS Biology, 17(5):e2006619, 2019. ISSN 1545-7885. doi: 10.1371/journal.pbio.2006619. Publisher: Public Library of Science.

47. Philip K Shiu, Gabriella R Sterne, Stefanie Engert, Barry J Dickson, and Kristin Scott. Taste quality and hunger interactions in a feeding sensorimotor circuit. eLife, 11:e79887, 2022. ISSN 2050-084X. doi: 10.7554/eLife.79887. Publisher: eLife Sciences Publications, Ltd.

48. Kathrin Steck, Samuel J Walker, Pavel M Itskov, Célia Baltazar, José-Maria Moreira, and Carlos Ribeiro. Internal amino acid state modulates yeast taste neurons to support protein homeostasis in drosophila. eLife, 7:e31625, 2018. ISSN 2050-084X. doi: 10.7554/eLife.31625. Publisher: eLife Sciences Publications, Ltd.

49. I. Tastekin, I. de Haan Vicente, R. J. Beresford, N. Otto, G. Dempsey, S. Waddell, and C. Ribeiro. Connectomics reveals a feed-forward swallowing circuit driving protein appetite. bioRxiv, 2025. doi: 10.1101/2025.08.25.671815. ISSN: 2692-8205 Pages: 2025.08.25.671815 Section: New Results.

50. Yan Chen and Hubert Amrein. Ionotropic receptors mediate drosophila oviposition preference through sour gustatory receptor neurons. Current biology, 27(18):2741–2750.e4, 2017. ISSN 1879-0445. doi: 10.1016/j.cub.2017.08.003.

51. Ji-Eun Ahn, Yan Chen, and Hubert Amrein. Molecular basis of fatty acid taste in drosophila. eLife, 6:e30115, 2017. ISSN 2050-084X. doi: 10.7554/eLife.30115. Publisher: eLife Sciences Publications, Ltd.

52. R. Naresh Singh and Shubha V. Nayak. Fine structure and primary sensory projections of sensilla on the maxillary palp of Drosophila melanogaster meigen (diptera : Drosophilidae). International Journal of Insect Morphology and Embryology, 14(5):291–306, 1985. ISSN 0020-7322. doi: 10.1016/0020-7322(85)90044-3.

53. Emily E. LeDue, Yu-Chieh Chen, Aera Y. Jung, Anupama Dahanukar, and Michael D. Gordon. Pharyngeal sense organs drive robust sugar consumption in drosophila. Nature Communications, 6(1):6667, 2015. ISSN 2041-1723. doi: 10.1038/ncomms7667. Publisher: Nature Publishing Group.

54. Jiun Sang, Subash Dhakal, Bhanu Shrestha, Dharmendra Kumar Nath, Yunjung Kim, Anindya Ganguly, Craig Montell, and Youngseok Lee. A single pair of pharyngeal neurons functions as a commander to reject high salt in drosophila melanogaster. eLife, 12, 2024. doi: 10.7554/eLife.93464.2. Publisher: eLife Sciences Publications Limited.

55. Yong Taek Jeong, Soo Min Oh, Jaewon Shim, Jeong Taeg Seo, Jae Young Kwon, and Seok Jun Moon. Mechanosensory neurons control sweet sensing in drosophila. Nature Communications, 7(1):12872, 2016. ISSN 2041-1723. doi: 10.1038/ncomms12872. Publisher: Nature Publishing Group.

56. Yao Zhou, Li-Hui Cao, Xiu-Wen Sui, Xiao-Qing Guo, and Dong-Gen Luo. Mechanosensory circuits coordinate two opposing motor actions in drosophila feeding. Science Advances, 5(5):eaaw5141, 2019. doi: 10.1126/sciadv.aaw5141. Publisher: American Association for the Advancement of Science.

57. Takaaki Miyazaki and Kei Ito. Neural architecture of the primary gustatory center of drosophila melanogaster visualized with GAL4 and LexA enhancer-trap systems. Journal of Comparative Neurology, 518(20):4147–4181, 2010. ISSN 1096-9861. doi: 10.1002/cne.22433. _eprint: https://onlinelibrary.wiley.com/doi/pdf/10.1002/cne.22433.

58. R. K. Murphey, Debra Possidente, Gerald Pollack, and D. J. Merritt. Modality-specific axonal projections in the CNS of the flies phormia and drosophila. Journal of Comparative Neurology, 290(2):185–200, 1989. ISSN 1096-9861. doi: 10.1002/cne.902900203. _eprint: https://onlinelibrary.wiley.com/doi/pdf/10.1002/cne.902900203.

59. R. N. Singh. Neurobiology of the gustatory systems of drosophila and some terrestrial insects. Microscopy Research and Technique, 39(6):547–563, 1997. ISSN 1059-910X. doi: 10.1002/(SICI)1097-0029(19971215)39:6<547::AID-JEMT7>3.0.CO;2-A.

60. Stuart Berg, Isabella R. Beckett, Marta Costa, Philipp Schlegel, Michał Januszewski, Elizabeth C. Marin, Aljoscha Nern, Stephan Preibisch, Wei Qiu, Shin-ya Takemura, Alexandra MC Fragniere, Andrew S. Champion, Diane-Yayra Adjavon, Michael Cook, Marina Gkantia, Kenneth J. Hayworth, Gary B. Huang, William T. Katz, Florian Kämpf, Zhiyuan Lu, Christopher Ordish, Tyler Paterson, Tomke Stürner, Eric T. Trautman, Catherine R. Whittle, Laura E. Burnett, Judith Hoeller, Feng Li, Frank Loesche, Billy J. Morris, Tobias Pietzsch, Markus W. Pleijzier, Valeria Silva, Yijie Yin, Iris Ali, Griffin Badalamente, Alexander Shakeel Bates, John Bogovic, Paul Brooks, Sebastian Cachero, Brandon S. Canino, Bhumpanya Chaisrisawatsuk, Jody Clements, Arthur Crowe, Inês de Haan Vicente, Georgia Dempsey, Erika Donà, Márcia dos Santos, Marisa Dreher, Christopher R. Dunne, Katharina Eichler, Samantha Finley-May, Miriam A. Flynn, Imran Hameed, Gary Patrick Hopkins, Philip M. Hubbard, Ladann Kiassat, Julie Kovalyak, Shirley A. Lauchie, Meghan Leonard, Alanna Lohff, Kit D. Longden, Charli A. Maldonado, Myrto Mitletton, Ilina Moitra, Sung Soo Moon, Caroline Mooney, Eva J. Munnelly, Nneoma Okeoma, Donald J. Olbris, Anika Pai, Birava Patel, Emily M. Phillips, Stephen M. Plaza, Alana Richards, Jennifer Rivas Salinas, Ruairí JV Roberts, Edward M. Rogers, Ashley L. Scott, Louis A. Scuderi, Pavithraa Seenivasan, Laia Serratosa Capdevila, Claire Smith, Rob Svirskas, Satoko Takemura, Ibrahim Tastekin, Alexander Thomson, Lowell Umayam, John J. Walsh, Holly Whittome, C. Shan Xu, Emily A. Yakal, Tansy Yang, Arthur Zhao, Reed George, Viren Jain, Vivek Jayaraman, Wyatt Korff, Geoffrey W. Meissner, Sandro Romani, Jan Funke, Christopher Knecht, Stephan Saalfeld, Louis K. Scheffer, Scott Waddell, Gwyneth M. Card, Carlos Ribeiro, Michael B. Reiser, Harald F. Hess, Gerald M. Rubin, and Gregory SXE Jefferis. Sexual dimorphism in the complete connectome of the drosophila male central nervous system. bioRxiv, 2025. doi: 10.1101/2025.10.09.680999. ISSN: 2692-8205 Pages: 2025.10.09.680999 Section: New Results.

61. Nils Eckstein, Alexander Shakeel Bates, Andrew Champion, Michelle Du, Yijie Yin, Philipp Schlegel, Alicia Kun-Yang Lu, Thomson Rymer, Samantha Finley-May, Tyler Paterson, Ruchi Parekh, Sven Dorkenwald, Arie Matsliah, Szi-Chieh Yu, Claire McKellar, Amy Sterling, Katharina Eichler, Marta Costa, Sebastian Seung, Mala Murthy, Volker Hartenstein, Gregory S. X. E. Jefferis, and Jan Funke. Neurotransmitter classification from electron microscopy images at synaptic sites in Drosophila melanogaster. Cell, 187(10):2574–2594.e23, 2024. ISSN 0092-8674. doi: 10.1016/j.cell.2024.03.016.

62. Linnea A. Weiss, Anupama Dahanukar, Jae Young Kwon, Diya Banerjee, and John R. Carlson. The molecular and cellular basis of bitter taste in Drosophila. Neuron, 69(2): 258–272, 2011. ISSN 0896-6273. doi: 10.1016/j.neuron.2011.01.001.

63. Shinsuke Fujii, Ahmet Yavuz, Jesse Slone, Christopher Jagge, Xiangyu Song, and Hubert Amrein. Drosophila sugar receptors in sweet taste perception, olfaction and internal nutrient sensing. Current biology : CB, 25(5):621–627, 2015. ISSN 0960-9822. doi: 10.1016/j.cub.2014.12.058.

64. Alexandria H Jaeger, Molly Stanley, Zachary F Weiss, Pierre-Yves Musso, Rachel CW Chan, Han Zhang, Damian Feldman-Kiss, and Michael D Gordon. A complex peripheral code for salt taste in drosophila. eLife, 7:e37167, 2018. ISSN 2050-084X. doi: 10.7554/eLife.37167. Publisher: eLife Sciences Publications, Ltd.

65. Sasha A. T. McDowell, Molly Stanley, and Michael D. Gordon. A molecular mechanism for high salt taste in Drosophila. Current Biology, 32(14):3070–3081.e5, 2022. ISSN 0960-9822. doi: 10.1016/j.cub.2022.06.012.

66. Sydney R. Walker, Marco Peña-Garcia, and Anita V. Devineni. Connectomic analysis of taste circuits in drosophila. Scientific Reports, 15(1):5278, 2025. ISSN 2045-2322. doi: 10.1038/s41598-025-89088-9. Publisher: Nature Publishing Group.

67. Jinfang Li, Rabiah Dhaliwal, Molly Stanley, Pierre Junca, and Michael D. Gordon. Functional imaging and connectome analyses reveal organizing principles of taste circuits in drosophila. Current Biology, 35(10):2391–2405.e4, 2025. ISSN 0960-9822. doi: 10.1016/j.cub.2025.04.035. Publisher: Elsevier.

68. Elena Starostina, Tong Liu, Vinoy Vijayan, Zheng Zheng, Kathleen K. Siwicki, and Claudio W. Pikielny. A drosophila DEG/ENaC subunit functions specifically in gustatory neurons required for male courtship behavior. Journal of Neuroscience, 32(13):4665–4674, 2012. ISSN 0270-6474, 1529-2401. doi: 10.1523/JNEUROSCI.6178-11.2012. Publisher: Society for Neuroscience Section: Articles.

69. Raphael Rytz, Vincent Croset, and Richard Benton. Ionotropic receptors (IRs): Chemosensory ionotropic glutamate receptors in Drosophila and beyond. Insect Biochemistry and Molecular Biology, 43(9):888–897, 2013. ISSN 0965-1748. doi: 10.1016/j.ibmb.2013.02.007.

70. Sang Hoon Kim, Youngseok Lee, Bradley Akitake, Owen M. Woodward, William B. Guggino, and Craig Montell. Drosophila TRPA1 channel mediates chemical avoidance in gustatory receptor neurons. Proceedings of the National Academy of Sciences, 107(18): 8440–8445, 2010. doi: 10.1073/pnas.1001425107. Publisher: Proceedings of the National Academy of Sciences.

71. Tsuyoshi Inoshita and Teiichi Tanimura. Cellular identification of water gustatory receptor neurons and their central projection pattern in drosophila. Proceedings of the National Academy of Sciences, 103(4):1094–1099, 2006. doi: 10.1073/pnas.0502376103. Publisher: Proceedings of the National Academy of Sciences.

72. Peter Cameron, Makoto Hiroi, John Ngai, and Kristin Scott. The molecular basis for water taste in drosophila. Nature, 465(7294):91–95, 2010. ISSN 1476-4687. doi: 10.1038/nature09011. Publisher: Nature Publishing Group.

73. Hany K. M. Dweck, Gaëlle J. S. Talross, Yichen Luo, Shimaa A. M. Ebrahim, and John R. Carlson. Ir56b is an atypical ionotropic receptor that underlies appetitive salt response in Drosophila. Current Biology, 32(8):1776–1787.e4, 2022. ISSN 0960-9822. doi: 10.1016/j.cub.2022.02.063.

74. Xiaonan Li, Yuanjie Sun, Shan Gao, Yan Li, Li Liu, and Yan Zhu. Taste coding of heavy metal ion-induced avoidance in Drosophila. iScience, 26(5):106607, 2023. ISSN 2589-0042. doi: 10.1016/j.isci.2023.106607.

75. Walter Fischler, Priscilla Kong, Sunanda Marella, and Kristin Scott. The detection of carbonation by the drosophila gustatory system. Nature, 448(7157):1054–1057, 2007. ISSN 1476-4687. doi: 10.1038/nature06101. Publisher: Nature Publishing Group.

76. Zev Wisotsky, Adriana Medina, Erica Freeman, and Anupama Dahanukar. Evolutionary differences in food preference rely on gr64e, a receptor for glycerol. Nature Neuroscience, 14(12):1534–1541, 2011. ISSN 1546-1726. doi: 10.1038/nn.2944. Publisher: Nature Publishing Group.

77. Elizabeth B Brown, Kreesha D Shah, Justin Palermo, Manali Dey, Anupama Dahanukar, and Alex C Keene. Ir56d-dependent fatty acid responses in drosophila uncover taste discrimination between different classes of fatty acids. eLife, 10:e67878, 2021. ISSN 2050-084X. doi: 10.7554/eLife.67878. Publisher: eLife Sciences Publications, Ltd.

78. Yichen Luo, Gaëlle J.S. Talross, and John R. Carlson. Function and evolution of ir52 receptors in mate detection in drosophila. Current Biology, 34(23):5395–5408.e6, 2024. ISSN 09609822. doi: 10.1016/j.cub.2024.10.001.

79. István Taisz, Erika Donà, Daniel Münch, Shanice N. Bailey, Billy J. Morris, Kimberly I. Meechan, Katie M. Stevens, Irene Varela-Martínez, Marina Gkantia, Philipp Schlegel, Carlos Ribeiro, Gregory S.X.E. Jefferis, and Dana S. Galili. Generating parallel representations of position and identity in the olfactory system. Cell, 186(12):2556–2573.e22, 2023. ISSN 00928674. doi: 10.1016/j.cell.2023.04.038.

80. Frederick Ling, Anupama Dahanukar, Linnea A. Weiss, Jae Young Kwon, and John R. Carlson. The molecular and cellular basis of taste coding in the legs of drosophila. Journal of Neuroscience, 34(21):7148–7164, 2014. ISSN 0270-6474, 1529-2401. doi: 10.1523/JNEUROSCI.0649-14.2014. Publisher: Society for Neuroscience Section: Articles.

81. Heesoo Kim, Colleen Kirkhart, and Kristin Scott. Long-range projection neurons in the taste circuit of drosophila. eLife, 6:e23386, 2017. ISSN 2050-084X. doi: 10.7554/eLife.23386. Publisher: eLife Sciences Publications, Ltd.

82. Pu Fan, Devanand S. Manoli, Osama M. Ahmed, Yi Chen, Neha Agarwal, Sara Kwong, Allen G. Cai, Jeffrey Neitz, Adam Renslo, Bruce S. Baker, and Nirao M. Shah. Genetic and neural mechanisms that inhibit Drosophila from mating with other species. Cell, 154 (1):89–102, 2013. ISSN 0092-8674. doi: 10.1016/j.cell.2013.06.008.

83. Tetsuya Miyamoto and Hubert Amrein. Suppression of male courtship by a drosophila pheromone receptor. Nature Neuroscience, 11(8):874–876, 2008. ISSN 1546-1726. doi: 10.1038/nn.2161.

84. Benjamin R Kallman, Heesoo Kim, and Kristin Scott. Excitation and inhibition onto central courtship neurons biases drosophila mate choice. eLife, 4:e11188, 2015. ISSN 2050-084X. doi: 10.7554/eLife.11188. Publisher: eLife Sciences Publications, Ltd.

85. Anupama Dahanukar, Ya-Ting Lei, Jae Young Kwon, and John R. Carlson. Two Gr genes underlie sugar reception in Drosophila. Neuron, 56(3):503–516, 2007. ISSN 0896-6273. doi: 10.1016/j.neuron.2007.10.024.

86. Soh Kohatsu, Noriko Tanabe, Daisuke Yamamoto, and Kunio Isono. Which sugar to take and how much to take? two distinct decisions mediated by separate sensory channels. Frontiers in Molecular Neuroscience, 15, 2022. ISSN 1662-5099. doi: 10.3389/fnmol.2022.895395. Publisher: Frontiers.

87. Yu-Chieh David Chen, Scarlet Jinhong Park, Ryan Matthew Joseph, William W. Ja, and Anupama Arun Dahanukar. Combinatorial pharyngeal taste coding for feeding avoidance in adult drosophila. Cell Reports, 29(4):961–973.e4, 2019. ISSN 2211-1247. doi: 10.1016/j.celrep.2019.09.036.

88. Salil S. Bidaye, Meghan Laturney, Amy K. Chang, Yuejiang Liu, Till Bockemühl, Ansgar Büschges, and Kristin Scott. Two brain pathways initiate distinct forward walking programs in Drosophila. Neuron, 108(3):469–485.e8, 2020. ISSN 0896-6273. doi: 10.1016/j.neuron.2020.07.032.

89. Anton Miroschnikow, Philipp Schlegel, and Michael J. Pankratz. Making feeding decisions in the Drosophila nervous system. Current Biology, 30(14):R831–R840, 2020. ISSN 0960-9822. doi: 10.1016/j.cub.2020.06.036.

90. Daniel Münch, Dennis Goldschmidt, and Carlos Ribeiro. The neuronal logic of how internal states control food choice. Nature, 607(7920):747–755, 2022. ISSN 1476-4687. doi: 10.1038/s41586-022-04909-5.

91. Ibrahim Tastekin, Julia Riedl, Verena Schilling-Kurz, Alex Gomez-Marin, James W. Truman, and Matthieu Louis. Role of the subesophageal zone in sensorimotor control of orientation in Drosophila larva. Current Biology, 25(11):1448–1460, 2015. ISSN 0960-9822. doi: 10.1016/j.cub.2015.04.016.

92. Xinyue Cui, Matthew R. Meiselman, Staci N. Thornton, and Nilay Yapici. A gut-brain-gut interoceptive circuit loop gates sugar ingestion in drosophila. bioRxiv, 2024. doi: 10.1101/2024.09.02.610892. Pages: 2024.09.02.610892 Section: New Results.

93. Andrea Manzo, Marion Silies, Daryl M. Gohl, and Kristin Scott. Motor neurons controlling fluid ingestion in drosophila. Proceedings of the National Academy of Sciences of the United States of America, 109(16):6307–6312, 2012. ISSN 0027-8424. doi: 10.1073/pnas.1120305109.

94. Tomke Stürner, Paul Brooks, Laia Serratosa Capdevila, Billy J. Morris, Alexandre Javier, Siqi Fang, Marina Gkantia, Sebastian Cachero, Isabella R. Beckett, Elizabeth C. Marin, Philipp Schlegel, Andrew S. Champion, Ilina Moitra, Alana Richards, Finja Klemm, Leonie Kugel, Shigehiro Namiki, Han S. J. Cheong, Julie Kovalyak, Emily Tenshaw, Ruchi Parekh, Jasper S. Phelps, Brandon Mark, Sven Dorkenwald, Alexander S. Bates, Arie Matsliah, Szi-chieh Yu, Claire E. McKellar, Amy Sterling, H. Sebastian Seung, Mala Murthy, John C. Tuthill, Wei-Chung Allen Lee, Gwyneth M. Card, Marta Costa, Gregory S. X. E. Jefferis, and Katharina Eichler. Comparative connectomics of drosophila descending and ascending neurons. Nature, 643(8070):158–172, 2025. ISSN 1476-4687. doi: 10.1038/s41586-025-08925-z. Publisher: Nature Publishing Group.

95. Pavel M. Itskov, José-Maria Moreira, Ekaterina Vinnik, Gonçalo Lopes, Steve Safarik, Michael H. Dickinson, and Carlos Ribeiro. Automated monitoring and quantitative analysis of feeding behaviour in drosophila. Nature Communications, 5(1):4560, 2014. ISSN 2041-1723. doi: 10.1038/ncomms5560. Publisher: Nature Publishing Group.

96. Gabriella R Sterne, Hideo Otsuna, Barry J Dickson, and Kristin Scott. Classification and genetic targeting of cell types in the primary taste and premotor center of the adult drosophila brain. eLife, 10:e71679, 2021. ISSN 2050-084X. doi: 10.7554/eLife.71679. Publisher: eLife Sciences Publications, Ltd.

97. Yiming Chen, Yen-Chu Lin, Tzu-Wei Kuo, and Zachary A. Knight. Sensory detection of food rapidly modulates arcuate feeding circuits. Cell, 160(5):829–841, 2015. ISSN 0092-8674, 1097-4172. doi: 10.1016/j.cell.2015.01.033. Publisher: Elsevier.

98. Alastair S. Garfield, Bhavik P. Shah, Christian R. Burgess, Monica M. Li, Chia Li, Jennifer S. Steger, Joseph C. Madara, John N. Campbell, Daniel Kroeger, Thomas E. Scammell, Bakhos A. Tannous, Martin G. Myers, Mark L. Andermann, Michael J. Krashes, and Bradford B. Lowell. Dynamic GABAergic afferent modulation of AgRP neurons. Nature Neuroscience, 19(12):1628–1635, 2016. ISSN 1546-1726. doi: 10.1038/nn.4392. Publisher: Nature Publishing Group.

99. Truong Ly, Jun Y. Oh, Nilla Sivakumar, Sarah Shehata, Naymalis La Santa Medina, Heidi Huang, Zhengya Liu, Wendy Fang, Chris Barnes, Naz Dundar, Brooke C. Jarvie, Anagh Ravi, Olivia K. Barnhill, Chelsea Li, Grace R. Lee, Jaewon Choi, Heeun Jang, and Zachary A. Knight. Sequential appetite suppression by oral and visceral feedback to the brainstem. Nature, 624(7990):130–137, 2023. ISSN 1476-4687. doi: 10.1038/s41586-023-06758-2. Publisher: Nature Publishing Group.

100. Claus Brandt, Hendrik Nolte, Sinika Henschke, Linda Engström Ruud, Motoharu Awazawa, Donald A. Morgan, Paula Gabel, Hans-Georg Sprenger, Martin E. Hess, Stefan Günther, Thomas Langer, Kamal Rahmouni, Henning Fenselau, Marcus Krüger, and Jens C. Brüning. Food perception primes hepatic ER homeostasis via melanocortin-dependent control of mTOR activation. Cell, 175(5):1321–1335.e20, 2018. ISSN 0092-8674. doi: 10.1016/j.cell.2018.10.015.

101. Hiroshi Tsuneki, Masanori Sugiyama, Toshihiro Ito, Kiyofumi Sato, Hiroki Matsuda, Kengo Onishi, Koharu Yubune, Yukina Matsuoka, Sanaka Nagai, Towa Yamagishi, Takahiro Maeda, Kosuke Honda, Akira Okekawa, Shiro Watanabe, Keisuke Yaku, Daisuke Okuzaki, Ryota Otsubo, Masanori Nomoto, Kaoru Inokuchi, Takashi Nakagawa, Tsutomu Wada, Teruhito Yasui, and Toshiyasu Sasaoka. Food odor perception promotes systemic lipid utilization. Nature Metabolism, 4(11):1514–1531, 2022. ISSN 2522-5812. doi: 10.1038/s42255-022-00673-y. Publisher: Nature Publishing Group.

102. Luke K. Burke, Tamana Darwish, Althea R. Cavanaugh, Sam Virtue, Emma Roth, Joanna Morro, Shun-Mei Liu, Jing Xia, Jeffrey W. Dalley, Keith Burling, Streamson Chua, Toni Vidal-Puig, Gary J. Schwartz, and Clémence Blouet. mTORC1 in AGRP neurons integrates exteroceptive and interoceptive food-related cues in the modulation of adaptive energy expenditure in mice. eLife, 6:e22848, 2017. ISSN 2050-084X. doi: 10.7554/eLife.22848.

103. Meet Zandawala and Jayati Gera. Leptin- and cytokine-like unpaired signaling in drosophila. Molecular and Cellular Endocrinology, 584:112165, 2024. ISSN 1872-8057. doi: 10.1016/j.mce.2024.112165.

104. Yiming Chen and Zachary A. Knight. Making sense of the sensory regulation of hunger neurons. BioEssays, 38(4):316–324, 2016. ISSN 1521-1878. doi: 10.1002/bies.201500167. _eprint: https://onlinelibrary.wiley.com/doi/pdf/10.1002/bies.201500167.

105. Leland C. Sudlow, Robert S. Edgecomb, and Larry L. Murdock. Regulation of labellar and tarsal taste thresholds in the black blowfly, phormia regina. Journal of Experimental Biology, 130(1):219–234, 1987. ISSN 0022-0949. doi: 10.1242/jeb.130.1.219.

106. Verónica María Corrales-Carvajal, Aldo A Faisal, and Carlos Ribeiro. Internal states drive nutrient homeostasis by modulating exploration-exploitation trade-off. eLife, 5:e19920, 2016. ISSN 2050-084X. doi: 10.7554/eLife.19920. Publisher: eLife Sciences Publications, Ltd.

107. Dennis Goldschmidt, Ibrahim Tastekin, Daniel Münch, Jin-Yong Park, Hannah Haberkern, Lúcia Serra, Célia Baltazar, Vivek Jayaraman, Gerald M. Rubin, and Carlos Ribeiro. A neuronal substrate for translating nutrient state and resource density estimations into foraging decisions. bioRxiv, 2023. doi: 10.1101/2023.07.19.549514. Pages: 2023.07.19.549514 Section: New Results.

108. Julie H. Simpson. Descending control of motor sequences in drosophila. Current Opinion in Neurobiology, 84:102822, 2024. ISSN 0959-4388. doi: 10.1016/j.conb.2023.102822.

109. Aya Yanagawa, Alexandra M. A. Guigue, and Frédéric Marion-Poll. Hygienic grooming is induced by contact chemicals in drosophila melanogaster. Frontiers in Behavioral Neuroscience, 8, 2014. ISSN 1662-5153. doi: 10.3389/fnbeh.2014.00254. Publisher: Frontiers.

110. Claire E. McKellar, Joshua L. Lillvis, Daniel E. Bath, James E. Fitzgerald, John G. Cannon, Julie H. Simpson, and Barry J. Dickson. Threshold-based ordering of sequential actions during drosophila courtship. Current biology: CB, 29(3):426–434.e6, 2019. ISSN 1879-0445. doi: 10.1016/j.cub.2018.12.019.

111. Paul Smolen, Douglas A. Baxter, and John H. Byrne. Modeling circadian oscillations with interlocking positive and negative feedback loops. Journal of Neuroscience, 21(17):6644–6656, 2001. ISSN 0270-6474, 1529-2401. doi: 10.1523/JNEUROSCI.21-17-06644.2001. Publisher: Society for Neuroscience Section: ARTICLE.

112. Philip K. Shiu, Gabriella R. Sterne, Nico Spiller, Romain Franconville, Andrea Sandoval, Joie Zhou, Neha Simha, Chan Hyuk Kang, Seongbong Yu, Jinseop S. Kim, Sven Dorkenwald, Arie Matsliah, Philipp Schlegel, Szi-chieh Yu, Claire E. McKellar, Amy Sterling, Marta Costa, Katharina Eichler, Alexander Shakeel Bates, Nils Eckstein, Jan Funke, Gregory S. X. E. Jefferis, Mala Murthy, Salil S. Bidaye, Stefanie Hampel, Andrew M. Seeds, and Kristin Scott. A drosophila computational brain model reveals sensorimotor processing. Nature, 634(8032):210–219, 2024. ISSN 1476-4687. doi: 10.1038/s41586-024-07763-9. Publisher: Nature Publishing Group.

113. Nikhil Sharma, Kali Flaherty, Karina Lezgiyeva, Daniel E. Wagner, Allon M. Klein, and David D. Ginty. The emergence of transcriptional identity in somatosensory neurons. Nature, 577(7790):392–398, 2020. ISSN 1476-4687. doi: 10.1038/s41586-019-1900-1.

114. Chen Ran, Jack C. Boettcher, Judith A. Kaye, Catherine E. Gallori, and Stephen D. Liberles. A brainstem map for visceral sensations. Nature, 609(7926):320–326, 2022. ISSN 1476-4687. doi: 10.1038/s41586-022-05139-5. Publisher: Nature Publishing Group.

115. Kazuhiro Nakamura and Yoshiko Nakamura. Hunger and satiety signaling: Modeling two hypothalamomedullary pathways for energy homeostasis. BioEssays: News and Reviews in Molecular, Cellular and Developmental Biology, 40(8):e1700252, 2018. ISSN 1521-1878. doi: 10.1002/bies.201700252.

116. Danielle S. Lafferty, Jeremiah Isaac, Joelyz S. Wolcott, Amy Phan, Lily Reck, and Andrew Lutas. An amygdalopontine pathway promotes motor programs of ingestion. bioRxiv, 2025. doi: 10.1101/2025.06.05.657686. Pages: 2025.06.05.657686 Section: New Results.

117. H. J. Grill and R. Norgren. The taste reactivity test. II. mimetic responses to gustatory stimuli in chronic thalamic and chronic decerebrate rats. Brain Research, 143(2):281–297, 1978. ISSN 0006-8993. doi: 10.1016/0006-8993(78)90569-3.

118. J. B. Travers. Motor control of feeding and drinking. In Larry R. Squire, editor, Encyclopedia of Neuroscience, pages 1001–1007. Academic Press, 2009. ISBN 978-0-08-045046-9. doi: 10.1016/B978-008045046-9.00449-6.

119. Zepeng Yao and Kristin Scott. Serotonergic neurons translate taste detection into internal nutrient regulation. Neuron, 110(6):1036–1050.e7, 2022. ISSN 08966273. doi: 10.1016/j.neuron.2021.12.028.

120. Tino Just, Hans Wilhelm Pau, Ulrike Engel, and Thomas Hummel. Cephalic phase insulin release in healthy humans after taste stimulation? Appetite, 51(3):622–627, 2008. ISSN 0195-6663. doi: 10.1016/j.appet.2008.04.271.

121. Michael L. Power and Jay Schulkin. Anticipatory physiological regulation in feeding biology: Cephalic phase responses. Appetite, 50(2):194–206, 2008. ISSN 0195-6663. doi: 10.1016/j.appet.2007.10.006.

