## Supplementary Figures for "From Sensory Detection to Motor Action: The Comprehensive *Drosophila* Taste-Feeding Connectome"

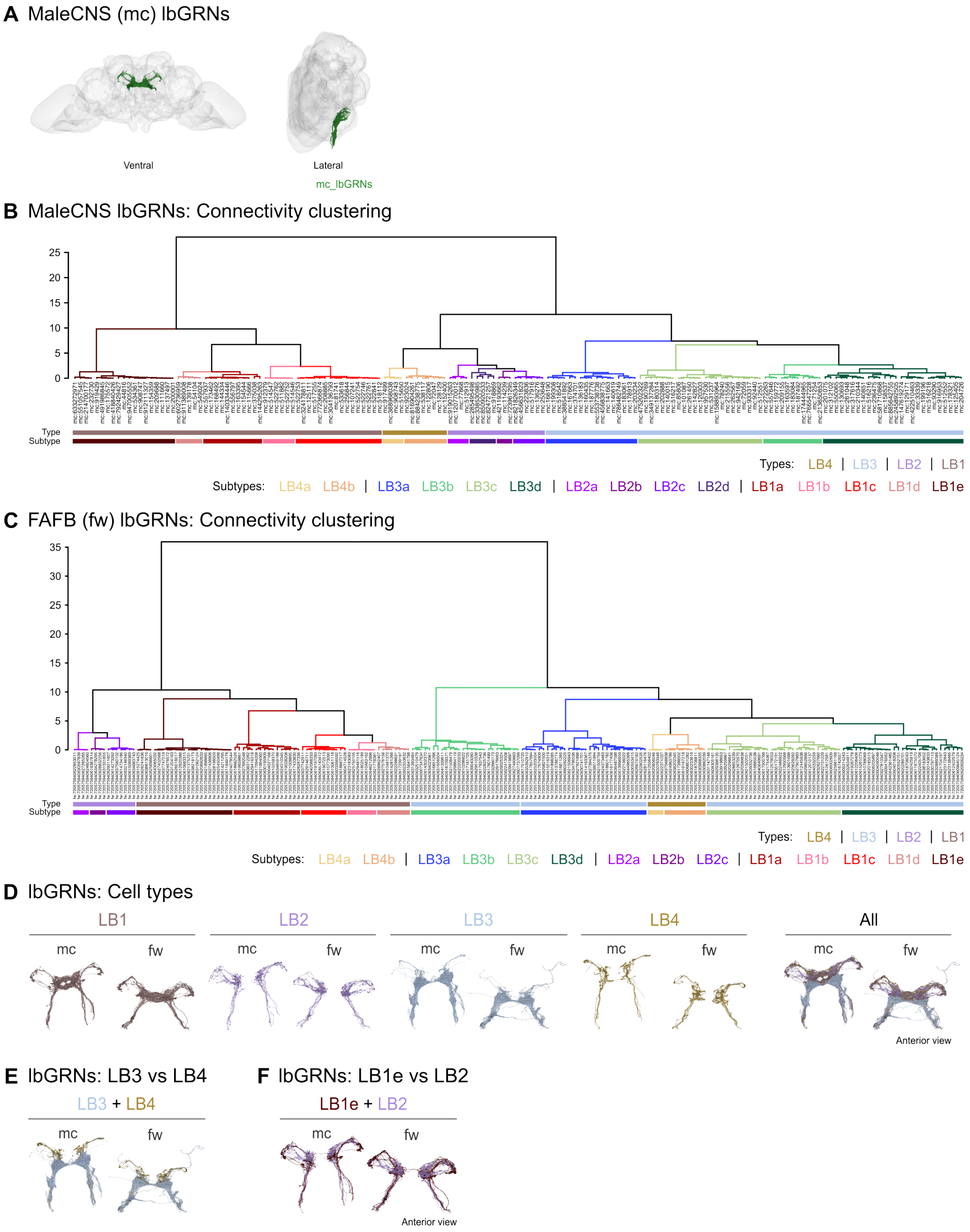


**Figure S1: Cell typing the labellar bristle GRNs.** **A.** Rendering of lbGRNs in the male CNS in two different views, ventral and lateral. **B.** Hierarchical clustering based on outputs of the lbGRNs identified in the male CNS using cosine similarity and a defined threshold of 2 synapses. The neurons were divided into four types, LB1-4, which were, in turn, subdivided into the following subtypes: LB1a-e, LB2a-d, LB3a-d, and LB4a-b. **C.** Connectivity clustering of the FAFB – FlyWire lbGRNs performed using the same conditions as in B. The same four lbGRN types were defined (top colored bars), LB1-4, each, further subdivided into the corresponding subtypes (bottom colored bars): LB1a-e, LB2a-c, LB3a-d, and LB4a-b. **D.** Left: anterior view of renderings of each lbGRN type across the male CNS and FAFB – Flywire. Right: anterior view of the individual lbGRN type renderings overlaid for each connectome. **E.** Anterior view of the renderings of the LB3 and LB4 types in both connectomes to illustrate their different projection fields. **F.** Anterior view of the renderings of the LB1e subtype and the LB2 type overlaid in both connectomes to demonstrate their close morphologies, despite distinct direct downstream connectivity.

**A** Male CNS (mc) tpGRNs

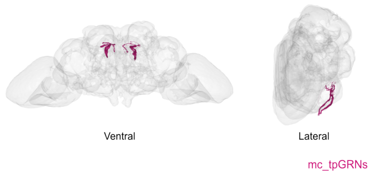


**B** Male CNS tpGRNs: Connectivity clustering

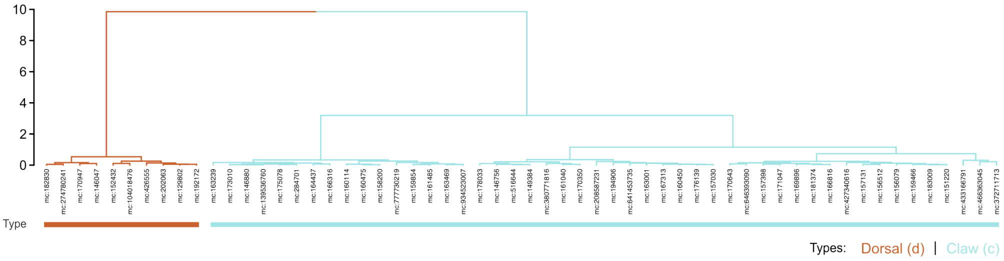


**Figure S2: Cell typing the taste peg GRNs. A.** Rendering of tpGRNs in the male CNS in two different views, ventral and lateral, to reveal their SEZ arborizations. **B.** Hierarchical clustering based on outputs of the tpGRNs identified in the male CNS using cosine similarity and a defined threshold of 2 synapses, reveals two different cell types (colored accordingly), which we named dorsal and claw tpGRNs.

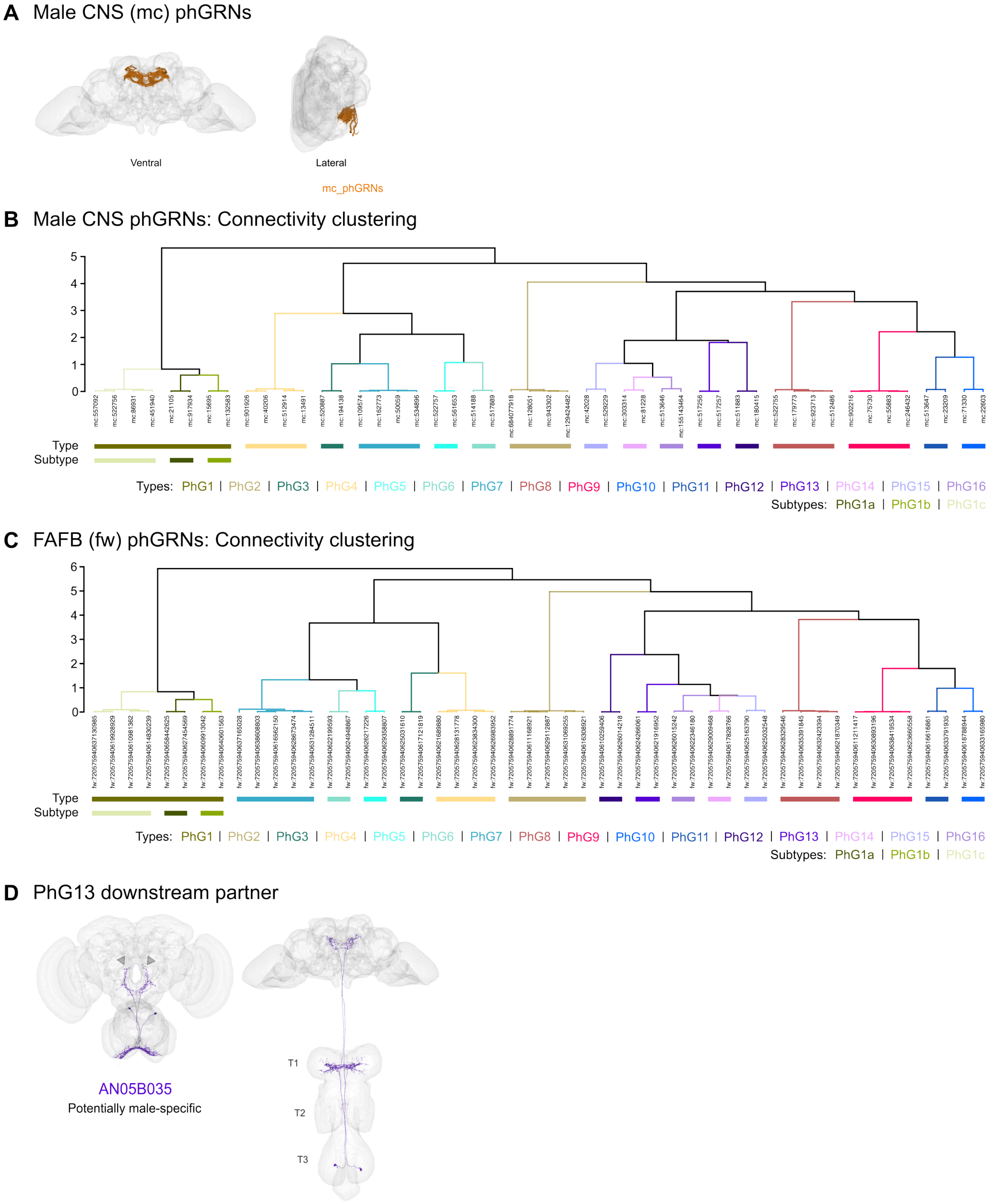


**Figure S3: Cell typing the pharyngeal GRNs.** **A.** Rendering of phGRNs in the male CNS in two different views, ventral and lateral, to reveal their SEZ projection pattern. **B.** Hierarchical clustering based on outputs of the phGRNs identified in male CNS using cosine similarity and a defined threshold of 5 synapses. The sixteen phGRN cell types are rendered in different colors. **C.** Connectivity clustering of the FAFB – FlyWire phGRNs performed using the same conditions as in B., which resulted in the PhG1-16 types, , and PhG1a-c subtypes. **D.** Rendering of AN05B035, a potentially male-specific AN, which is one of the top downstream partners of the PhG13 type. The AN is shown in two distinct views: anterior (left) to highlight the AN's SEZ projections, especially the branches that innervate the known sexually dimorphic region around the esophagus, and ventral (right) to reveal its arborization in the T1 segment of the VNC.

**A** Male CNS (mc) lgAGRNs

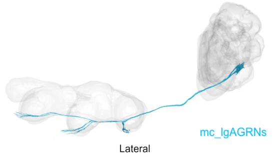


**B** Male CNS lgAGRNs: Connectivity clustering

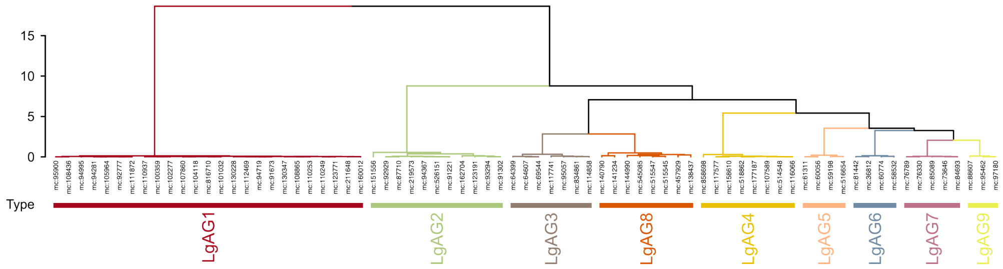


**C** Male CNS lgAGRNs: Counts

| | LgAG1 | | LgAG2 | | LgAG3 | | LgAG4 | | LgAG5 | | LgAG6 | | LgAG7 | | LgAG8 | | LgAG9 | |
|---|---|---|---|---|---|---|---|---|---|---|---|---|---|---|---|---|---|---|
| | R | L | R | L | R | L | R | L | R | L | R | L | R | L | R | L | R | L |
| Frontleg | 5 | 5 | 2 | 2 | 0 | 0 | 0 | 0 | 2 | 2 | 2 | 2 | 2 | 2 | 0 | 0 | 2 | 1 |
| Midleg | 4 | 3 | 2 | 2 | 1 | 2 | 2 | 1 | 0 | 0 | 0 | 0 | 1 | 0 | 2 | 1 | 0 | 0 |
| Hindleg | 4 | 4 | 2 | 1 | 2 | 2 | 2 | 3 | 0 | 0 | 0 | 0 | 0 | 0 | 2 | 3 | 0 | 0 |

**D** FAFB (fw) lgAGRNs: Connectivity clustering

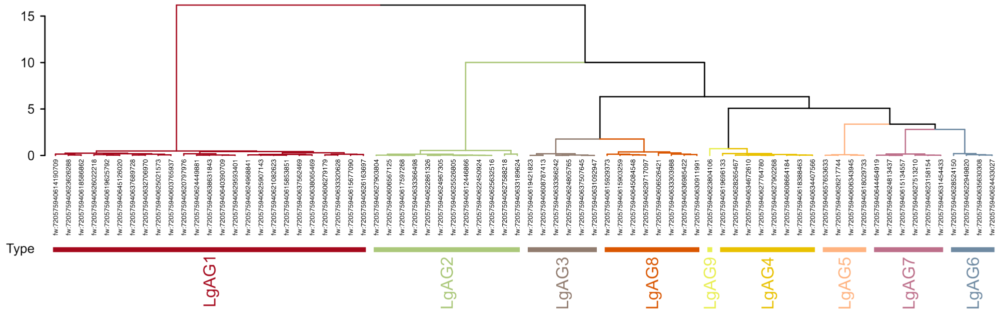


**Figure S4: Cell typing the leg bristle ascending GRNs. A.** Lateral view of the lgAGRN renderings in the male CNS. **B.** Hierarchical clustering based on outputs of the lgAGRNs identified in the male CNS using cosine similarity and a defined threshold of 2 synapses. The nine lgAGRN cell types are rendered in different colors. **C.** Counts per side of each male LgAG type across the fore-, mid-, and hindlegs, which reveals that the types can be grouped into three categories: one including the types that span all legs, a second corresponding to mid- and hindleg exclusive types, and a third including most foreleg types. **D.** Connectivity clustering of the FAFB – FlyWire lgAGRNs performed using the same conditions as in B., resulting in the corresponding LgAG1-9 types.

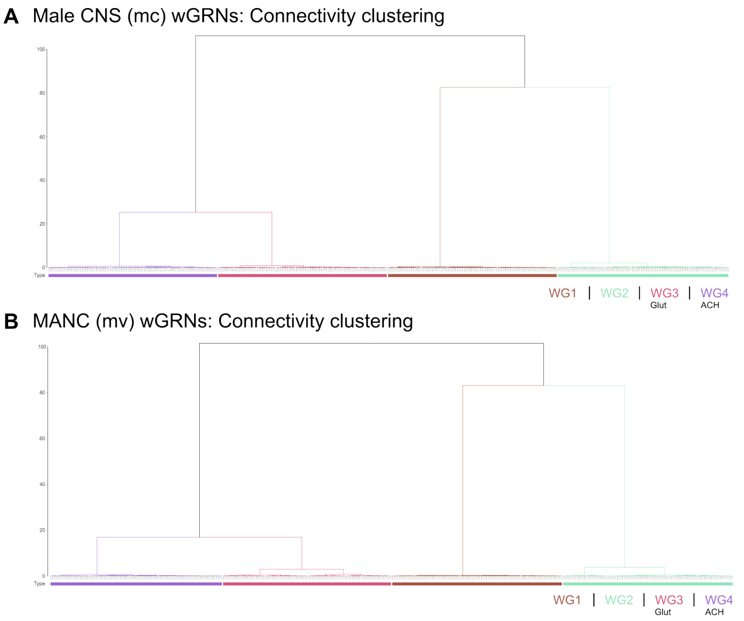


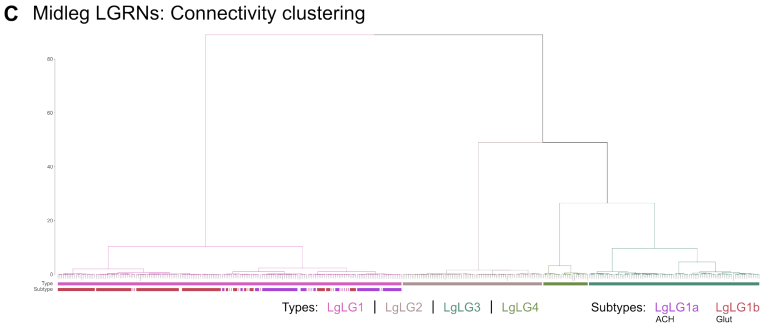


**D** Midleg LGRNs: Counts

| | | R | L |
|---|---|---|---|
| LgLG1 (a/b) | mc | 52 (25/26) | 48 (23/24) |
| | mv | 44 (12/32) | 45 (8/37) |
| LgLG2 | mc | 22 | 20 |
| | mv | 21 | 14 |
| LgLG3 | mc | 26 | 23 |
| | mv | 24 | 21 |
| LgLG4 | mc | 5 | 9 |
| | mv | 6 | 5 |

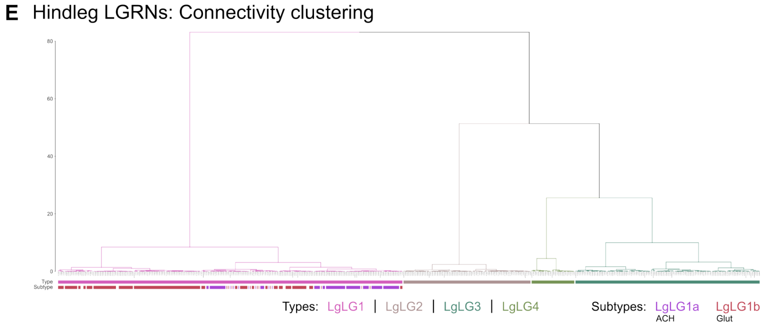


**F** Hindleg LGRNs: Counts

| | | R | L |
|---|---|---|---|
| LgLG1 (a/b) | mc | 51 (25/26) | 53 (25/24) |
| | mv | 52 (12/40) | 49 (8/41) |
| LgLG2 | mc | 20 | 21 |
| | mv | 21 | 14 |
| LgLG3 | mc | 27 | 28 |
| | mv | 29 | 26 |
| LgLG4 | mc | 8 | 6 |
| | mv | 6 | 6 |

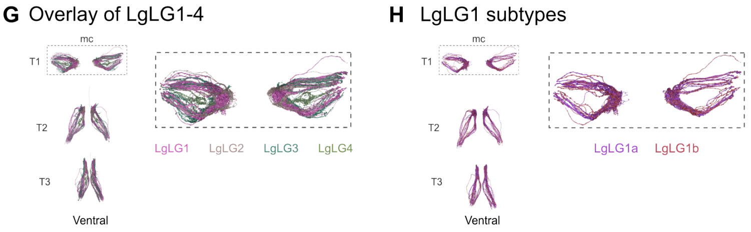


**Figure S5: Cell typing the wing and leg bristle local GRNs. A.** Hierarchical clustering of the wGRNs identified in the male CNS based on their outputs. The clustering was done using cosine similarity and a defined threshold of 2 synapses. The identified four wGRN types are rendered in different colors. The predicted neurotransmitter released by WG3 and WG4 GRNs, respectively, glutamate (Glut) and acetylcholine (ACH), is also indicated. **B.** Connectivity clustering of the MANC wGRNs using the same parameters as described in A. The corresponding four types are shown. **C.** Co-clustering of the male CNS and MANC midleg LGRNs performed using the conditions described in A.. Colored bars on top indicate the type, LgLG1-4, and the bottom ones the subtypes, LgLG1a-b. The latter were defined according to the predicted neurotransmitter release by each LgLG1 GRN, either ACH or Glut. **D.** Counts per side for each midleg LGRN type and subtype, when applicable, across male CNS and MANC. **E.** Co-clustering of the male CNS and MANC hindleg LGRNs as done for the midleg LGRNs, resulting in the corresponding types, LgLG1-4, and subtypes, LgLG1a-b. **F.** Counts per side for each hindleg LGRN type and subtype, when applicable, in both connectomes. **G.** Ventral view of the overlay of the renderings of the male CNS LgLG1-4. **H.** Ventral view of the overlay of the renderings of the male CNS LgLG1 subtypes, LgLG1a-b, revealing overlapping projection fields.

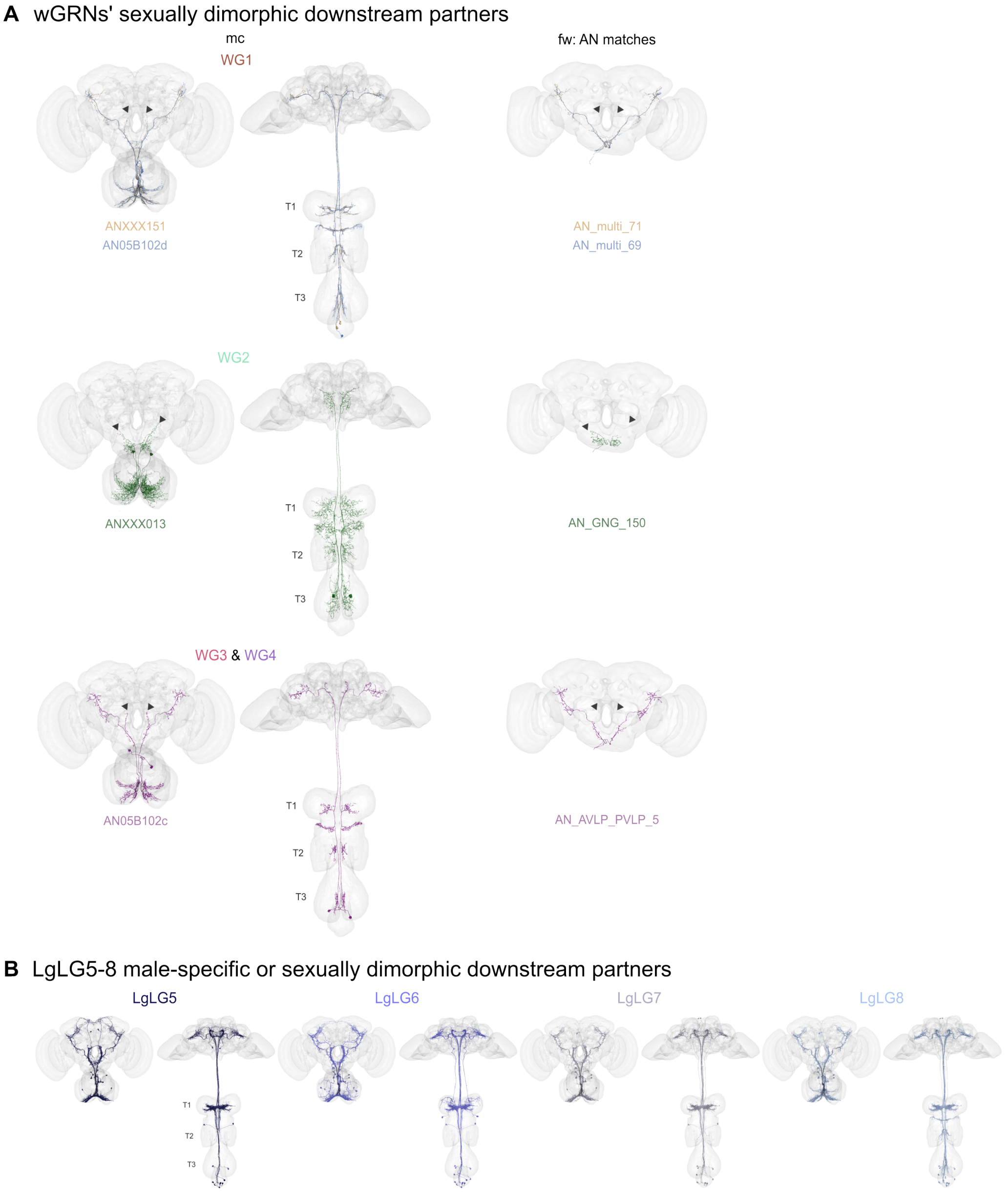


**Figure S6: Sexually dimorphic and male-specific downstream partners of VNC-projecting GRNs. A.** Renderings of sexually dimorphic partners of the wGRN types in the male CNS (left column, anterior and ventral views) and FAFB – FlyWire (right column, anterior view). The male CNS and FAFB – FlyWire cell types are indicated. Sexually dimorphic projections are highlighted with arrowheads. **B.** Male-specific or sexually dimorphic downstream partners of LgLG5-8 (left: anterior views, right: ventral views).

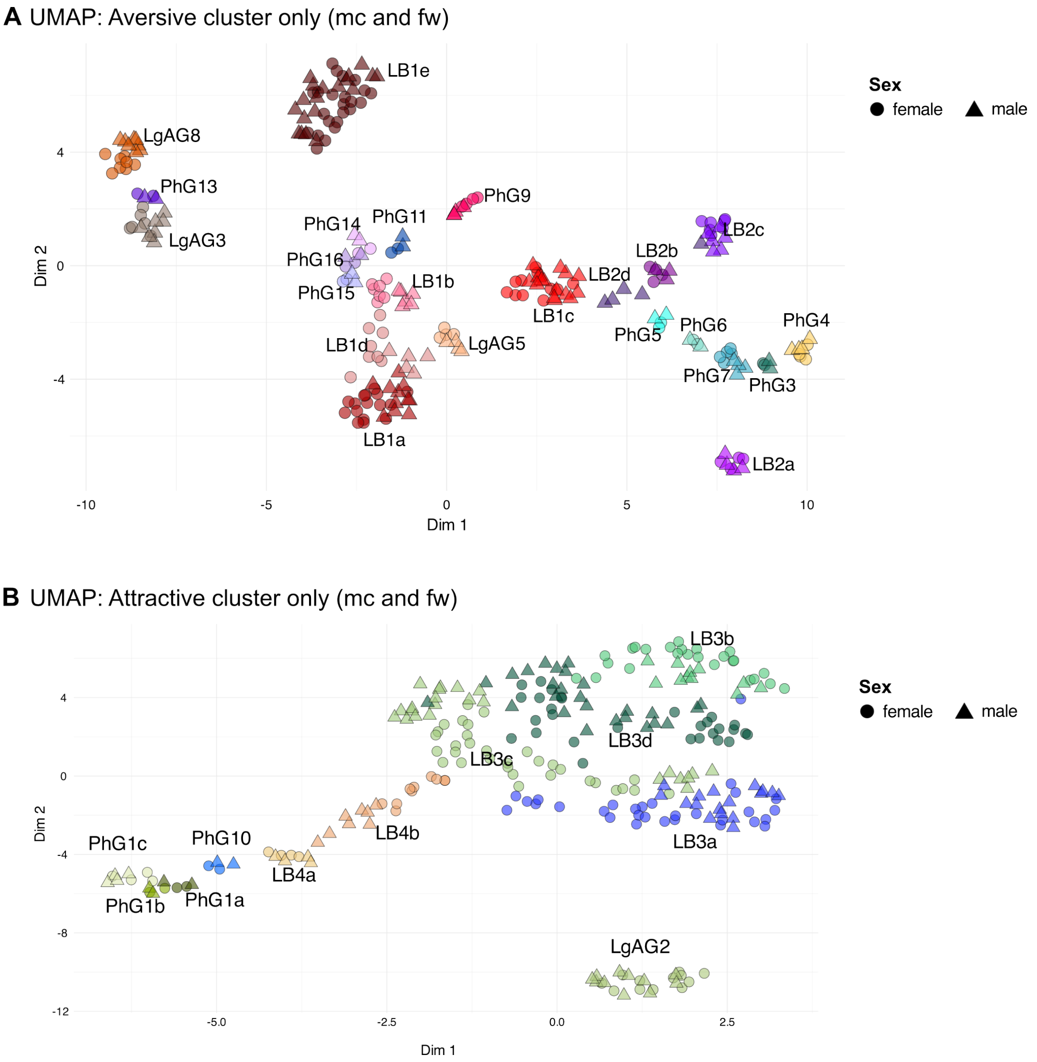


**Figure S7: UMAP embedding of the appetitive and aversive clusters. A**. and **B.** UMAP embedding of only **A.** the aversive or **B.** the attractive GRN types in the male CNS and FAFB – FlyWire. Each GRN is colored by its type. The shapes represent the sex of the animal. Circle: female (FAFB – FlyWire), triangle: male (male CNS).

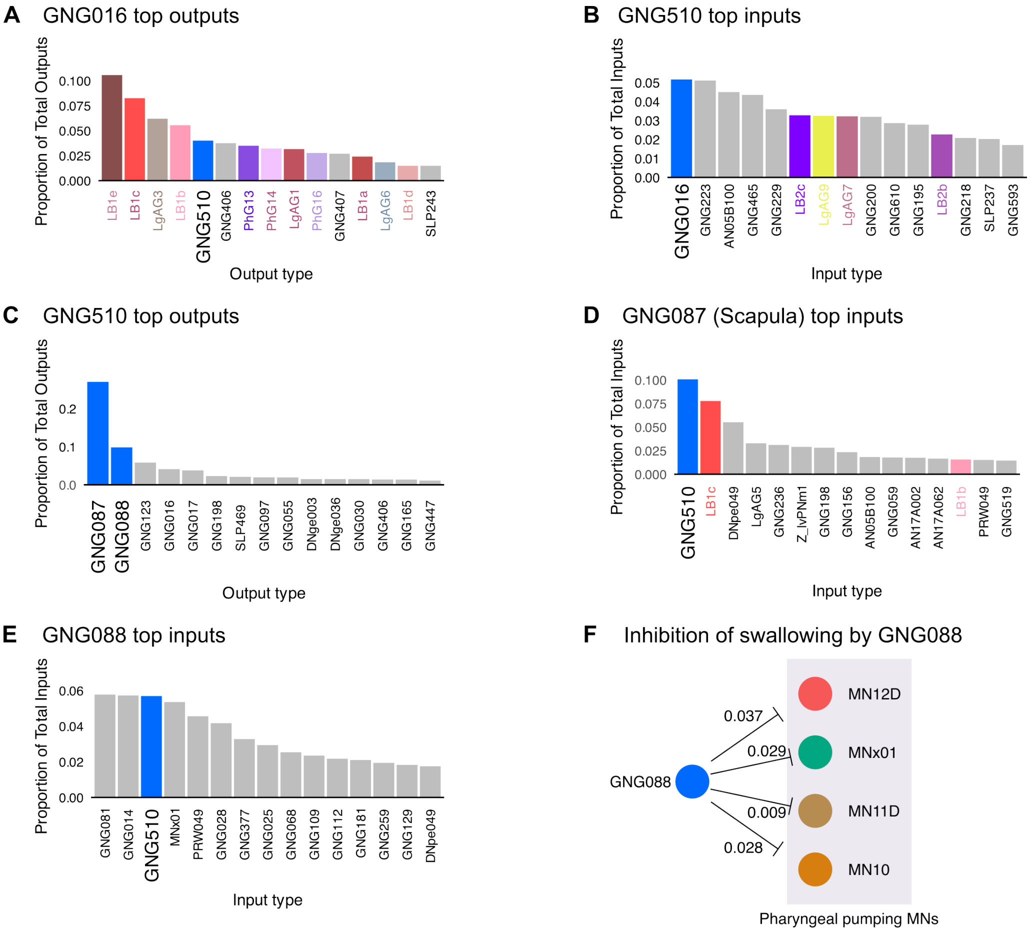


**Figure S8: Connectivity of key aversive downstream partners. A.** Top output partners of GNG016 shown as proportions of its total outputs. GRN types are marked by their respective colors. GNG510 is highlighted as the top downstream interneuron. **B.** The top input partners of GNG510 are shown as the proportion of its total inputs. GNG016 is highlighted as the top upstream partner. GRN types are marked by their respective colors. **C.** The top output partners of GNG510 are shown as the proportions of its total outputs. GNG087 and GNG088 are highlighted as the top downstream partners. **D.** Top input partners of GNG087 (Scapula) are shown as the proportion of its total inputs. GNG510 is highlighted as the top upstream partner. GRN types are marked by their respective colors. **E.** The top input partners of GNG088 are shown as the proportion of its total inputs. GNG510 is highlighted as the top upstream partner. **F.** GNG088 is a premotor neuron that is predicted to inhibit pharyngeal pumping motor neurons. Synaptic weights are given as the proportion of total inputs on the downstream MNs.

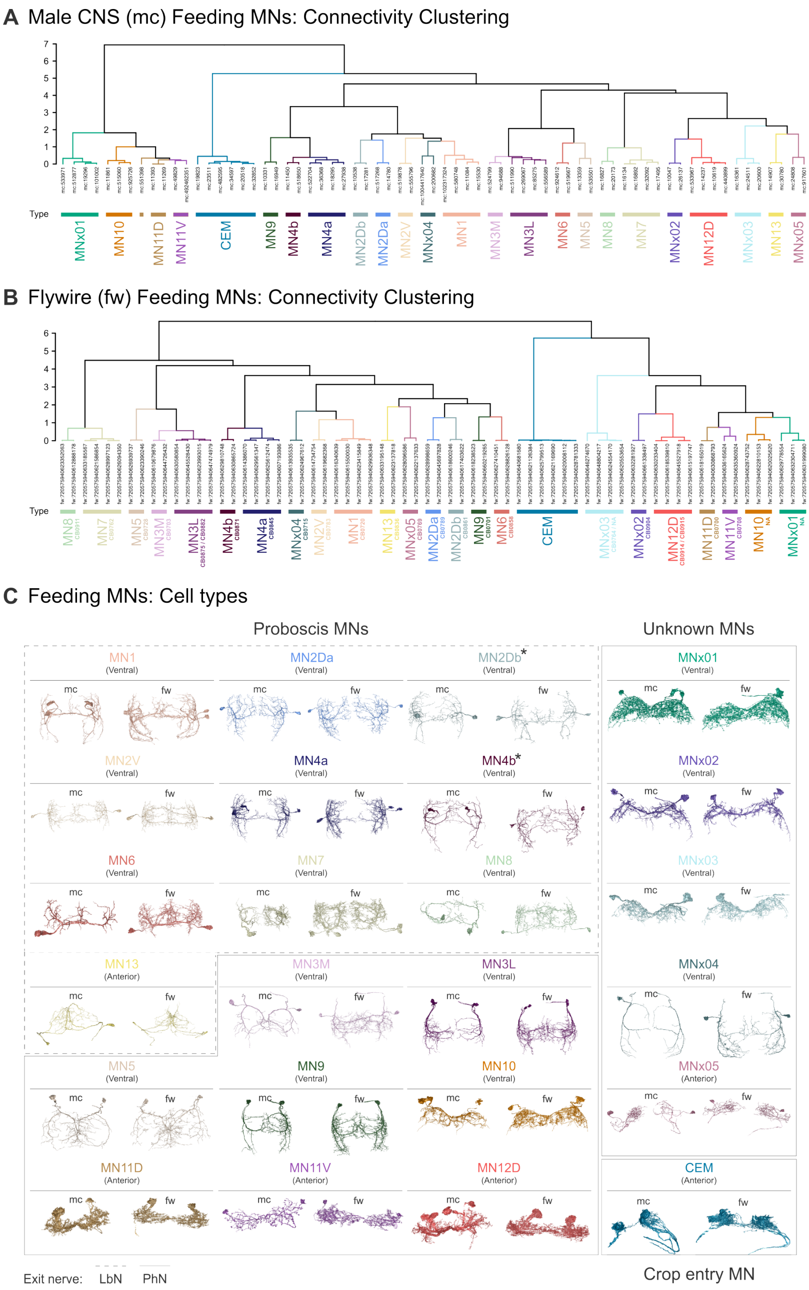


**D** Feeding MNs: Counts

| | | R | L |
|---|---|---|---|
| MN1 | mc | 2 | 2 |
| | fw | 2 | 2 |
| MN2Da | mc | 1 | 1 |
| | fw | 1 | 1 |
| MN2Db | mc | 1 | 1 |
| | fw | 1 | 1 |
| MN2V | mc | 1 | 1 |
| | fw | 1 | 1 |
| MN3M | mc | 1 | 1 |
| | fw | 1 | 1 |
| MN3L | mc | 2 | 2 |
| | fw | 2 | 2 |
| MN4a | mc | 2 | 2 |
| | fw | 2 | 2 |
| MN4b | mc | 1 | 1 |
| | fw | 1 | 1 |
| MN5 | mc | 1 | 1 |
| | fw | 1 | 1 |
| MN6 | mc | 1 | 1 |
| | fw | 1 | 1 |
| MN7 | mc | 2 | 2 |
| | fw | 2 | 2 |
| MN8 | mc | 1 | 1 |
| | fw | 1 | 1 |
| MN9 | mc | 1 | 1 |
| | fw | 1 | 1 |
| MN10 | mc | 2 | 1 |
| | fw | 2 | 1 |
| MN11D | mc | 2 | 1 |
| | fw | 1 | 1 |
| MN11V | mc | 1 | 1 |
| | fw | 1 | 1 |
| MN12D | mc | 2 | 2 |
| | fw | 2 | 2 |
| MN13 | mc | 1 | 1 |
| | fw | 1 | 1 |
| CEM | mc | 2 | 4 |
| | fw | 3 | 3 |
| MNx01 | mc | 1 | 3 |
| | fw | 3 | 0 |
| MNx02 | mc | 1 | 1 |
| | fw | 1 | 1 |
| MNx03 | mc | 1 | 2 |
| | fw | 1 | 3 |
| MNx04 | mc | 1 | 1 |
| | fw | 1 | 1 |
| MNx05 | mc | 1 | 1 |
| | fw | 1 | 1 |

**Figure S9: Cell typing of the feeding motor neurons. A.** Hierarchical clustering of the feeding motor neurons (MNs) identified in the male CNS based on their inputs. The clustering was done using cosine similarity and a defined threshold of 5 synapses. The defined MN types are rendered in different colors. **B.** Connectivity clustering of the FAFB – FlyWire feeding MNs using the same parameters as described above. The corresponding MN types are rendered in different colors and, when existing, the FAFB – FlyWire cell type is also indicated. **C.** Rendering of each feeding MN type in either a ventral or an anterior view across the male CNS and FAFB – FlyWire. The MNs' exiting nerve is indicated with a dotted or a solid line: the labial (LbN) or the pharyngeal (PhN) nerves. The MNs are divided into proboscis MNs, crop entry-controlling MNs, and the unknown ones, which, to our knowledge, have never been described in the literature. **D.** Counts per side for each feeding MN type in the two connectomes.

**Muscle**

| | | 1 | 2D | 2V | 3L | 3M | 4 | 5 | 6 | 7 | 8 | 9 | 10 | 11D | 11V | 12D |
|---|---|---|---|---|---|---|---|---|---|---|---|---|---|---|---|---|
| Positioning | Rostrum | + | + | + | - | - | - | ? | - | + | - | + | ? | - | - | - |
| | Haustellum | + | - | - | + | + | + | ? | + | + | - | + | ? | - | - | - |
| | Labellum | - | - | - | - | - | - | ? | + | + | - | - | ? | - | - | - |
| Ingestive | Labellar spread | - | - | - | - | - | - | ? | - | + | + | - | ? | - | - | - |
| | Pumping | - | - | - | - | - | - | +* | - | - | - | - | +* | + | + | + |

* Proposed function based on muscle insertion site

**Figure S10: Proboscis muscles and their functions based on the literature.** The muscles involved in proboscis positioning and ingestion are highlighted. Rostrum: rostrum movements, haustellum: haustellum movements, labellum: labellar movements except labellar abduction (spreading). Labellar spreading and pharyngeal pumping are grouped as ingestive processes. Functions are derived from[24,25]. Plus sign (+) indicates that the function as been experimentally validated. The minus sign (-) indicates that the muscle has not been shown to be involved in the movement. Asterisk (*) indicates a proposed function based on the insertion site of the muscle.

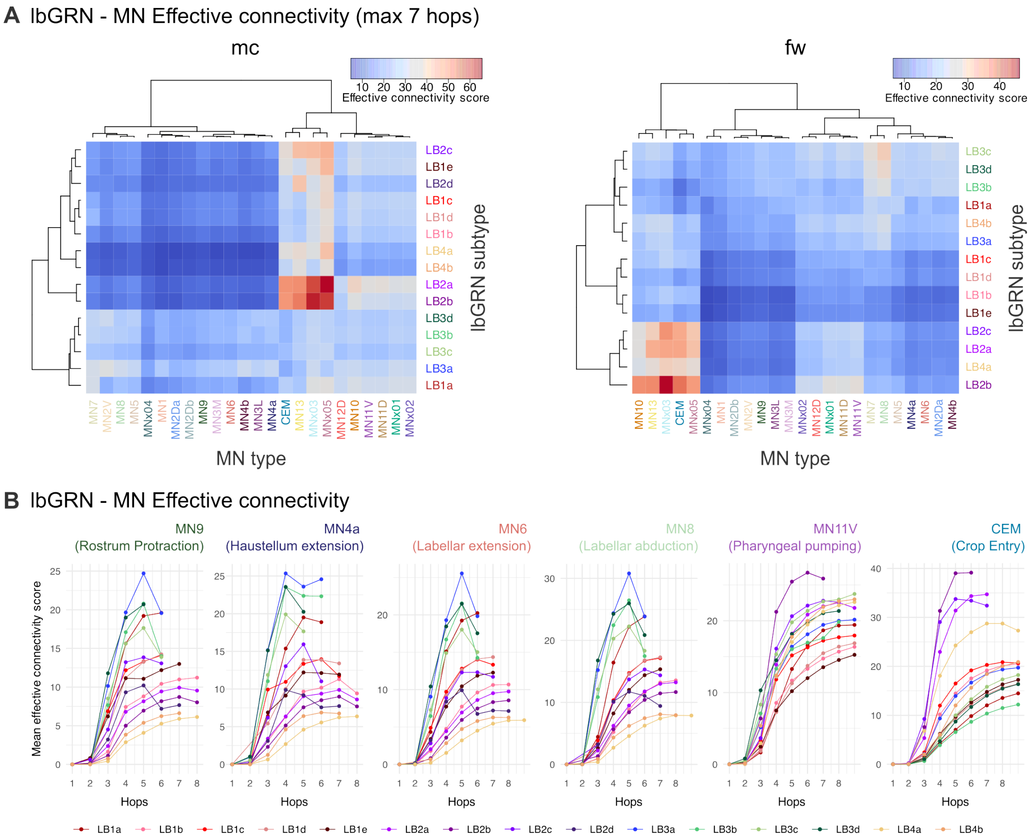


**Figure S11: Effective connectivity of labellar bristle GRNs to feeding motor neurons. A.** Heatmap for mean effective connectivity scores from lbGRNs to feeding MN types in the male CNS (left panel) and FAFB – FlyWire (right panel). A maximum of seven hops was used. Ward's method was used for hierarchical clustering and building the dendrograms. **B.** Effective connectivity of lbGRNs to MN types that control different steps of proboscis positioning, ingestion, and crop entry across different hops (maximum 10 hops, see Reachability analysis in Methods). Motor neurons and their roles are indicated.

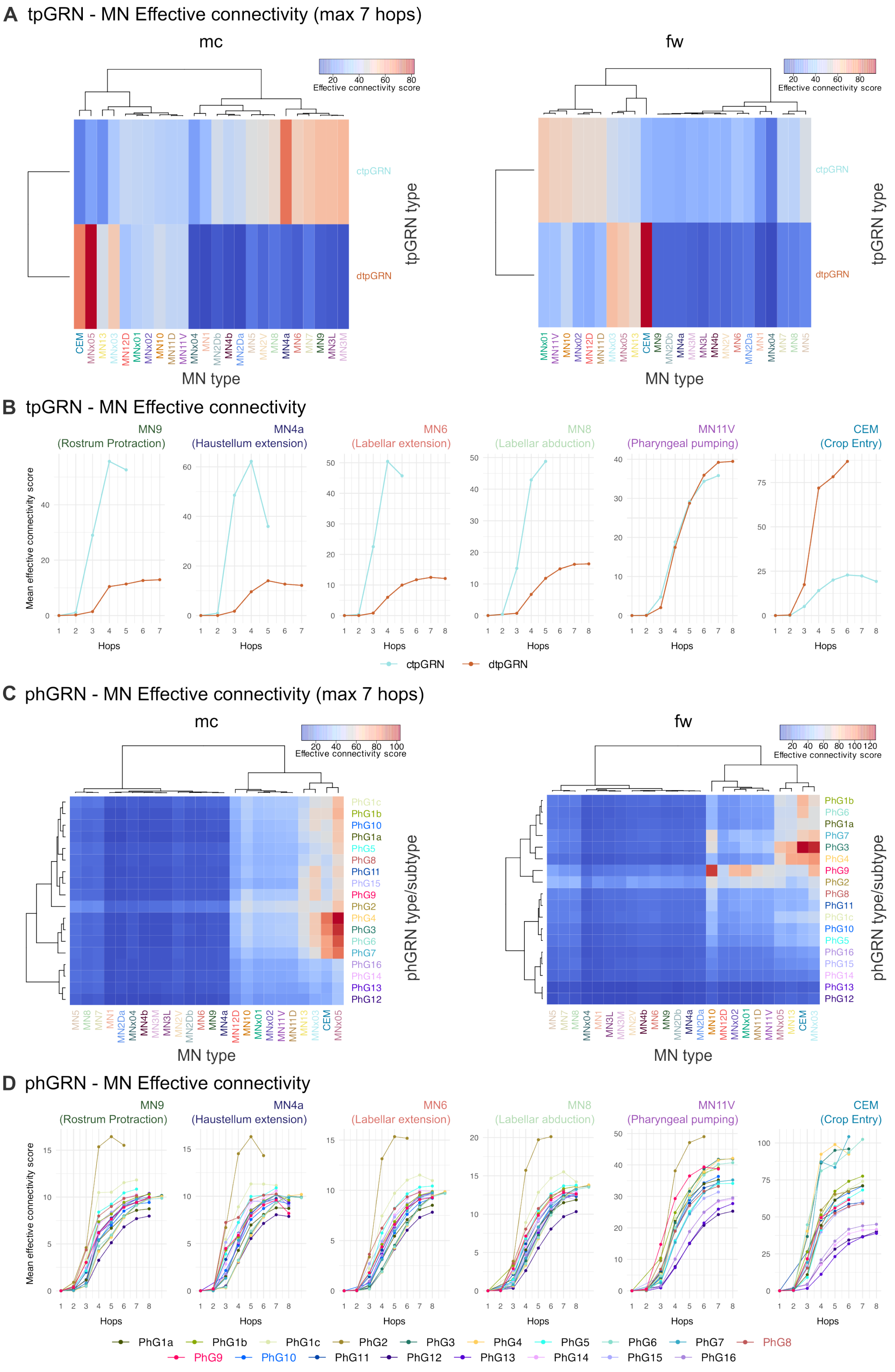


**Figure S12: Effective connectivity of taste peg and pharyngeal GRNs to feeding motor neurons. A.** Heatmap for mean effective connectivity scores from tpGRNs to feeding MN types in the male CNS (left panel) and FAFB – FlyWire (right panel). A maximum of seven hops was used. Ward's method was used for hierarchical clustering and building the dendrograms. **B.** Effective connectivity of tpGRNs to MN types that control different steps of proboscis positioning, ingestion, and crop entry across different hops (maximum 10 hops). Motor neurons and their roles are indicated. **C.** Heatmap for mean effective connectivity scores from phGRNs to feeding MN types in male CNS (left panel) and FAFB – FlyWire (right panel). A maximum of seven hops was used. Ward's method was used for hierarchical clustering and building the dendrograms. **D.** Effective connectivity of phGRNs to MN types that control different steps of proboscis positioning, ingestion, and crop entry across different hops (maximum 10 hops). Motor neurons and their roles are indicated.

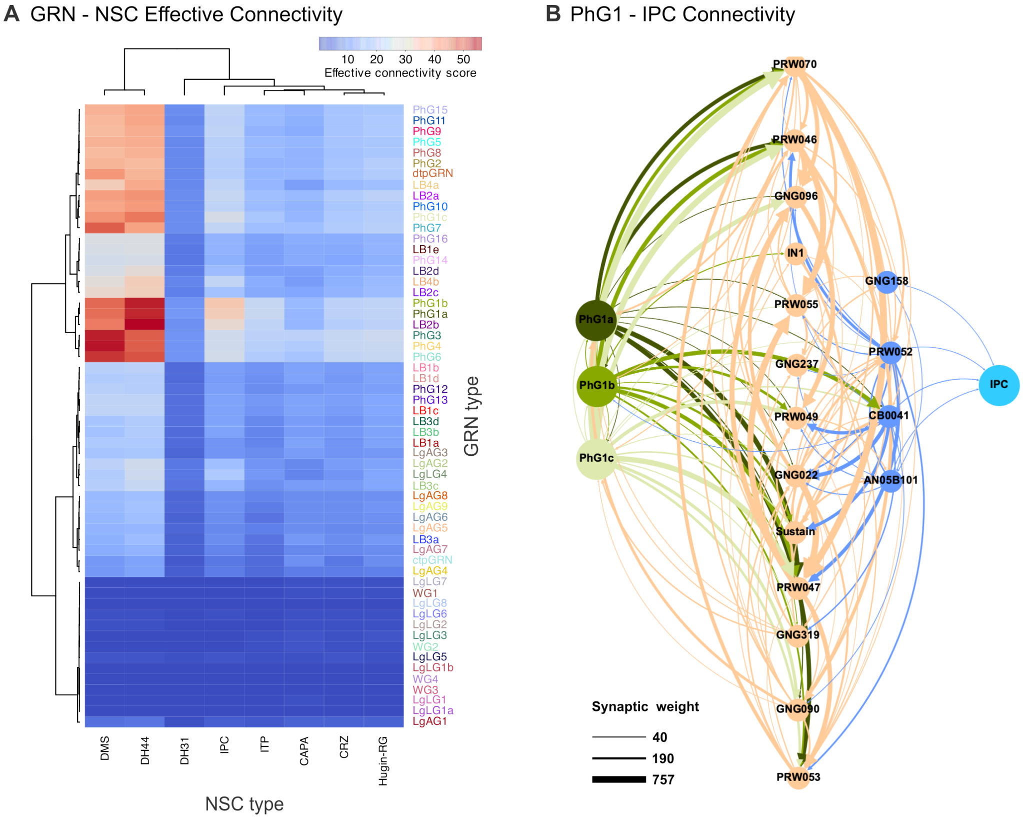


**Figure S13: Effective connectivity of GRN types with neurosecretory cells. A.** Effective connectivity between GRN types and neurosecretory cell types plotted as a heatmap. A maximum of seven hops was used. Ward's method was used for hierarchical clustering and building the dendrograms. **B.** The neural pathway between PhG1 subtypes and IPCs. Second-order neurons are colored in light orange. Pre-IPC neurons are colored in light blue. If a downstream cell type has been previously studied or a driver line exists, the name from the literature is used. Absolute synaptic weights are given in the legend.

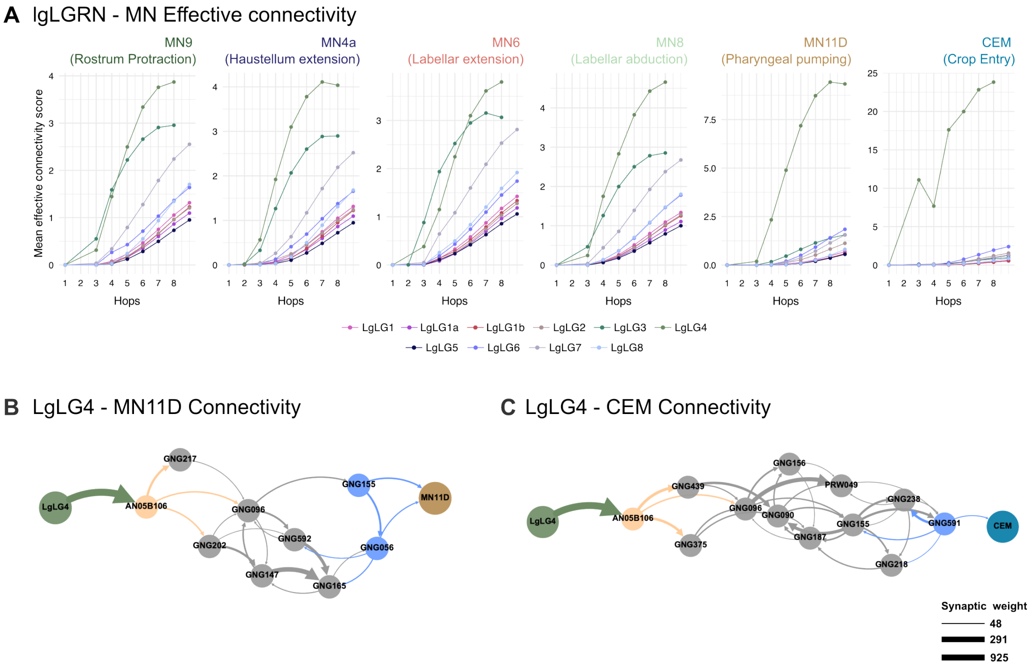


**Figure S14: Effective connectivity of leg bristle local GRNs to feeding motor neurons. A.** Effective connectivity of lgLGRNs to MN types that control different steps of proboscis positioning, ingestion, and crop entry across different hops (maximum 10 hops). Motor neurons and their roles are indicated. **B.** and **C.** The sensorimotor circuit that connects LgLG4 to **B.** pharyngeal pumping motor neuron MN11D and **C.** crop entry motor neuron CEM. The male CNS cell types are indicated. Absolute synaptic weights are given in the legend.

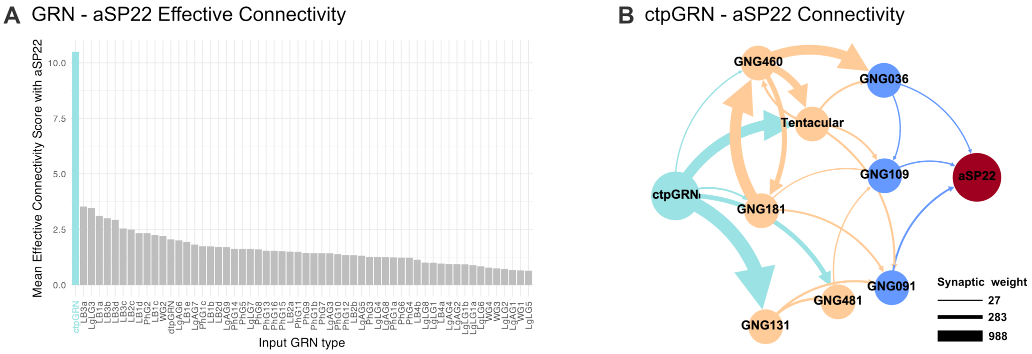


**Figure S15: Effective connectivity of ctpGRNs to aSP22. A.** Mean effective connectivity of each GRN type to courtship-promoting descending neuron aSP22. ctpGRN, which has the highest score, is highlighted. **B.** The neural pathway that connects ctpGRN to aSP22 in three hops. Second-order neurons are colored in light orange. Pre-DN neurons are colored in light blue. If a downstream cell type has been previously studied or a driver line exists, the name in the literature is used. Absolute synaptic weights are given in the legend.
